# Multimodal engineering of extracellular vesicles for efficient intracellular protein delivery

**DOI:** 10.1101/2023.04.30.535834

**Authors:** Xiuming Liang, Dhanu Gupta, Junhua Xie, Elien Van Wonterghem, Lien Van Hoecke, Justin Hean, Zheyu Niu, Oscar P. B. Wiklander, Wenyi Zheng, Rim Jawad Wiklander, Rui He, Doste R. Mamand, Jeremy Bost, Guannan Zhou, Houze Zhou, Samantha Roudi, Antje M. Zickler, André Görgens, Daniel W. Hagey, Olivier G. de Jong, Aileen Geobee Uy, Yuanyuan Zong, Imre Mäger, Carla Martin Perez, Thomas C. Roberts, Pieter Vader, Antonin de Fougerolles, Matthew J. A. Wood, Roosmarijn E. Vandenbroucke, Joel Z. Nordin, Samir EL Andaloussi

## Abstract

Extracellular vesicles (EVs) are promising tools to transfer macromolecular therapeutic molecules to recipient cells, however, efficient functional intracellular protein delivery by EVs remains challenging. Here, we have developed novel and versatile systems that leverage selected molecular tools to engineer EVs for robust cytosolic protein delivery both *in vitro* and *in vivo*. These systems, termed VSV-G plus EV-sorting Domain-Intein-Cargo (VEDIC) and VSV-G-Foldon-Intein-Cargo (VFIC), exploit an engineered mini-intein (intein) protein with self-cleavage activity to link cargo to an EV-sorting domain and release it from the EV membrane inside the EV lumen. In addition, we utilize the fusogenic protein VSV-G to facilitate endosomal escape and cargo release from the endosomal system to the cytosol of recipient cells. Importantly, we demonstrate that the combination of the self-cleavage intein, fusogenic protein and EV-sorting domain are indispensable for efficient functional intracellular delivery of cargo proteins by engineered EVs. As such, nearly 100% recombination and close to 80% genome editing efficiency in reporter cells were observed by EV-transferred Cre recombinase and Cas9/sgRNA RNPs, respectively. Moreover, EV-mediated Cre delivery by VEDIC or VFIC engineered EVs resulted in significant *in vivo* recombination in Cre-LoxP R26-LSL-tdTomato reporter mice following both local and systemic injections. Finally, we applied these systems for improved treatment of LPS-induced systemic inflammation by delivering a super-repressor of NF-ĸB activity. Altogether, this study describes a platform by which EVs can be utilized as a vehicle for the efficient intracellular delivery of macromolecular therapeutics for treatments of disease.

**Graphic summary: Development of VEDIC and VFIC systems for high-efficiency intracellular protein delivery in vitro and in vivo.:** Intein in tripartite fusion protein (EV-sorting Domain-Intein-Cargo) performs C-terminal cleavage during the process of EV-biogenesis, resulting in enriched free cargo proteins inside of vesicles. Together with fusogenic protein, VSV-G, these engineered EVs achieve high-efficiency intracellular delivery of cargo protein (Cre and super repressor of NF-ĸB) or protein complex (Cas9/sgRNA RNPs) both in reporter cells and in mice models.

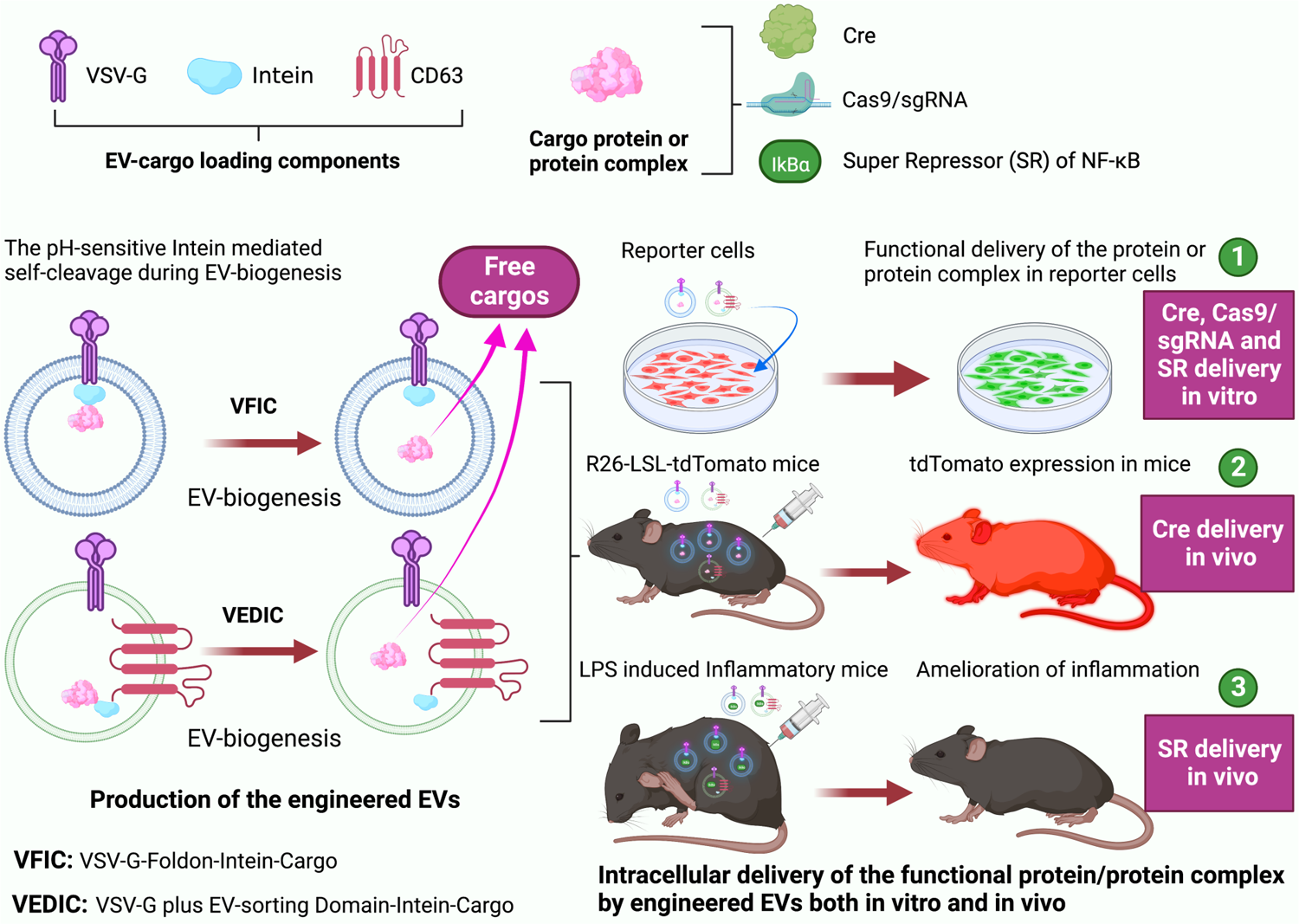

## Introduction

Protein-based therapeutics have a unique potential for the treatment of diverse diseases. While the development of therapeutic proteins directed towards extracellular targets was quite successful with a broad range being approved for clinical application (*1*, *2*), intracellular delivery of proteins remains challenging due to the inherent impermeability of the plasma membrane (*3*, *4*). Thus, numerous strategies have been developed to facilitate intracellular protein delivery: For instance, the iTOP system exploits NaCl-mediated hyperosmolality and transduction compounds to achieve high-efficiency delivery of proteins into primary cells, but has limited potential for *in vivo* applications (*5*). Cell-penetrating peptides (CPPs) have also shown promise in some applications, but suffer from issues with endosomal entrapment and toxicity (*6–8*). Finally, various nanocarriers, such as lipid nanoparticles (LNPs) and polymers, are frequently utilized for intracellular delivery of proteins, but encounter similar limitations to CPPs (*9–13*).

One drawback to these strategies is that their synthetic properties induce various side effects when they are applied *in vivo*. Thus, one strategy to overcome these issues is to harness natural delivery vehicles – extracellular vesicles (EVs). EVs are lipid-bilayer enclosed particles that are secreted and taken up by all cell types to mediate intercellular trafficking of biologically active molecules and thus show great potential for the delivery of therapeutic proteins into recipient cells (*14–16*). However, there are presently two main issues that must be solved in order to achieve efficient intracellular delivery of proteins by EVs: i) Enrichment of therapeutic proteins inside EVs in a soluble (or non-tethered) and active form; and ii) endosomal escape of therapeutic proteins into the target cell cytosol.

The enrichment of specific proteins inside of EVs has previously been achieved by fusing the target proteins to the cytoplasmic domain of EV-sorting proteins, such as CD63 (*17*). However, as these proteins remain bound to the EV membrane, this approach is not suitable for cytosolic delivery of soluble proteins. To address this issue, several technologies introducing cleavable linker peptides between the target protein and EV-sorting domains have been developed to release the bound proteins from the membrane into the EV lumen (*18*, *19*). For example, the EXPLORs system, using an optically reversible protein-protein interaction module, was exploited to liberate soluble protein cargo inside EVs in the absence of blue light (*20*). Although several of the above technologies have achieved intracellular protein delivery by modifying EVs, these solutions were either dependent on extracellular conditions or required multiple components to be co-expressed. In the present study, we innovatively exploited the engineered self-cleavage mini-intein (intein) which was derived from *Mycobacterium tuberculosis (Mtu) recA* (*21*), to connect the EV-sorting protein with the cargo protein, facilitating cargo protein liberation from the EV-sorting protein. This intein comprises the first 110 and the last 58 amino acids of the 441–amino acid *Mtu recA* intein, with four additional mutations (C1A, D24G, V67L, and D422G) introduced to enable C-terminal cleavage in a pH-sensitive manner at 37°C (*21*).

Since endocytosis is the most common mode of biomacromolecule uptake by cells, endosomal entrapment constitutes the primary barrier to the functional intracellular delivery of both therapeutic proteins and EVs (*22*, *23*). Interestingly, fusogenic proteins derived from viruses have been found to mediate endosomal escape and facilitate the release of EV cargo into the cell cytosol (*24*, *25*). In this study, we made use of the the fusogenic protein, vesicular stomatitis virus G glycoprotein (VSV-G), as both an efficient endosomal escape booster and EV-sorting protein. Here, we have developed two systems that solve the aforementioned problems above in tandem to achieve high-efficiency intracellular protein delivery harnessing engineered EVs. Using VSV-G plus EV-sorting Domain-Intein-Cargo (VEDIC) and VSV-G-Foldon-Intein-Cargo (VFIC), we observed efficient intracellular delivery of functional and therapeutic proteins both *in vitro* and *in vivo*. Altogether, this work demonstrates the great potential for EV-based therapeutic protein delivery.

## Results

### Development of the VEDIC system for high-efficiency intracellular protein delivery

To assess the potential of EVs for intracellular delivery of functional proteins, we tested various approaches to load and deliver Cre recombinase. To this end, we designed a range of constructs (Fig. 1A) which were transiently transfected in HEK293T cells. EVs were subsequently harvested from these cells and used to treat Traffic Light (TL) fluorescent Cre reporter cells (Fig. 1B) (*26*). In these cells, LoxP recombination by Cre results in the excision of the red fluorescent protein (RFP) DsRed which subsequently leads to the permanent expression of green fluorescent protein (GFP), allowing the detection of successful Cre protein delivery with single-cell resolution (Fig. 1B). EVs from cells overexpressing Cre alone or CD63-Cre (Cre fused to CD63 to enhance EV-enrichment) did not induce any recombination in reporter cells analyzed by flow cytometry (Fig. 1D and fig. S1, A and C). To liberate Cre from CD63 inside the EV lumen, the intein was introduced between CD63 and Cre (Fig. 1, A and C) (*21*). However, no recombination was observed in reporter cells upon treatment with CD63-Intein-Cre EVs either, despite confirmation by Western Blot analysis that Cre was enriched and released from CD63 in the isolated EVs (Fig. 1D and fig. S1D). Previous studies have shown that EV endosomal escape following endocytosis is a limiting factor for cargo delivery, however fusogenic viral proteins enhanced EV cargo endosomal escape (*27*). Therefore, a comprehensive screen of 40 human- and two virus-derived fusogenic proteins was performed to identify efficient, potentially human-derived, fusogenic proteins. These constructs were co-transfected with CD63-Intein-Cre, and EVs were once again added to the reporter cells. While none of the human-derived fusogenic proteins induced recombination, the fusogenic viral protein VSV-G significantly boosted Cre delivery, so that 66% and 98% of HeLa-TL and T47D-TL cells, respectively, expressed GFP two days after treatment with engineered EVs (Fig. 1E and fig. S1B). The other fusogenic viral protein tested, EPFV (envelope of prototype foamy virus), also enhanced recombination in reporter cells, though to a lesser extent (Fig. 1E and fig. S1B). Subsequently, we co-expressed VSV-G in the EV producing cells alongside our previously engineered constructs; Cre, CD63-Cre, Intein-Cre and CD63-Intein-Cre, and incubated the isolated EVs with reporter cells for 48 hours (h). Of the conditions tested, only EVs carrying VSV-G and CD63-Intein-Cre achieved significant activity in reporter cells (Fig. 1F), indicating that the EV-sorting domain (CD63), self-cleavage protein (intein) and endosomal escape booster (VSV-G) were indispensable for the intracellular delivery of Cre by engineered EVs. We term this approach the VEDIC (VSV-G plus EV-Sorting Domain-Intein-Cargo) system.

**Fig. 1.**
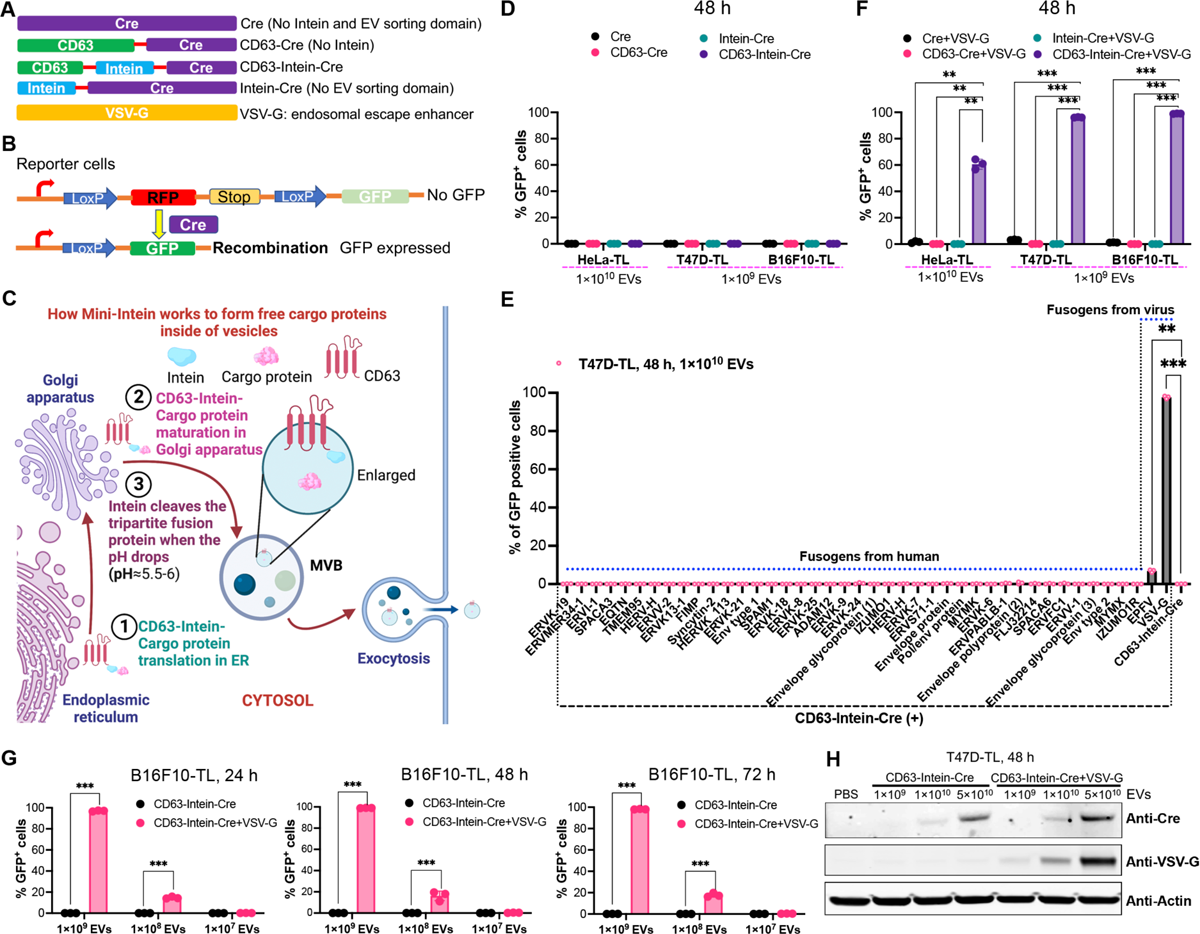
Development of the VEDIC system for high-efficiency intracellular protein delivery by EVs. (**A**) Schematic of constructs used for developing the VEDIC system. (**B**) Schematic of intein cleavage and intraluminal cargo release during EV-biogenesis. (**C**) Schematic of fluorescence reporter construct expressed in the reporter cells generated to measure Cre delivery. (**D**) Percentage of GFP positive reporter cells after adding EVs for 2 days, as evaluated by flow cytometry. Unless indicated, the experiments were performed in 96-well plates. (**E**) Fusogen screen in T47D-TL cells after a two-day incubation period with EVs. (**F**) Percentage of GFP positive reporter cells after exposure to EVs derived from VSV-G co-transfected cells. (**G**) EV dose-dependent recombination in B16F10-TL reporter cells mediated by CD63-Intein-Cre+VSV-G EVs. (**H**) Cre and VSV-G protein was detected in T47D-TL reporter cells by Western blot analysis, 48 hours (h) after addition of engineered EVs loaded with Cre in 24-well plates. Two-way ANOVA multiple comparisons test was used for analysis of (F) and (G); One-way ANOVA multiple comparisons test was used for analysis of (E). Data are shown as mean±SD, * p < 0.05; ** p < 0.01; *** p < 0.001.

One potential limiting factor for this strategy is the required presence of both VSV-G and CD63-Intein-Cre in the same vesicles. To explore this, we co-transfected HEK293T cells with VSV-G-mNeonGreen (VSV-G-mNG) and CD63 expression plasmids, and performed image-based single-vesicle flow cytometry on the isolated vesicles using an APC-conjugated anti-CD63 antibody according to the method we published before (*28*). This showed a high degree of co-localization, as 84% of mNG+ vesicles were APC+, while 86% of the APC+ vesicles were mNG + (fig. S1E). Next, different doses of the VEDIC EVs were added to various reporter cell lines, which were assayed at multiple time points. We observed a clear pattern of dose- and time-dependent recombination in B16F10-TL, HeLa-TL and T47D-TL cells, as observed by flow cytometry and fluorescent microscopy (Fig. 1G, fig. S2, A and B, and fig. S3, A and B). For VEDIC EVs, a time-lapse video obtained from live imaging of HeLa-TL recipient cells showcased the gradual recombination occurring over time (Movie S1), while no GFP signal was observed in the absence of VSV-G (Movie S2). Furthermore, the protein expression levels of Cre and VSV-G was evaluated in T47D-TL and B16F10-TL cells and found to correlate with the activation of GFP signal in these cell lines (Fig. 1H and fig. S3D). Moreover, VEDIC EVs were applied to multiple hard-to-transfect cell lines (MSC-TL, THP-1-TL, Raw264.7-TL, and K562-TL cells) and significant recombination was observed 2 days after EV treatment in all cases (fig. S2, C to F and fig. S3C). Furthermore, CD63 was replaced with other known EV-sorting domains (CD81, CD9, and PTGFRN) and found that addition of EVs isolated from cells transfected with all these constructs resulted in significant recombination in recipient cells when combined with VSV-G (fig. S3E). For most of the selected EV-sorting domains (CD63, CD81, and CD9), the recombination levels achieved after treatment with high-dose of EVs outperformed the efficiency of direct transient transfection of Cre plasmids using Lipofectamine 2000 in reporter cells (Fig 1F, fig. S3E and fig. S4).

In addition to adding engineered EVs to reporter cells, the EV-mediated recombination efficiency in co-culture was assessed: using direct co-culture (co-culture), IBIDI co-culture µ-slide (IBIDI) and Transwell co-culture (Transwell) assays (fig. S5A and S6A) (*29*). For direct co-culture assays, EV-producing and reporter cells were cultured at various ratios and GFP levels were evaluated one day after incubation (fig. S5A). Here, VEDIC cells enabled significant recombination in various reporter cells (fig. S5, B to E). For IBIDI assays, reporter cells were seeded in the inner reservoir as shown in fig. S6A, and the feeder cells (EV-producing cells) were seeded in the 8 surrounding reservoirs. After 4 days of incubation, co-culture with the VEDIC donor cells exhibited pronounced levels of GFP activation in the reporter cells (fig. S6, B and E). For Transwell assays, the reporter cells were seeded in the bottom compartment and the EV-producing cells in the upper chamber. In line with the previous results, significant increases in GFP positive cells were only observed after co-culture with the VEDIC donor cells, after 4 days of incubation (fig. S6, C, D and F).

### Development of the VFIC system to further improve intracellular protein delivery by EVs

To further improve the VEDIC system, which requires transfection of several plasmids, we aimed to combine all the essential components for intracellular EV-cargo delivery into a single construct. To determine whether VSV-G itself could be employed as an efficient EV-loading protein, VSV-G-mNG was transfected into HEK293T cells and the isolated vesicles were analyzed by single-vesicle flow cytometry after the EVs were stained with a CD63-APC antibody. The total number of mNG+ vesicles was far greater than CD63+ vesicles (Fig. 2 A). When VSV-G-mNG was co-transfected with CD63 plasmid in HEK293T cells, the total number of isolated mNG+ vesicles was similar to that of CD63+ vesicles (Fig. 2 B). These data imply that VSV-G could function as an efficient EV-sorting domain, similar to CD63. Thus, a VSV-G-Intein-Cre fusion protein was constructed, with and without a foldon component that has been previously demonstrated to enhance VSV-G trimer formation and function, and confirmed their expression (fig. S8C). A negative control construct was also generated whereby Cre was replaced with the bacteriophage protein MS2 (Fig. 2C and fig. S8C) (*30–32*). Upon adding these EVs to reporter cells, a dose- and time-dependent recombination in reporter cells was observed (Fig. 2, D and E, fig. S7, and fig. S8, A and B), and nearly 100% recombination in B16F10-TL and T47D-TL cells was detected in the high-dose treated groups (Fig. 2D, fig. S7B, and fig. S8A). The VSV-G-Foldon-Intein-Cre performed better than the VSV-G-Intein-Cre construct in low EV-dose experiments (Fig. 2D and fig. S7, A and B), demonstrating the ability of the foldon domain to enhance VSV-G loading and/or function. We term this VSV-G-Foldon-Intein-Cargo system VFIC. Time-lapse fluorescence microscopy video analysis revealed similar recombination dynamics by VFIC EVs as was previously observed using the VEDIC system in HeLa-TL cells (Movie S3). VFIC EV-mediated recombination was also observed in hard-to-transfect cells (MSC-TL, THP-1-TL, Raw264.7-TL, and K562-TL cells) (Fig. 2, F-I and fig. S8B). We subsequently performed the direct co-culture, IBIDI, and Transwell assays using the VFIC system, and obtained significant recombination in recipient cells by transfer of Cre protein from donor cells (fig. S9). It is worth noting that recombination was observed with as few as one EV-producing cell per fifty reporter cells in direct co-culture assays (Fig. S10). The recombination level achieved by high dose of VFIC EVs outperformed the efficiency observed by direct transient transfection of Cre plasmids using Lipofectamine 2000 in reporter cells (Fig. 2D, fig. S7, A and B, and fig. S11), again confirming the high efficiency of the engineered EVs.

**Fig. 2.**
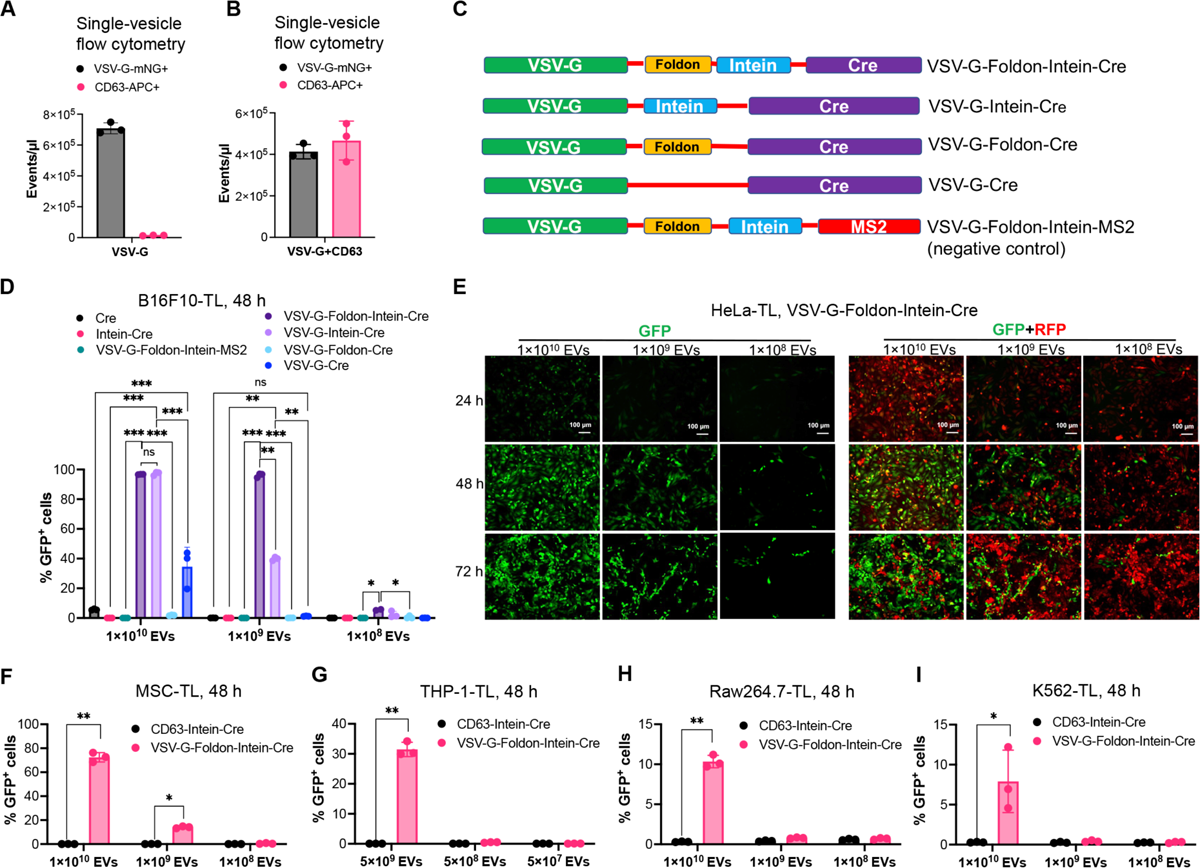
Development of the VFIC system to further improve intracellular protein delivery by EVs. (**A** and **B**) VSV-G+ and CD63+ vesicle numbers as determined by single-vesicle flow cytometry after transfection with VSV-G-mNeonGreen alone or VSV-G-mNeonGreen and CD63 together. (**C**) Schematic of constructs generated for developing the VFIC system. (**D**) EV dose-dependent recombination in B16F10-TL cells mediated by VSV-G-Foldon-Intein-Cre and VSV-G-Intein-Cre EVs as evaluated by flow cytometry. (**E**) Representative images showing GFP positive HeLa-TL cells 24, 48 and 72 h after exposure to VFIC EVs at different doses. Scale bar, 100 µm. (**F-I**) Recombination in hard-to-transfect reporter cells (MSC-TL, THP-1-TL, Raw264.7-TL and K562-TL) mediated by VFIC EVs after 48 h. Two-way ANOVA multiple comparisons test was used for analysis of (D), and (F) to (I). Data are shown as mean±SD, * p < 0.05; ** p < 0.01; *** p < 0.001; ns: non-significant.

### The pH-sensitive intein performs C-terminal cleavage during EV-biogenesis

To confirm that the intein we used in this study performed C-terminal cleavage in a pH-sensitive manner, we introduced 2 mutants (H439Q and N440A) (Fig. 3A). The H439Q variant has a lower pH-sensitivity, which should lead to a reduced cleavage rate during EV-biogenesis, since the pH in MVBs is ∼6 (Fig. 3A, middle panel) (*33*). In contrast, the N440A variant is expected to abolish intein C-terminal cleavage activity, which should lead to minimal cleavage both in EV-producing cells and the isolated EVs (Fig. 3A, lower panel) (*34*). As anticipated, the decrease of C-terminal cleavage leading to reduced Cre cargo protein release was corroborated by western blot for the VEDIC system (Fig. 3B). Accordingly, the functional assays (adding isolated EVs to recipient cells or direct co-culture of VEDIC producer cells and reporter cells) indicated significantly decreased recombination in reporter cells when the 2 mutants were included in the VEDIC system (Fig. 3, C to E, fig. S12, and fig. S14, A and B). These results were further corroborated for the VFIC system (fig. S13 and fig. S14, C and D). Altogether, these findings support the notion that the intein used for the VEDIC and VFIC technologies enables robust C-terminal cleavage to form soluble cargo proteins in the EV-lumen during EV-biogenesis in a pH-dependent manner (Fig. 1C).

**Fig. 3.**
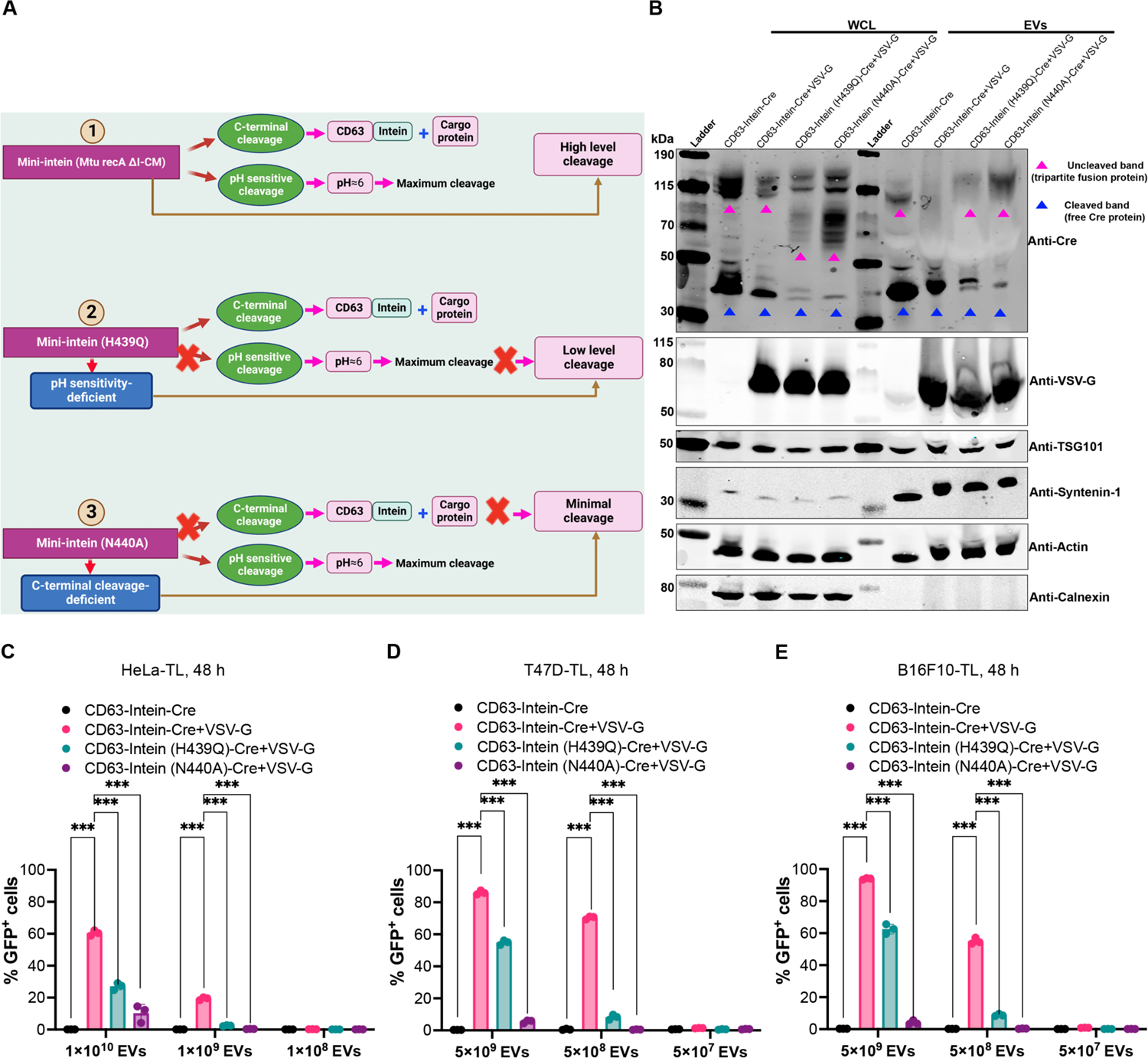
The pH-sensitive intein performs C-terminal cleavage during EV-biogenesis. (**A**) Schematic illustration of the cleavage mechanism of various engineered intein variants. (**B**) Protein expression of various engineered mutant intein constructs in whole cell lysates (WCL) and isolated EVs derived from HEK-293T cells evaluated by western blot analysis. Lysates from 5×10^5^ EV-producing cells and 1×10^10^ engineered EVs were used for the assay. TSG101, syntenin-1 and β-actin were used as EV markers and Calnexin was used as a cellular organelle marker (endoplasmic reticulum) and should be absent in EV samples. (**C** to **E**) Recombination in reporter cells mediated by EVs derived from engineered cells using mutant inteins (H439Q and N440A) in the VEDIC system. Two-way ANOVA multiple comparisons test was used for analysis of (C) to (E). Data are shown as mean±SD, * p < 0.05; *** p < 0.001. ns: non-significant.

### VSV-G boosts endosomal escape following receptor-mediated endocytosis in recipient cells

To ascertain the role of VSV-G in endosomal escape and endocytosis, we introduced mutations that have been described to disrupt the fusogenic (VSV-G P127D) or LDL-R receptor binding capacity (VSV-G K47Q), respectively (Fig. 4, A and B) (*35–37*). In order to assess EV uptake and trafficking, Cre was replaced by mNG in the VEDIC system, generating a CD63-intein-mNG construct, which was co-transfected with VSV-G to produce EVs that were added to Huh7 cells, a hepatic cell line that has been shown previously to efficiently take up EVs (*38*). Confocal microscopy was used to evaluate the uptake of the vesicles 48 h after incubation and showed a punctate distribution of green fluorescent signal in recipient cells in the absence of VSV-G, indicating endosomal entrapment (Fig. 4C). However, when VSV-G was included, the mNG signal diffused into the cytosol (Fig. 4C), suggestive of endosomal escape. Of note, when VSV-G P127D was used, mNG instead exhibited a punctate distribution (Fig. 4D), confirming that endosomal escape was mediated by the fusogenic activity of VSV-G. In line with our hypothesis, mNG was furthermore observed in punctate form when a directly fused CD63-mNG (without intein) was co-transfected with VSV-G (Fig. 4D). This observation confirms that a strategy to release cargo from the EV membrane, such as the use of an intein, is required to facilitate release of the protein into the cytosol of the recipient cell (Fig. 1F).

**Fig. 4.**
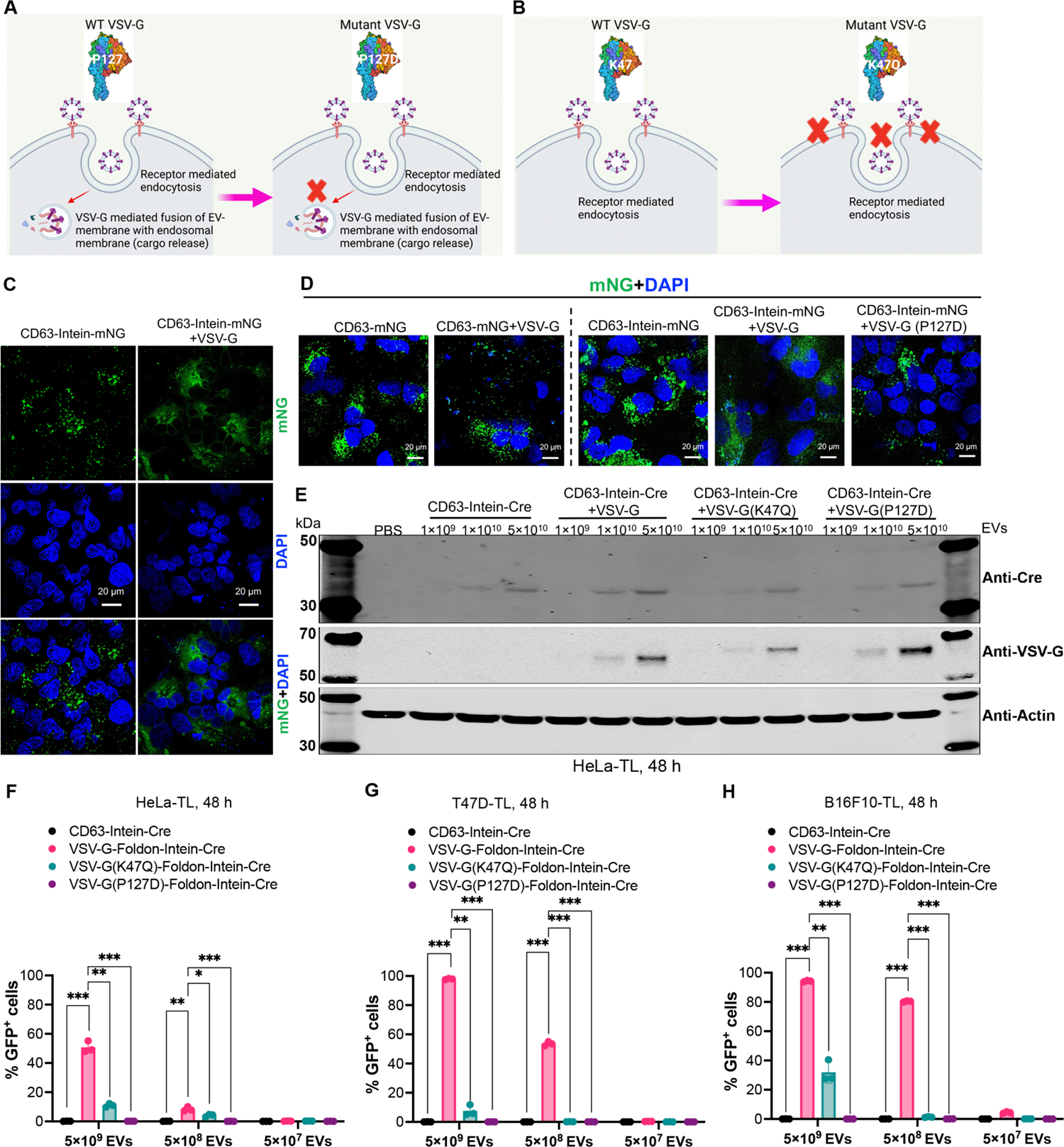
VSV-G enhances endosomal escape following receptor-mediated endocytosis of VSV-G-engineered EVs into recipient cells. (**A** and **B**) Properties of the two VSV-G mutants: VSV-G P127D loses the capacity to mediate fusion between the EV-and endosomal membranes and VSV-G K47Q is unable to bind to LDLR on the cell surface. (**C**) Confocal immunofluorescence demonstrating the subcellular distribution of mNG in the presence or absence of wild type VSV-G engineered EVs in Huh7 cells. Scale bar, 20 µm. (**D)** Subcellular distribution of mNG in different groups after adding the indicated engineered EVs determined by confocal immunofluorescence. Scale bar, 20 µm. (**E**) Western blot evaluation of protein levels of Cre and VSV-G in HeLa-TL reporter cells after addition of engineered EVs with wild type, P127D or K47Q VSV-G in 24-well plates. (**F**-**H**) Percentage of GFP positive HeLa-TL, T47D-TL and B16F10-TL cells after adding wild type, P127D or K47Q VSV-G CD63-Intein-Cre EVs, as evaluated by flow cytometry. Two-way ANOVA multiple comparisons test was used for analysis of (F) to (H). Data are shown as mean±SD, * p < 0.05; ** p < 0.01; *** p < 0.001. ns: non-significant.

Next, the two mutant VSV-G constructs were used for Cre protein delivery. Decreased Cre protein delivery with the same doses of EVs in HeLa-TL recipient cells was observed using western blot analysis for both mutants compared to the wide-type VSV-G group, given that free-Cre protein level was equal in these EV groups (Fig. 4E and fig. S15E) (*37*). Utilization of the two VSV-G mutants in the VEDIC system resulted in significantly decreased Cre-mediated recombination in reporter cells with VSV-G K47Q and complete loss of reporter activity with VSV-G P127D (fig. S15, A to D, and fig. S16). Furthermore, we reproduced these results using the VFIC system (Fig. 4, F to H, fig. S17, and fig. S18), which supports the hypothesis that VSV-G functions both to facilitate endosomal escape and to induce receptor-mediated endocytosis of engineered EVs into recipient cells.

### Robust gene editing by Cas9/sgRNA RNPs and meganuclease targeting PCSK9 using the VEDIC and VFIC systems

Since Cre and mNG proteins were successfully delivered using the VEDIC and VFIC systems, we next tested whether these delivery strategies could be applied to more relevant therapeutic cargos. To this end, Cre was replaced with Cas9 and CD63-Intein-Cas9 and VSV-G-Foldon-Intein-Cas9 fusion proteins were generated to encapsulate Cas9-ribonucleoprotein (Cas9-RNP) into engineered EVs (Fig. 5, A and B, and fig. S19B). As a read-out for Cas9-ribonucleoprotein delivery, we employed CRISPR reporter (Stoplight, SL) cells that constitutively express mCherry followed by a short linker region that includes a sgRNA target region and a stop codon, followed by two eGFP open reading frames (ORFs), +1nt and +2nt out of frame respectively. Upon successful Cas9-RNP delivery, non-homologous end joining (NHEJ)-mediated +1nt and +2nt frameshifts in the targeting linker region will result in bypassing of the stop codon and the permanent expression of eGFP. (Fig. 5C) (*39*). Significant dose- and time-dependent genome editing was observed in cells treated with EVs derived from HEK293T cells expressing either CD63-Intein-Cas9 + VSV-G + a targeting sgRNA or VSV-G-Foldon-Intein-Cas9 + a targeting sgRNA EVs (Fig. 5, D and E, and fig. S19A), indicating successful Cas9-RNP delivery. For VSV-G-Foldon-Intein-Cas9-RNP, close to 80% gene editing efficiency was achieved, which is most likely the maximum achievable for this reporter construct and significantly higher than demonstrated by directly transfecting Cas9 and sgRNA plasmids into the reporter cells (fig. S19C) (*39*). Interestingly, the efficiency of VSV-G-Foldon-Intein-Cas9-RNPs was significantly higher than that of CD63-Intein-Cas9-RNP + VSV-G, suggestive of a potential advantage of the VFIC system over the VEDIC system for CRISPR delivery (Fig. 5, D and E).

**Fig. 5.**
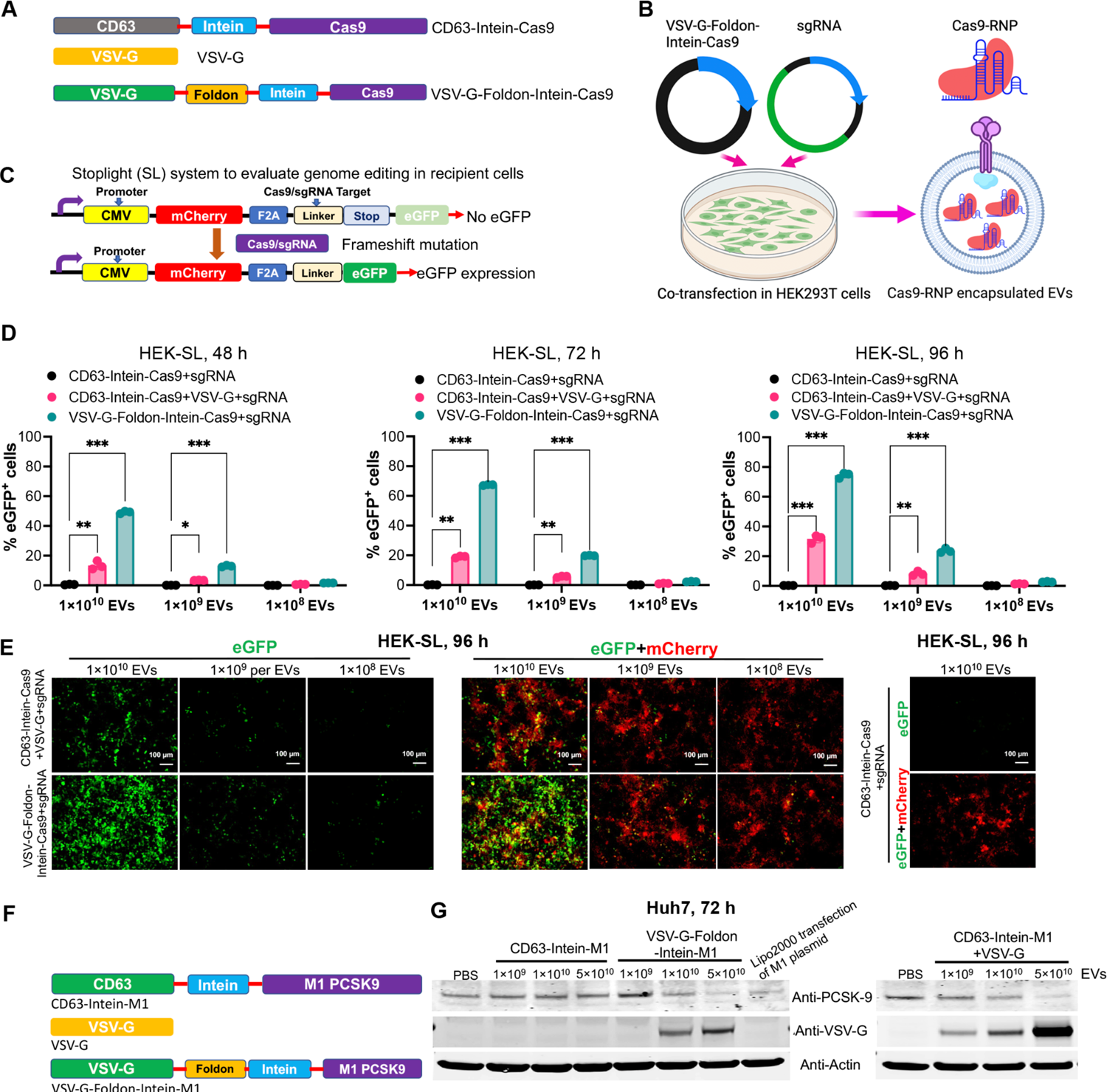
Robust gene editing by Cas9/sgRNA RNP and meganuclease targeting PCSK9 using the VFIC and VEDIC systems. (**A**) Constructs generated for Cas9/sgRNA RNPs delivery. **(B)** Schematic illustration about how the Cas9/sgRNA RNPs were encapsulated into engineered EVs. (**C**) Schematic of reporter cells used to functionally assess Cas9/sgRNA RNPs delivery by engineered EVs. (**D**) Percentage of eGFP positive cells after addition of engineered EVs, as measured by flow cytometry at 48, 72 and 96 h after EV addition. (**E**) Immunofluorescence demonstrated gene-editing in recipient cells after treatment with different doses of engineered EVs after 4 days. Scale bar, 100 µm. (**F**) Constructs generated for EV-mediated delivery of meganuclease targeting PCSK9. (**G**) Western blot analysis of PCSK9 and VSV-G protein delivery in Huh7 cells after treatment with different doses of EVs in 24-well plates. Two-way ANOVA multiple comparisons test was used for analysis of (D). Data are shown as mean±SD, * p < 0.05; ** p < 0.01; *** p < 0.001.

Next, VEDIC and VFIC constructs for the delivery of a previously described meganuclease targeting *PCSK9* were generated. Upon EV exposure to cells, a significant decrease of PCSK9 protein levels was observed in a dose-dependent manner (Fig. 5, F and G) (*40*).

### Cre-mediated recombination in melanoma-xenograft and R26-LSL-tdTomato reporter mice by VEDIC and VFIC systems after local injection

Based on the promising results achieved *in vitro*, the *in vivo* applicability of VEDIC and VFIC-engineered EVs was assessed. To evaluate VEDIC and VFIC-mediated Cre delivery *in vivo*, an intratumoral (IT) injection of engineered EVs was conducted in C57BL/6 mice bearing subcutaneous B16F10-TL melanoma xenografts (Fig. 6A). 4 days after injection, tumors were harvested for immunohistochemistry analysis, which showed significant GFP signals following treatment with VEDIC and VFIC EVs, demonstrating successful Cre delivery (Fig. 6B). Next, R26-LSL-tdTomato reporter mice, whereby successful Cre delivery excises stop cassettes that are present between the CAG promoter and an ORF of the red fluorescent protein, tdTomato, resulting in its expression, were exploited for *in vivo* study. These mice were injected with VEDIC and VFIC EVs through intracerebroventricular (ICV) injection to assess Cre delivery in the brain (Fig. 6C) (*26*). One week after injections, the brains were harvested and stained for tdTomato expression. As shown in Fig. 6D, tdTomato expression was detected in the cerebellum, cortex, and hippocampus of VEDIC- and VFIC EV treated mice, and to a much lesser extent in mice treated with CD63-Intein-Cre alone, albeit the CD63-intein-Cre EVs did not generate recombination *in vitro* without VSV-G (Fig. 6D). Moreover, we additionally observed trace amounts of tdTomato expression in the olfactory bulb and thalamus of VEDIC and VFIC treated animals (fig. S20A). To further identify which cell types internalized the engineered EVs, the slides were co-stained for both tdTomato and specific cell-marker genes. We found apparent colocalization of tdTomato with GFAP (astrocyte marker) and IBA1 (microglia marker), but only marginal colocalization of NeuN (neuronal marker) in the corpus callosum and hippocampus in the engineered EV-treated animals (Fig. 6, E to G and fig. S20, B and C). Based on the above results, we concluded that the engineered EVs mainly delivered their cargo to astrocytes and microglia in the brain.

**Fig. 6.**
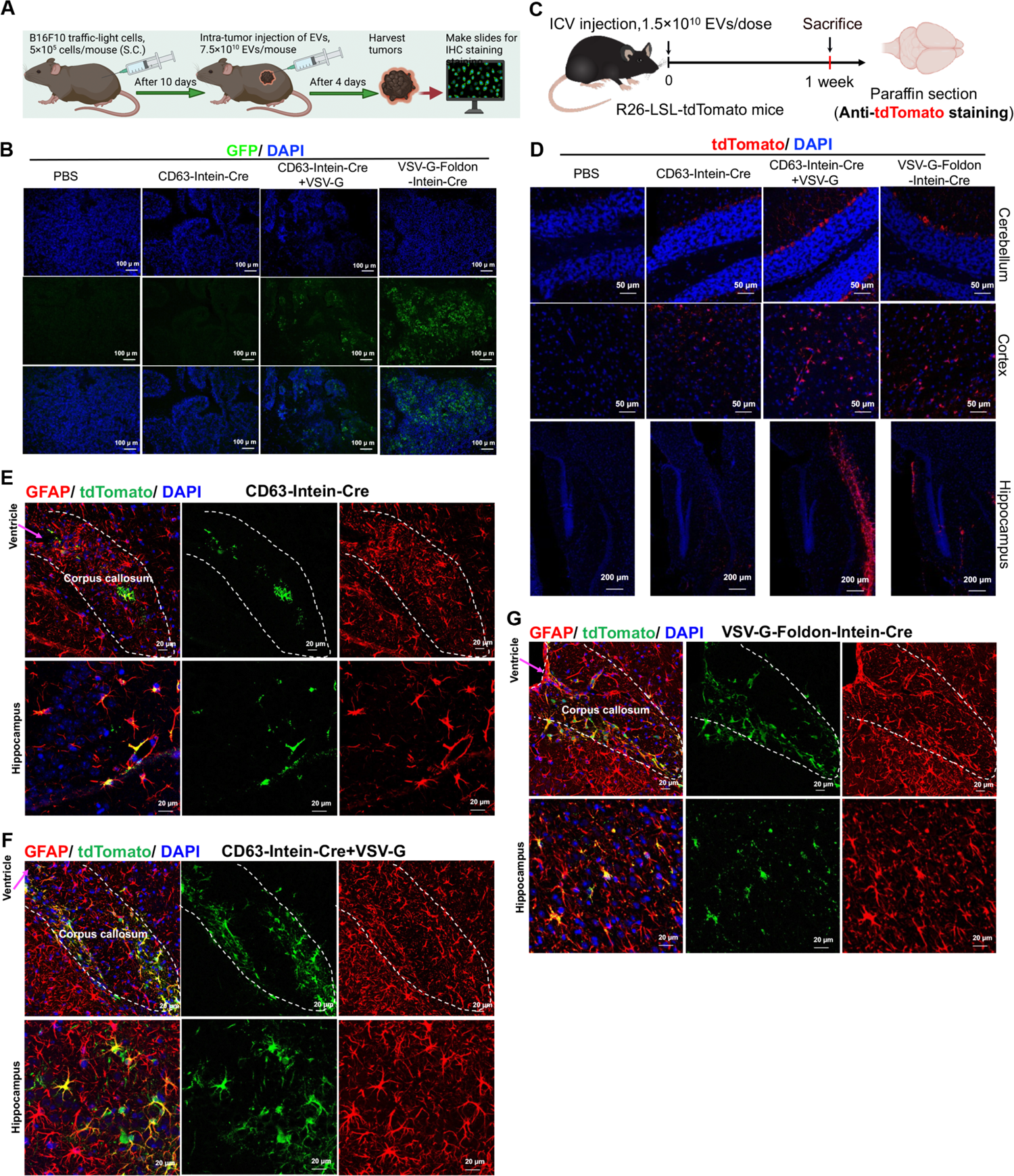
Cre recombination in melanoma-xenograft and Cre-LoxP R26-LSL-tdTomato reporter mice by VEDIC and VFIC EV-mediated Cre delivery after local (ICV) injection. **(A)** Workflow for the intratumoral injection model. (**B**) Representative immunofluorescence images from tumor tissues after intratumoral injection of engineered EVs to induce GFP expression. Scale bar, 100 µm. n=3 mice per group. (**C**) Workflow for the intracerebroventricular (ICV) injection of engineered EVs to deliver Cre in the brains of Cre-LoxP R26-LSL-tdTomato reporter mice. (**D**) TdTomato expression in different regions of brain after ICV injection of engineered EVs, as determined by immunofluorescence. Scale bar, 50 µm for cerebellum and cortex and 200 µm for hippocampus. n=3 mice per group, representative images. (**E** to **G**) Co-staining of tdTomato with the astrocyte marker GFAP in corpus callosum (outlined by dashed lines) and hippocampus one week after ICV injection of engineered EVs. Scale bar, 20 µm. n=3 mice per group, representative images.

### Cre recombination in R26-LSL-tdTomato reporter mice following systemic VEDIC and VFIC EV-mediated Cre delivery

One pilot *ex vivo* study was performed by adding engineered EVs to liver primary cells harvested from R26-LSL-tdTomato reporter mice and significant recombination of the cells by VEDIC and VFIC EVs was found (fig. S21A). Next, high-doses of engineered EVs were administered via intraperitoneal (IP) injection into R26-LSL-tdTomato reporter mice. The liver, spleen and heart were harvested for immunofluorescence analysis one week after injection (Fig. 7A). A substantial number of cells in the liver and spleen were found to be tdTomato positive following injection of both VEDIC and VFIC EVs, but not after injection of CD63-Intein-Cre EVs (Fig. 7B). In contrast, we did not observe any significant tdTomato expression in the heart (fig. S21B).

**Fig. 7.**
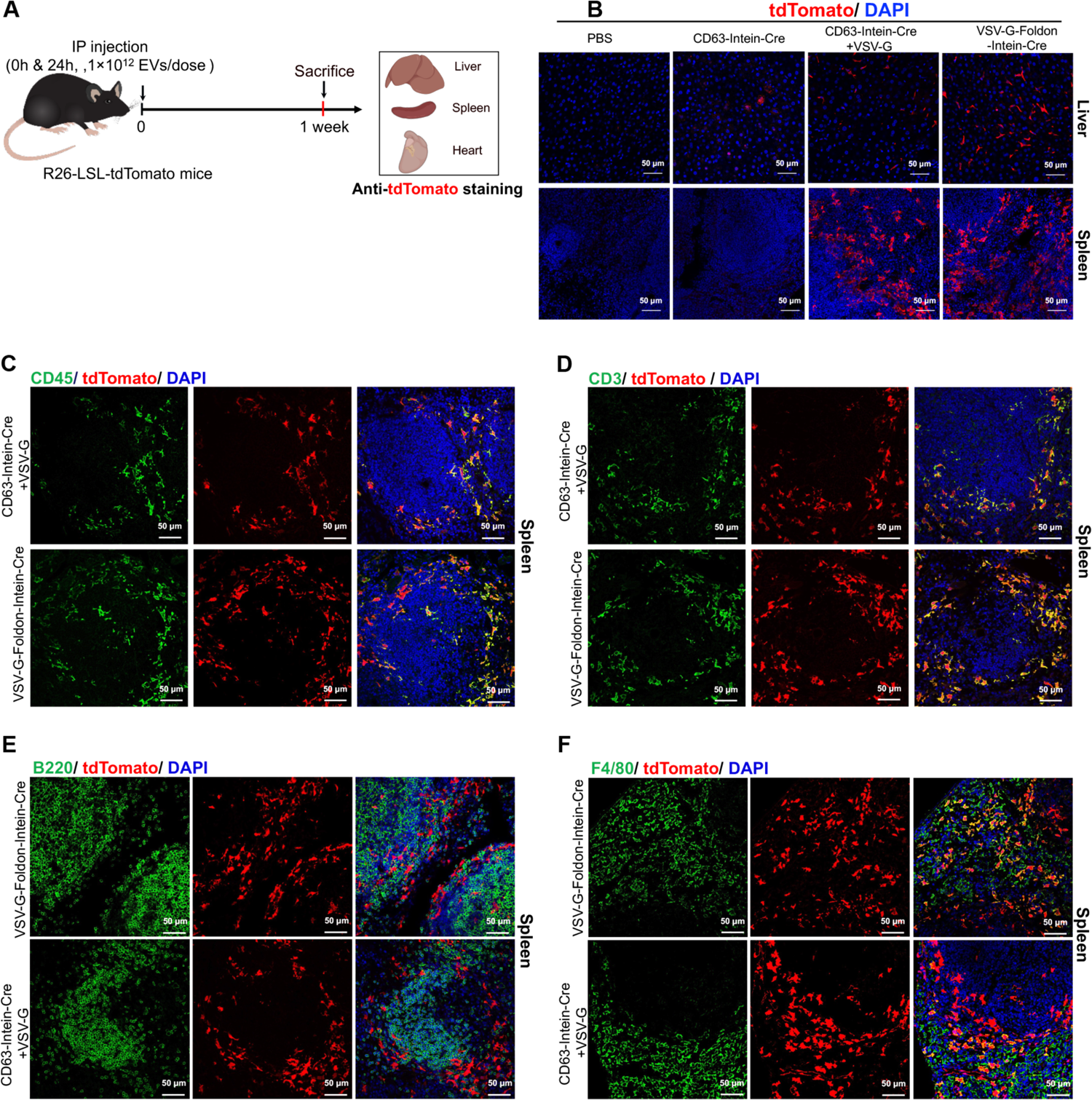
Cre recombination in R26-LSL-tdTomato reporter mice following systemic VEDIC and VFIC EV-mediated Cre delivery. (**A**) Schematic workflow for the intraperitoneal (IP) injection of engineered EVs into Cre-LoxP R26-LSL-tdTomato reporter mice. (**B**) TdTomato expression in liver and spleen after IP injection of engineered EVs one week later. Scale bar, 50 µm. n=3 mice per group, representative images. (**C**) Co-staining of tdTomato with the general leukocyte marker CD45 in spleen as detected by immunofluorescence one week after IP injection of engineered EVs. Scale bar, 50 µm. n=3 mice per group, representative images. (**D**) Co-staining of tdTomato with the T cell marker CD3 in spleen as detected by immunofluorescence one week after IP injection of engineered EVs. Scale bar, 50 µm. n=3 mice per group, representative images. (**E**) Co-staining of tdTomato with the B cell marker B220 in spleen one week after IP injection of engineered EVs. Scale bar, 50 µm. n=3 mice per group, representative images. (**F**) Co-staining of tdTomato and the macrophage marker F4/80 in spleen one week after IP injection of engineered EVs. Scale bar, 50 µm. n=3 mice per group, representative images.

After co-staining of tdTomato together with cell specific markers, a high degree of functional delivery to leukocytes (CD45 positive), especially to the T cell population (CD3) and macrophages (F4/80), was detected (Fig. 7, C, D, and F, and fig. S21, C, D, and F). In contrast, B cells (B220) showed low recombination events *in vivo* (Fig. 7E and fig. S21E).

### Treatment of Lipopolysaccharide-induced systemic inflammation by VEDIC- and VFIC-mediated delivery of the NF-ĸB super-repressor

In order to demonstrate the applicability of our systems for the treatment of disease, we applied the VEDIC and VFIC EVs to treat lipopolysaccharide (LPS)-induced systemic inflammation by delivering a previously reported super-repressor of NF-ĸB activity (SR) (fig. S22A) (*41*). To accomplish this, CD63-Intein-SR, VSV-G-Intein-SR, and VSV-G-Foldon-Intein-SR constructs were generated, and HEK-NF-ĸB luciferase reporter cells were used as a read-out for functional *in vitro* assessment of the system (Fig. 8, A and B, and fig. S22D). In this reporter cell line, IĸB is degraded upon inflammatory stimuli such as LPS or TNF-α, which allows NF-ĸB to translocate to the nucleus and drive downstream luciferase reporter gene expression, since luciferase is driven by a minimal NF-ĸB promoter (Fig. 8C) (*42*). First, we confirmed that the NF-ĸB SR, which is constitutionally active and maintains the NF-ĸB in the cytoplasm, inhibited the HEK-NF-ĸB luciferase reporter activation (fig. S22, B and C) (*43*). SR delivered by VSV-G-Foldon-Intein-SR, VSV-G-Intein-SR, and VSV-G+CD63-Intein-SR EVs successfully inhibited TNF-α-mediated NF-ĸB signaling activation evidenced by decreased reporter gene expression (Fig. 8, D and E). This provided the rationale for testing these EVs in a murine LPS-induced model of systemic inflammation (Fig. 8F). Thus, engineered EVs were injected 4 h before and 6 h after injecting LPS to allow for, and enhance, the binding of EV-delivered SR to NF-ĸB, respectively. Subsequently, the weight and mortality of the mice were measured at 24 and 48 h after LPS treatment. Compared to the PBS and CD63-Intein-SR injected groups, the body weight and survival of VSV-G+CD63-Intein-SR and VSV-G-Foldon-Intein-SR treated animals was significantly improved after 48 h (Fig. 8, G and H). Histological assessment of the liver revealed a decrease in inflammatory cells at the portal areas as well as a significant alleviation of the hydropic degeneration of hepatocytes by treatment with both VEDIC and VFIC engineered EVs (Fig. 8I). These results demonstrate the adaptability and therapeutic potential of the VEDIC and VFIC systems for the treatment of disease.

**Fig. 8.**
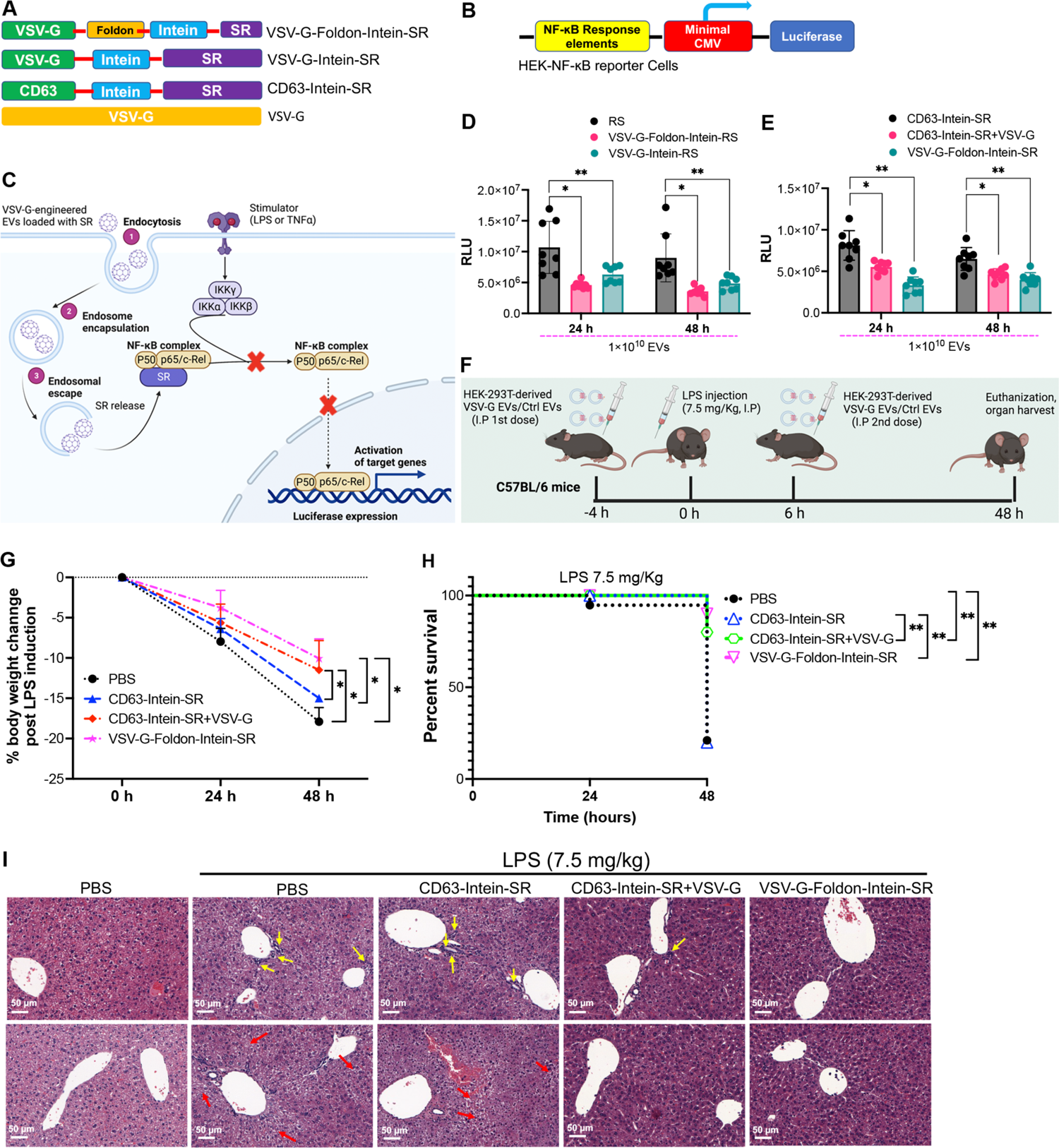
Treatment of LPS-induced systemic inflammation using VEDIC-EV and VFIC-EV mediated delivery of a super-repressor of NF-ĸB. (A and B) Design of the constructs and reporter cells utilized for delivery and assessment of a super-repressor of NF-ĸB by engineered EVs. (C) Schematic illustration how the EV-delivered super-repressor of NF-ĸB inhibit NF-ĸB activity. (D and E) Luciferase activity from HEK-NF-ĸB reporter cells, 24 or 48 h after treatment with engineered EVs respectively (TNF-α stimulation for 6 h before harvesting cells), in 24-well plates. (F) Schematic illustration of the workflow for the treatment of LPS-induced systemic inflammation by engineered EVs in mice (5×10^11^ EVs/mouse). (G and H) Percentage body weight loss and group survival in mice after LPS and engineered EV injections. n=10 mice per group. (I) Representative histology images (hematoxylin-eosin stain) of liver to show the aggregation of inflammatory cells (upper panel, yellow arrows indicate the aggregated inflammatory cells in portal areas) and the hydropic degeneration of hepatocytes (lower panel, red arrows indicate the hydropic degeneration of hepatocytes) after LPS induction. Scale bar = 50 µm. Two-way ANOVA multiple comparisons test was used for analysis of (D), (E) and (G); Log-rank (Mantel-Cox) test was used for the analysis of survival curve. Data are shown as mean±SD for (C), (D) and (F). * p < 0.05; ** p < 0.01.

## Discussion

By fine-tuning our engineering strategies, we have identified solutions to the major bottlenecks, the enrichment of free-form active cargo into EVs and their endosomal escape in recipient cells, to EV-mediated intracellular protein delivery. The resulting VEDIC and VFIC systems achieved an unprecedented level of efficiency for EV-mediated intracellular delivery of functional proteins *in vitro* and *in vivo*. Owing to the versatility of the VEDIC and VFIC systems, we anticipate that many other therapeutic proteins or protein complexes of interest could be efficiently delivered, apart from Cre, super-repressor of NF-ĸB and Cas9-RNPs explored in this work. As such, these approaches may hold great potential for the further therapeutic development.

This work has demonstrated that functional cargo delivery by EVs requires a simultaneous application of strategies to enrich and liberate cargo inside EVs, as well as to induce their release from endosomes in recipient cells. By using an intein to connect an EV-sorting domain to a cargo protein, combined with the fusogenic protein VSV-G on the EV-surface, we developed VEDIC system to achieve robust intracellular delivery of functional cargos. Within our targeted loading and delivery strategies, we found that both the EV-sorting and fusogenic proteins could be replaced by alternative proteins. However, after screening a large number of candidate proteins, we found that CD63 and VSV-G were the most potent mediators of these processes, respectively. Importantly, there is very likely space for improvement via the identification of more potent loading scaffolds and fusogenic proteins.

One of the major innovations of this study is to utilize the self-cleaving engineered intein to liberate cargo proteins in their soluble active form inside of engineered EVs, facilitating cargo release from the endosomal compartment to the cytosol in recipient cells. Inteins are naturally occurring, self-splicing proteins which excise themselves from a precursor protein to join the flanking regions (exteins) to restore host gene function (*44*). After random mutagenesis and genetic selection, the engineered pH-sensitive intein derived from *Mtu recA,* which performs C-terminal cleavage, was identified (*21*). This engineered intein, which cleaved most efficiently when the pH is reduced to 6 at 37℃, was utilized in affinity fusion-based protein purification (*45*). These properties of the engineered intein were perfectly adapted for EV-engineering to achieve optimal amount of released active cargo (C-terminal cleavage) inside of the EV-lumen since EV-biogenesis occurs at low pH (5.5 to 6 for the MVB) (*46*). A variety of other inteins were engineered (*47*, *48*), and potential better candidates which increase both the loading and cleavage of precursors in EVs may further improve the VEDIC and VFIC systems.

Observing that VSV-G functions as both an EV-sorting domain and endosomal escape facilitator resulted in the design of the all-in-one VFIC system, which circumvents the limiting step of requiring the simultaneous co-transfection of multiple plasmids, which is often necessary in other technologies, and our VEDIC system. The VFIC approach led to a further increase in the efficiency of cargo delivery (Fig. 5, C and D and Fig. 8E). In addition, we speculate that VFIC/VEDIC have advantages over virus-like particles (VLPs) because they likely bear less nonspecifically packaged cellular components than the VLPs used by other groups since our systems are devoid of any viral assembly components (*27*, *49*). This may lead to fewer sides including inflammation induction upon therapeutic applications compared to VLPs.

Alongside the robust intracellular protein delivery observed *in vitro* in reporter cells, our systems achieved successful *in vivo* protein delivery in mice, both through local (IT and ICV) and systemic (IP) delivery. We profiled the cell types that internalized Cre-loaded engineered EVs in the brain after ICV injection, and in spleen and liver after IP injection in R26-LSL-tdTomato reporter mice. We found evidence that astrocytes and microglia were the cell types targeted by the engineered EVs in the brain, while T cells and macrophages primarily internalized the engineered EVs in spleen and liver. However, due to the broad tropism of VSV-G, other cell types may additionally internalize the engineered EVs. In addition, EV-isolation methods, EV source cell or route of EV-injection may all affect the *in vivo* distribution of EVs.

In order to better understand the effects of VSV-G on EV uptake and endosomal escape, we studied uptake of CD63-Intein-mNG engineered EVs using two mutants (K47Q and P127D, disrupting receptor-mediated EV uptake and endosomal escape respectively) of VSV-G (*36*, *37*). We found that VSV-G affects both endocytosis and membrane fusion that are necessary to induce the diffuse distribution of mNG, indicative of functional cytosolic protein delivery. These findings underscore the important role of endosomal escape for the cytosolic delivery of engineered EV cargo proteins in recipient cells. Moreover, these results demonstrate the value of CD63-Intein-mNG as a tool to study EV uptake and cargo delivery as a convenient assay to explore this process. These data also indicate that receptor-mediated endocytosis is the primary pathway responsible for the entry of VSV-G engineered EVs and subsequent cargo delivery into recipient cells.

One of the main concerns for the application of VSV-G is its potential immunogenicity *in vivo*, which would hamper repeated administration of engineered vesicles (*50*). A potential way to avoid repeated injections is to design systems that function as a single dose (*51*). As such, it is notable that we observed close to 80% *in vitro* gene editing efficiency by EV-mediated delivery of Cas9-RNPs in reporter cells, and significantly reduced PCSK9 expression by delivering a meganuclease. Aside from Cas9-RNPs, more advanced genome editing tools, such as base editors and prime editors, could be potentially delivered using the VEDIC and VFIC systems (*52*, *53*). We believe that there is great potential for such single-dose treatments to address a wide variety of genetic diseases if therapeutically relevant levels of gene editing can be achieved. EV-mediated delivery of Cas9 RNPs has a potential advantage over viral delivery of CRISPR/Cas expression systems, due to the lower off-target potential expected from the shorter exposure to such gene editing agents (*54–56*).

Furthermore, we applied engineered EVs for the treatment of LPS-induced systemic inflammation, demonstrating that therapeutic levels of intracellular protein delivery were achieved by our systems *in vivo*. These observations demonstrate the therapeutic potential of these approaches, which shows exciting promise for the potential development of treatments for a wide array of pathologies, such as lysosomal storage diseases (LSDs) and enzyme deficiencies (*57*, *58*). To summarize, the VEDIC and VFIC systems developed in this study allow for robust intracellular functional delivery of therapeutic proteins, both *in vitro* and *in vivo*. In addition, the high genome editing efficiency achieved by Cas9-RNP delivery implies that this strategy may lead to potential applications in the treatment of genetic diseases.

## Supporting information

Suppelmentary figures

## ACKNOWLEDGMENTS

We apologize to the researchers whose work is closely related to this study but cannot be cited because of limited space. The confocal imaging was performed at the Live Cell Imaging unit/Nikon Center of Excellence, BioNut, Karolinska Institutet. **Funding:** This study was supported by Cancerfonden (211762 Pj 01 H), H2020 (EXPERT), the Swedish foundation of Strategic Research (FormulaEx, SM19-0007), ERC CoG (DELIVER) and the Swedish Research Council (4–258/2021) got by S.E.-A.; by the Swedish Research Council (VR), Grant for half-time research position in a clinical environment (2021–02407), and the CIMED junior investigator grant got by J.Z.N.; by the Swedish Research Council (VR), Grant for half-time research position in a clinical environment (2022–02449), and the CIMED junior investigator grant (FoUI-976434) got by O.W.; and by Evox Therpeutics. **Author contributions:** Conceptualization: X.L., D.G., J.H., I.M., J.Z.N and S.E.-A. Methodology: X.L., J.X., Z.N., E.V.W., L.V.H., O.W., W.Z., R.J.W., R.H., D.R.M., J.B., G.Z., H.Z., S.R., A.Z., A.G., D.W.H., O.G.J., A.G.U., Y.Z., C.M.P., T.C.R., and R.E.V. Investigation: X.L., D.G., J.X., Z.N., E.V.W., L.V.H., O.W., W.Z., R.J.W., R.H., D.R.M., J.B., G.Z., H.Z., S.R., A.Z., A.G., D.W.H., O.G.J., A.G.U., Y.Z., C.M.P., T.C.R., Z.J.N., R.E.V., and S.E.-A. Visualization: X.L., D.G., J.X., Z.J.N., R.E.V., and S.E.-A. Funding acquisition: Z.J.N., O.W., and S.E.-A. Supervision: S.E.-A. Writing: X.L. Editing: X.L., D.H., O.G.J., T.C.R., Z.J.N., O.W., D.G., A.G.U., and S.E.-A.

## Competing interests

O.W., J.Z. N., D.G. A.G., M.J.A. W. and S.E.-A. are consultants and stakeholders in Evox Therapeutics Limited, Oxford, United Kingdom. A.D. F and J. H are employees of Evox Therapeutics Limited, Oxford, United Kingdom. All other authors declare no conflict of interest. **Data and materials availability:** All the data are available in the manuscript or in the supplementary information. Materials will be made available under a material transfer agreement upon request to S.E.-A. and Evox Therapeutics Limited, Oxford, United Kingdom.

## Materials and methods

### Construct generation

All the transgenes, except those purchased from Addgene and the human-derived fusogens, were ordered from IDT (Integrated DNA Technologies, USA). The transgenes were first cloned into the pLEX vector backbone using EcoRI and XhoI sites. The constructs used in this study were then generated from the ordered fragments through restriction enzyme digestion and subsequent self-ligation. VSV-G-Intein-Cre and VSVG-Foldon-Cre were generated from the VSV-G-Foldon-Intein-Cre construct using Kpn2I and MluI, respectively. For VSV-G-Cre, VSV-G-Intein-Cre was digested with MluI. CD63-Intein-SS was generated by digesting VSV-G-Foldon-Intein-SS and CD63-Intein-Cre with BamHI and Xhol, and then inserting SS into the resulting CD63-Intein vector. Similarly, to synthesize CD63-Intein-Cas9 and VSV-G-Foldon-Intein-Cas9 constructs, CD63-Intein-Cre and VSV-G-Foldon-Intein-Cre were digested with BamHI and Xhol, and Cas9 inserted into CD63-Intein and VSV-G-Foldon-Intein, respectively. The 40 human-derived fusogens were ordered from Twist (Twist Bioscience) and the vector used was pTwist CMV BetaGlobin.

### Cell culture

HEK-293T cells used to produce the functional EVs in this study were maintained in Dulbecco’s Modified Eagle Medium (DMEM) (high glucose) supplemented with 10% fetal bovine serum (FBS) (Gibco, USA) and 1% Antibiotic-Antimycotic (Gibco, USA). Cells were cultured at 37℃ in a humidified air atmosphere containing 5% CO_2_. The reporter cell lines (Hela-TL, T47D-TL, B16F10-TL, Raw264.7-TL, HEK-Blue-NF-ĸB, and HEK-SL) were cultured using the same medium and the same conditions as HEK-293T cells. THP-1-TL, K562-TL and MSC-TL cells were cultivated in Roswell Park Memorial Institute (RPMI) 1640 medium (high glucose) supplemented with 10% FBS and 1% Anti-anti.

### Plasmid transfection

HEK-293T cells were seeded into 15-cm dishes, with the numbers of dishes decided according to the amount of EVs to be used in indicated experiments. Polyethylenimine (PEI, Polysciences) was utilized for the transfection of plasmid/s according to the protocol provided by the manufacturer. The ratio of PEI to plasmid was 2:1 in this study. For single plasmid transfection, 30 µg plasmid was used for each plate. For co-transfection of 2 plasmids, 20 µg of each plasmid was used while for 3 plasmids, 15 µg of each plasmid was used.

### EV production

EVs were produced by transient transfection of the transgenes using polyethylenimine. To be specific, HEK-293T cells were seeded into 15-cm dishes at a density of 5 million cells per dish using complete DMEM medium. After 2 days, the cells were transfected with the transgenes and the medium changed to Opti-MEM (Gibco, USA) with 1% Anti-anti 6 h post-transfection. After 48 h, the conditioned medium (CM) was collected and centrifuged (700 × *g* for 5 min followed by 2,000 × *g* for 10 min). The supernatant was then filtered through a 0.22 µm filter system.

### EV isolation

Tangential flow filtration (TFF, MicroKross, 20 cm^2^, Spectrum labs) was used to isolate EVs from the filtered CM. Particles greater than the 300 kDa cutoff of the TFF were retained in the system and concentrated. These particles were further concentrated using Amicon Ultra-15 100 kDa (Millipore) spin filters, which were centrifuged at 4,000 × *g* for 30 min to several hours at 4°C, depending on the amount EVs in the samples. Lastly, the concentrated EVs were collected in maxirecovery 1.5 ml Eppendorf tubes (Axygene, USA) and quantified using Nanoparticle Tracking Analysis (NTA).

### Nanoparticle Tracking Analysis

EV samples were diluted with freshly 0.22 µm-filtered PBS before checking the particle sizes and concentrations using the NanoSight NS500 instrument. Five videos of more than 30 second durations each were taken at the camera level of 15 in light scatter mode. All the samples were analysed with the same setting using the NTA 2.3 software for Nanosight.

### MACSQuant flow cytometry

After the various traffic-light reporter cells were added with EVs at different time points, or after the reporter cells were co-cultured with EV-producing cells for 24 h, GFP expression was quantified using the MACSQuant Analyzer 10 flow cytometer (Miltenyi Biotec, Germany). Briefly, the cells in 96-well plates were washed with PBS once and trypsinized for 5 min at 37°C. The cells were then neutralized using cell medium supplemented with 10% FBS. After adding DAPI to check the cell viability, the cells were sampled by the MACSQuant analyzer using the settings of one specific reporter cell line for all the measurements. FlowJo (version 10.6.2) was used to calculate the percentage of GFP positive cells.

### Imaging flow cytometry

HEK-293T cells were either transfected with VSV-G-mNG construct only or co-transfected with VSV-G-mNG and CD63. Six hours post-transfection, the medium was changed to Opti-MEM medium (Gibco, USA) with 1% Anti-anti. After 2 days, the medium was harvested and centrifuged at 700 × *g* for 5 min, followed by 2,000 × *g* for 10 min and subsequently filtered through a 0.22 µm filter system. In a v-bottom 96-well plate, 25 µl of each sample was incubated with APC-labelled CD63 antibody (Miltenyi Biotec, Germany; 1 nM per well) overnight under dark conditions. The samples were then diluted 1000 times and transferred into an R-bottom 96-well plate. Amnis® CellStream instrument (Luminex, US) was utilized to evaluate the engineered EVs at single-vesicle level. The collected data was then analysed using FlowJo (version 10.6.2).

### EV-adding assay in reporter cells

The reporter cells used in this study were seeded into 96-well plates at the following densities: 1×10^4^ (Hela-TL), 2×10^4^ (T47D-TL), 1.5×10^4^ (B16F10-TL), 5×10^4^ (Raw264.7-TL), 5×10^4^ (THP-1-TL), 5×10^4^ (K562-TL), 1×10^4^ (MSC-TL), 8×10^3^ (HEK-SL) and 2×10^4^ (HEK-Blue-NF-ĸB) cells per well. The following day, appropriate doses of EVs were added directly onto each of the reporter cells. The doses of the EVs used and the time for incubation are indicated in each figure. GFP positivity was assessed by either fluorescence microscopy or MACSQuant flow cytometry. For HEK-Blue-NF-ĸB cells, the SS EVs were added directly into the wells for 48 hours and the stimulation (TNF-α, 10 ng/ml) added after another 6 hours. Luciferase signals from the cell lysate were evaluated using the GloMax 96 Microplate Luminometer (Promega, USA).

### Virus production

The transgenes were subcloned into our lenti-viral vector (Transfer plasmid, 22.5 µg/T175 flask) which were co-transfected with pCD/NL-BH (Helper plasmid, 22.5 µg/T175 flask) and pcoPE01 (Envelope plasmid, 7 µg/T175 flask) into HEK-293T cells and incubated overnight. The next morning, cell medium was changed to complete DMEM medium (with 10% FBS and 1% Antibiotic-Antimycotic) supplemented with sodium butyrate (Sigma-Aldrich). After 6 to 8 hours, the sodium butyrate containing DMEM medium was changed back to complete DMEM medium without additional chemicals. Nalgene Oak Ridge Centrifuge Tubes (Thermo Scientific) were used for harvesting viruses 22 to 24 hours after incubation with the normal medium. Briefly, the virus-particle-containing medium was collected and filtered using a 0.45 µm syringe filter (VWR), and then centrifuged at 25,000 × *g* for 90 min at 4°C. The supernatant was aspirated, and freshly prepared medium (IMDM with 20%FBS) was used to resuspend the virus pellets. The viruses were added directly into the target cells or stored at −80°C freezer for long-term storage.

### Stable reporter cell generation

B16F10, Raw264.7, THP-1, K562, and MSC cells were seeded in 6-well plates and the viruses were added into the cells the next day. Titration of viruses was performed using 3 doses: 2 µl, 10 µl, and 50 µl per well. After a one-day incubation of the viruses with target cells, the medium was changed back to normal complete medium (DMEM + 10% FBS + 1% Anti-anti for B16F10 and Raw264.7 cells; RPMI-1640 + 10% FBS + 1% Anti-anti for THP-1, K562, and MSC cells). Two days after virus transduction, the cells were trypsinized and resuspended in fresh medium. Resistance selection was performed by adding puromycin (2 µg/ml for B16F10 and MSC cells, 4 µg/ml for THP-1 and K562 cells, and 6 µg/ml for Raw264.7 cells). The cells were passaged under puromycin selection for approximately one week before the cells were utilized for downstream experiments.

### Direct co-culture of EV-producing cells with reporter cells

HEK-293T cells were seeded in a 6-well plate at a density of 0.5 million cells per well. The next day when the cells reached 60-70% confluence, constructs were transfected as indicated using Lipofectamine2000 (Invitrogen, USA) according to the manufacturer’s protocol. Six hours after transfection, the medium was changed to fresh medium (DMEM + 10%FBS + 1% Antibiotic-Antimycotic). Twenty-four hours after plasmid transfection, the cells were trypsinized, counted, and mixed with the corresponding reporter cells at ratios of 1:1 or 1:5, or other ratios as indicated in the figures (ratio=EV-producing cells: reporter cells) in a 96-well plate. After co-culturing for 24 h, the cells were trypsinized and measured using the MACSQuant flow cytometer to determine the percentage of GFP positive cells.

### IBIDI co-culture µ-slide assay

HEK-293T cells were seeded into a 6-well plate at a density of 5×10^4^ cells per well. The following day, constructs were transfected as indicated using Lipofectamine2000 (Invitrogen, USA) according to the manufacturer’s protocol. To avoid the toxicity of the Lipofectamine2000 on the HEK-293T cells, the medium was changed to fresh complete medium (DMEM + 10%FBS + 1% Anti-anti) after 6 h. The following day, the transfected cells were trypsinized and counted. The transfected cells (feeder cells or EV-producing cells) were seeded into the surrounding reservoirs of the ibidi µ-Slide while the recipient cells (traffic-light reporter cells) were added to the central reservoir, following cell numbers as indicated. The volume of medium used for each reservoir was 40 µl. Once the cells has attached, a further 400 µl of complete medium (DMEM + 10%FBS + 1% Anti-anti) was added into the slide slowly and carefully to immerse the walls between the central reservoir and the surrounding reservoirs such that cell-cell communication could be mediated by engineered EVs. Four days later, the GFP positive cells were measured using either a fluorescent microscope or the MACSQuant flow cytometer.

### Transwell co-culture assay

Similar to the IBIDI assay, HEK-293T cells were seeded into a 6-well plate for 24 h and then transfected with indicated constructs as indicated. The medium was changed to fresh complete medium after 6 h of transfection. One day after transfection, cells were trypsinized and counted, and the EV-producing cells were added to the top chamber of the transwell system (pore size=0.4 µm) while the reporter cells were seeded at the bottom. After 4 days of cell-cell communication by engineered EVs, GFP positivity was assessed by fluorescence microscopy or MACSQuant flow cytometry.

### Dynamic live imaging assay

Huh7 cells were plated 1 day before the experiment in a polymer-bottom cell culture plate (Ibidi, cat. no. 82426), with 5×10^4^ cells per well. 5×10^10^ EVs were added to the cells 3 hours prior imaging and Hoecsht dye for nucleus staining was added just before the live cell imaging. Confocal images were acquired on a Nikon C2 + confocal microscope equipped with an oil-immersion 60× objective with numerical aperture 1.4 (Nikon Instruments, Amsterdam, The Netherlands). The sample was excited and detected with appropriate excitation laser lines and emission filters and the fluorophores were imaged sequentially. The images were taken every hour and the whole process was 72 hours. The corresponding videos were generated by using the Nikon NIS-Elements Imaging Software.

### Confocal microscopy

Huh7 cells were seeded in polymer-bottom cell culture plates (Ibidi, cat. no. 82426) one day before the experiment, similar to the dynamic live imaging assay. EVs were added to the cells at doses as indicated one day after seeding. After adding EVs for 48 hours, Hoecsht dye was added before confocal microscopy imaging. The confocal images were taken the same way as described in dynamic live imaging assay. Image processing was performed by using Fiji software.

### Fluorescent microscopy

After addition of EVs, co-culture, IBIDI, and Transwell assays, the GFP positive cells were visualized under a fluorescent microscope. We chose the area for taking pictures randomly and set up the same parameters for the groups using one experiment. All the images were then processed with the same parameters using the Fiji software.

### Western blot analysis

Whole cell protein was isolated using RIPA buffer supplemented with a protease inhibitor cocktail, mixed with sample buffer (4×), and heated at 70℃ for 10 min. For EV samples, 1×10^10^ EVs were mixed with sample buffer (4×) and heated at 70℃ for 10 min. Samples were then loaded onto a NuPAGE 4-12% Bis-Tris Protein Gel (Thermo Scientific) and ran at 120 V for 2 h in NuPAGE MES SDS running buffer (Thermo Scientific). Proteins were transferred from the gel to the membrane using iBlot 2 Transfer Stacks (Thermo Scientific). The membrane was blocked with Intercept blocking buffer (LI-COR Biosciences) for 1 h at room temperature in a shaker after which it was incubated with primary antibodies overnight at 4℃. The membrane was washed with TBS-T buffer three times for 5 min each and incubated with corresponding secondary antibodies for 1 h at room temperature in a shaker. After washing with TBS-T buffer three times and with PBS once, the membrane was scanned using the Odyssey infrared imaging system (LI-COR).

### Intra-tumor injection model

C57BL/6 mice (5 weeks of age, 20 g body weight) were acclimated to their new surroundings at least one week before the experiment. B16F10-TL cells resuspended in PBS were inoculated subcutaneously into the mice at a density of 0.5 million cells per mouse. Ten days after inoculation when obvious tumors were formed, engineered EVs were injected directly into the tumors. The injected volume was 50 µL per mouse with 7.5× 10^10^ EVs. Four days after intra-tumor injection of EVs, the mice were sacrificed, and tumors were harvested and fixed in PFA. Tumor tissues were stained immunohistochemically for GFP expression while tissues in lysis buffer were homogenised using a tissue lyzer machine.

### LPS-induced inflammation model

C57BL/6 mice (5 weeks of age, 20 g body weight) were acclimated to their new surroundings at least one week before the experiment. Animals were injected (IP) with engineered EVs 4 h before IP injection of LPS (Sigma, USA) at the dose of 7.5 mg/Kg. Six hours after LPS induction, engineered EVs were IP injected once more to boost the intracellular delivery of the protein cargos by EVs. The survival rate and body weight of the mice with LPS induction were recorded for 2 days. Forty-eight hours after LPS induction, the mice were euthanized, and blood was sampled using heart puncture. The mice were sacrificed, and organs (liver, lung, spleen and kidney) were harvested and fixed with PFA. H&E (haematoxylin and eosin) staining was performed to check the extent of damage to the organs induced by LPS. The damage of the tissues was evaluated by a professional pathologist and histological scores of the tissues were given accordingly.

### IHC staining for melanoma tissues

Tissue sections were fixed at 65℃ for 1 hour before the slides were subjected to deparaffinization and rehydration as follows: Xylene for 20 min, 100% ethanol for 3 min twice, 95% ethanol for 3 min, 70% ethanol for 3 min, and then 50% ethanol for 3 min. Afterwards, the slides were rinsed in running cold tap water for 5 min followed by antigen retrieval using citrate buffer, pH 6.0 (Sigma). After antigen retrieval, the slides were washed with PBS three times for 5 min each and immersed in blocking buffer for 30 min at 37℃. The slides were then incubated with primary anti-GFP antibody (Abcam, ab290, 1:200 dilution) overnight after blocking. The following day, after washing the slides with PBS three times for 5 min each, the slides were incubated with secondary antibody Goat Anti Rabbit IgG H&L (Alexa Fluor 488) (Abcam, ab150077, 1:500 dilution) for 30 min at 37℃, followed by washing with PBS three times for 5 min each. The slides were mounted using ProLong Diamond Antifade Mountant with DAPI (Thermo Scientific) and sealed with nail polish. Images were taken using a confocal microscope (Nikon, Japan).

### IHC staining for tissues from Cre-LoxP R26-LSL-tdTomato reporter mice

Tissue sections (5 µm) were prepared from organs of ICV- and IP-injected R26-LSL-tdTomato reporter mice and then were deparaffinized in xylene and ethanol, boiled in citrate buffer for 20 min, and blocked with 5% goat serum in PBS-T (PBS containing 0.3% Triton X-100) solution for 1 h at room temperature. The sections were then stained with primary antibodies in blocking buffer at 4°C overnight. After washing with PBS, sections were stained with appropriate fluorophore-conjugated secondary antibodies in PBS or PBS containing 0.1% Triton X-100 for 1 to 2 h before washing and mounting. A Zeiss LSM780 confocal microscope or Zeiss Axioscan Z.1 was used for imaging.

### Statistics

Statistical tests for the biological replicates used in this study are reported in each figure legend. GraphPad software was utilized for the statistical analysis and the data presented as mean±SD. A two-tailed student’s *t*-test was used for the comparisons of two individual groups. One-way ANOVA or Two-way ANOVA multiple comparisons test was used for the analysis of the multiple groups. Log-rank (Mantel-Cox) test was used for the survival comparisons. Statistical significance was set up as * p < 0.05, ** p < 0.01; *** p < 0.001; **** p < 0.0001; ns: non-significant.

## Notes

### Summary of Updates

The initial submission missed one grant and we added this grant to the main manuscript.

## REFERENCES

1. W. A. Messersmith, D. J. Ahnen, edi t or i a l s Targeting EGFR in Colorectal Cancer, 1834–1836 (2022).

2. M. Reck, T. Wehler, F. Orlandi, N. Nogami, C. Barone, D. Moro-Sibilot, M. Shtivelband, J. L. G. Larriba, J. Rothenstein, M. Früh, W. Yu, Y. Deng, S. Coleman, G. Shankar, H. Patel, C. Kelsch, A. Lee, E. Piault, M. A. Socinski, Safety and patient-reported outcomes of atezolizumab plus chemotherapy with or without bevacizumab versus bevacizumab plus chemotherapy in non-small-cell lung cancer. J. Clin. Oncol. 38, 2530–2542 (2020).

3. M. P. Stewart, A. Sharei, X. Ding, G. Sahay, R. Langer, K. F. Jensen, In vitro and ex vivo strategies for intracellular delivery. Nature. 538, 183–192 (2016).

4. R. Goswami, T. Jeon, H. Nagaraj, S. Zhai, V. M. Rotello, Accessing Intracellular Targets through Nanocarrier-Mediated Cytosolic Protein Delivery. Trends Pharmacol. Sci. 41, 743–754 (2020).

5. D. S. D’Astolfo, R. J. Pagliero, A. Pras, W. R. Karthaus, H. Clevers, V. Prasad, R. J. Lebbink, H. Rehmann, N. Geijsen, Efficient intracellular delivery of native proteins. Cell. 161, 674–690 (2015).

6. J. Fu, C. Yu, L. Li, S. Q. Yao, Intracellular Delivery of Functional Proteins and Native Drugs by Cell-Penetrating Poly(disulfide)s. J. Am. Chem. Soc. 137, 12153–12160 (2015).

7. A. Steinauer, J. R. LaRochelle, S. L. Knox, R. F. Wissner, S. Berry, A. Schepartz, HOPS-dependent endosomal fusion required for efficient cytosolic delivery of therapeutic peptides and small proteins. Proc. Natl. Acad. Sci. U. S. A. 116, 512–521 (2019).

8. J. S. Wadia, R. V. Stan, S. F. Dowdy, Transducible TAT-HA fusogenic peptide enhances escape of TAT-fusion proteins after lipid raft macropinocytosis. Nat. Med. 10, 310–315 (2004).

9. N. Boehnke, J. P. Straehla, H. C. Safford, M. Kocak, M. G. Rees, M. Ronan, D. Rosenberg, C. H. Adelmann, R. R. Chivukula, N. Nabar, A. G. Berger, N. G. Lamson, J. H. Cheah, H. Li, J. A. Roth, A. N. Koehler, P. T. Hammond, Massively parallel pooled screening reveals genomic determinants of nanoparticle delivery. Science (80-.). 377 (2022), doi:10.1126/science.abm5551.

10. P. Chakravarty, W. Qian, M. A. El-Sayed, M. R. Prausnitz, Delivery of molecules into cells using carbon nanoparticles activated by femtosecond laser pulses. Nat. Nanotechnol. 5, 607–611 (2010).

11. B. Liu, M. Ianosi-Irimie, S. Thayumanavan, Reversible Click Chemistry for Ultrafast and Quantitative Formation of Protein-Polymer Nanoassembly and Intracellular Protein Delivery. ACS Nano. 13, 9408–9420 (2019).

12. M. Yan, J. Du, Z. Gu, M. Liang, Y. Hu, W. Zhang, S. Priceman, L. Wu, Z. H. Zhou, Z. Liu, T. Segura, Y. Tang, Y. Lu, A novel intracellular protein delivery platform based on single-protein nanocapsules. Nat. Nanotechnol. 5, 48–53 (2010).

13. J. A. Zuris, D. B. Thompson, Y. Shu, J. P. Guilinger, J. L. Bessen, J. H. Hu, M. L. Maeder, J. K. Joung, Z. Y. Chen, D. R. Liu, Cationic lipid-mediated delivery of proteins enables efficient protein-based genome editing in vitro and in vivo. Nat. Biotechnol. 33, 73–80 (2015).

14. S. Kamerkar, V. S. Lebleu, H. Sugimoto, S. Yang, C. F. Ruivo, S. A. Melo, J. J. Lee, R. Kalluri, Exosomes facilitate therapeutic targeting of oncogenic KRAS in pancreatic cancer. Nature. 546, 498–503 (2017).

15. I. K. Herrmann, M. J. A. Wood, G. Fuhrmann, Extracellular vesicles as a next-generation drug delivery platform. Nat. Nanotechnol. 16, 748–759 (2021).

16. M. Mehanny, C. M. Lehr, G. Fuhrmann, Extracellular vesicles as antigen carriers for novel vaccination avenues. Adv. Drug Deliv. Rev. 173, 164–180 (2021).

17. X. Liang, Z. Niu, V. Galli, N. Howe, Y. Zhao, O. P. B. Wiklander, W. Zheng, R. J. Wiklander, G. Corso, C. Davies, J. Hean, E. Kyriakopoulou, D. R. Mamand, R. Amin, J. Z. Nordin, D. Gupta, S. E. L. Andaloussi, Extracellular vesicles engineered to bind albumin demonstrate extended circulation time and lymph node accumulation in mouse models. J. Extracell. vesicles. 11 (2022), doi:10.1002/jev2.12248.

18. S. Banskota, A. Raguram, S. Suh, S. W. Du, J. R. Davis, E. H. Choi, X. Wang, S. C. Nielsen, G. A. Newby, P. B. Randolph, M. J. Osborn, K. Musunuru, K. Palczewski, D. R. Liu, Engineered virus-like particles for efficient in vivo delivery of therapeutic proteins. Cell. 185, 250–265.e16 (2022).

19. M. Somiya, S. Kuroda, Reporter gene assay for membrane fusion of extracellular vesicles. J. Extracell. vesicles. 10 (2021), doi:10.1002/jev2.12171.

20. N. Yim, S. W. Ryu, K. Choi, K. R. Lee, S. Lee, H. Choi, J. Kim, M. R. Shaker, W. Sun, J. H. Park, D. Kim, W. Do Heo, C. Choi, Exosome engineering for efficient intracellular delivery of soluble proteins using optically reversible protein-protein interaction module. Nat. Commun. 7, 1–9 (2016).

21. D. W. Wood, W. Wu, G. Belfort, V. Derbyshire, M. Belfort, A genetic system yields self-cleaving inteins for bioseparations. Nat. Biotechnol. 17, 889–892 (1999).

22. L. A. Mulcahy, R. C. Pink, D. R. F. Carter, Routes and mechanisms of extracellular vesicle uptake. J. Extracell. vesicles. 3, 1–14 (2014).

23. J. Gilleron, W. Querbes, A. Zeigerer, A. Borodovsky, G. Marsico, U. Schubert, K. Manygoats, S. Seifert, C. Andree, M. Stöter, H. Epstein-Barash, L. Zhang, V. Koteliansky, K. Fitzgerald, E. Fava, M. Bickle, Y. Kalaidzidis, A. Akinc, M. Maier, M. Zerial, Image-based analysis of lipid nanoparticle-mediated siRNA delivery, intracellular trafficking and endosomal escape. Nat. Biotechnol. 31, 638–646 (2013).

24. X. Zhang, Q. Xu, Z. Zi, Z. Liu, C. Wan, L. Crisman, J. Shen, X. Liu, Programmable Extracellular Vesicles for Macromolecule Delivery and Genome Modifications. Dev. Cell. 55, 784–801.e9 (2020).

25. M. Albanese, Y. F. A. Chen, C. Hüls, K. Gärtner, T. Tagawa, E. Mejias-Perez, O. T. Keppler, C. Göbel, R. Zeidler, M. Shein, A. K. Schütz, W. Hammerschmidt, MicroRNAs are minor constituents of extracellular vesicles that are rarely delivered to target cells (2021), vol. 17.

26. A. Zomer, S. C. Steenbeek, C. Maynard, J. Van Rheenen, Studying extracellular vesicle transfer by a Cre-loxP method. Nat. Protoc. 11, 87–101 (2016).

27. P. E. Mangeot, S. Dollet, M. Girard, C. Ciancia, S. Joly, M. Peschanski, V. Lotteau, Protein transfer into human cells by vsv-g-induced nanovesicles. Mol. Ther. 19, 1656– 1666 (2011).

28. A. Görgens, M. Bremer, R. Ferrer-tur, F. Murke, T. Tertel, P. A. Horn, Optimisation of imaging flow cytometry for the analysis of single extracellular vesicles by using fluorescence-tagged vesicles as biological reference material. 8 (2019), doi:10.1080/20013078.2019.1587567.

29. A. Zomer, C. Maynard, F. J. Verweij, A. Kamermans, R. Schäfer, E. Beerling, R. M. Schiffelers, E. De Wit, J. Berenguer, S. I. J. Ellenbroek, T. Wurdinger, D. M. Pegtel, J. Van Rheenen, In vivo imaging reveals extracellular vesicle-mediated phenocopying of metastatic behavior. Cell. 161, 1046–1057 (2015).

30. A. B. Vogel, I. Kanevsky, Y. Che, K. A. Swanson, A. Muik, M. Vormehr, L. M. Kranz, K. C. Walzer, S. Hein, A. Güler, J. Loschko, M. S. Maddur, A. Ota-Setlik, K. Tompkins, J. Cole, B. G. Lui, T. Ziegenhals, A. Plaschke, D. Eisel, S. C. Dany, S. Fesser, S. Erbar, F. Bates, D. Schneider, B. Jesionek, B. Sänger, A. K. Wallisch, Y. Feuchter, H. Junginger, S. A. Krumm, A. P. Heinen, P. Adams-Quack, J. Schlereth, S. Schille, C. Kröner, R. de la Caridad Güimil Garcia, T. Hiller, L. Fischer, R. S. Sellers, S. Choudhary, O. Gonzalez, F. Vascotto, M. R. Gutman, J. A. Fontenot, S. Hall-Ursone, K. Brasky, M. C. Griffor, S. Han, A. A. H. Su, J. A. Lees, N. L. Nedoma, E. H. Mashalidis, P. V. Sahasrabudhe, C. Y. Tan, D. Pavliakova, G. Singh, C. Fontes-Garfias, M. Pride, I. L. Scully, T. Ciolino, J. Obregon, M. Gazi, R. Carrion, K. J. Alfson, W. V. Kalina, D. Kaushal, P. Y. Shi, T. Klamp, C. Rosenbaum, A. N. Kuhn, Ö. Türeci, P. R. Dormitzer, K. U. Jansen, U. Sahin, BNT162b vaccines protect rhesus macaques from SARS-CoV-2. Nature. 592, 283–289 (2021).

31. K. Papanikolopoulou, V. Forge, P. Goeltz, A. Mitraki, Formation of Highly Stable Chimeric Trimers by Fusion of an Adenovirus Fiber Shaft Fragment with the Foldon Domain of Bacteriophage T4 Fibritin. J. Biol. Chem. 279, 8991–8998 (2004).

32. J. Callanan, S. R. Stockdale, A. Shkoporov, L. A. Draper, R. P. Ross, C. Hill, Expansion of known ssRNA phage genomes: From tens to over a thousand. Sci. Adv. 6 (2020), doi:10.1126/sciadv.aay5981.

33. S. Chong, G. E. Montello, A. Zhang, E. J. Cantor, W. Liao, M. Xu, J. Benner, Utilizing the C-terminal cleavage activity of a protein splicing element to purify recombinant proteins in a single chromatographic step. 26, 5109–5115 (1998).

34. P. Van Roey, B. Pereira, Z. Li, K. Hiraga, M. Belfort, V. Derbyshire, T. H. P. Isermann, Crystallographic and Mutational Studies of Mycobacterium tuberculosis recA Mini-inteins Suggest a Pivotal Role for a Highly Conserved Aspartate Residue, 162–173 (2007).

35. J. Votteler, C. Ogohara, S. Yi, Y. Hsia, U. Nattermann, D. M. Belnap, N. P. King, W. I. Sundquist, Designed proteins induce the formation of nanocage-containing extracellular vesicles. Nature. 540, 292–295 (2016).

36. B. L. Fredericksen, M. A. Whitt, Vesicular stomatitis virus glycoprotein mutations that affect membrane fusion activity and abolish virus infectivity. J. Virol. 69, 1435–1443 (1995).

37. J. Nikolic, L. Belot, H. Raux, P. Legrand, Y. Gaudin, A. A. Albertini, Structural basis for the recognition of LDL-receptor family members by VSV glycoprotein. Nat. Commun. 9, 1–12 (2018).

38. T. Wan, J. Zhong, Q. Pan, T. Zhou, Y. Ping, X. Liu, Exosome-mediated delivery of Cas9 ribonucleoprotein complexes for tissue-specific gene therapy of liver diseases. 2017, 1–14 (2022).

39. O. G. de Jong, D. E. Murphy, I. Mäger, E. Willms, A. Garcia-Guerra, J. J. Gitz-Francois, J. Lefferts, D. Gupta, S. C. Steenbeek, J. van Rheenen, S. El Andaloussi, R. M. Schiffelers, M. J. A. Wood, P. Vader, A CRISPR-Cas9-based reporter system for single-cell detection of extracellular vesicle-mediated functional transfer of RNA. Nat. Commun. 11, 1–13 (2020).

40. L. Wang, J. Smith, C. Breton, P. Clark, J. Zhang, L. Ying, Y. Che, J. Lape, P. Bell, R. Calcedo, E. L. Buza, A. Saveliev, V. V. Bartsevich, Z. He, J. White, M. Li, D. Jantz, J M. Wilson, Meganuclease targeting of PCSK9 in macaque liver leads to stable reduction in serum cholesterol. Nat. Biotechnol. 36, 717–725 (2018).

41. J. Schreiber, P. A. Efron, J. E. Park, L. L. Moldawer, A. Barbul, Adenoviral gene transfer of an NF-κB super-repressor increases collagen deposition in rodent cutaneous wound healing. Surgery. 138, 940–946 (2005).

42. T. Liu, L. Zhang, D. Joo, S. C. Sun, NF-κB signaling in inflammation. Signal Transduct. Target. Ther. 2 (2017), doi:10.1038/sigtrans.2017.23.

43. S. Kim, S. A. Lee, H. Yoon, M. Y. Kim, J. K. Yoo, S. H. Ahn, C. H. Park, J. Park, B. Y. Nam, J. T. Park, S. H. Han, S. W. Kang, N. H. Kim, H. S. Kim, D. Han, J. I. Yook, C. Choi, T. H. Yoo, Exosome-based delivery of super-repressor IκBα ameliorates kidney ischemia-reperfusion injury. Kidney Int. 100, 570–584 (2021).

44. A. S. Aranko, A. Wlodawer, Nature’s recipe for splitting inteins. 27, 263–271 (2014).

45. D. W. Wood, J. A. Camarero, Intein Applications: From Protein Purification and Labeling to Metabolic Control. J. Biol. Chem. 289, 14512–14519 (2014).

46. M. P. Bebelman, P. Bun, S. Huveneers, G. Van Niel, D. M. Pegtel, F. J. Verweij, Real-time imaging of multivesicular body – plasma membrane fusion to quantify exosome release from single cells. Nat. Protoc. 15 (2020), doi:10.1038/s41596-019-0245-4.

47. D. S. Kelley, C. W. Lennon, Z. Li, M. R. Miller, N. K. Banavali, H. Li, M. Belfort, stress sensor. Nat. Commun., doi:10.1038/s41467-018-06554-x.

48. B. Dassa, N. London, B. L. Stoddard, O. Schueler-furman, Fractured genes: a novel genomic arrangement involving new split inteins and a new homing endonuclease family. 37, 2560–2573 (2009).

49. S. Johnson, J. X. Wheeler, R. Thorpe, M. Collins, Y. Takeuchi, Y. Zhao, Mass spectrometry analysis reveals differences in the host cell protein species found in pseudotyped lentiviral vectors. Biologicals. 52, 59–66 (2018).

50. S. Kuate, C. Stahl-Hennig, H. Stoiber, G. Nchinda, A. Floto, M. Franz, U. Sauermann, S. Bredl, L. Deml, R. Ignatius, S. Norley, P. Racz, K. Tenner-Racz, R. M. Steinman, R. Wagner, K. Überla, Immunogenicity and efficacy of immunodeficiency virus-like particles pseudotyped with the G protein of vesicular stomatitis virus. Virology. 351, 133–144 (2006).

51. D. Rosenblum, A. Gutkin, R. Kedmi, S. Ramishetti, N. Veiga, A. M. Jacobi, M. S. Schubert, D. Friedmann-Morvinski, Z. R. Cohen, M. A. Behlke, J. Lieberman, D. Peer, CRISPR-Cas9 genome editing using targeted lipid nanoparticles for cancer therapy. Sci. Adv. 6 (2020), doi:10.1126/sciadv.abc9450.

52. L. W. Koblan, M. R. Erdos, C. Wilson, W. A. Cabral, J. M. Levy, Z. M. Xiong, U. L. Tavarez, L. M. Davison, Y. G. Gete, X. Mao, G. A. Newby, S. P. Doherty, N. Narisu, Q. Sheng, C. Krilow, C. Y. Lin, L. B. Gordon, K. Cao, F. S. Collins, J. D. Brown, D. R. Liu, In vivo base editing rescues Hutchinson–Gilford progeria syndrome in mice. Nature. 589, 608–614 (2021).

53. A. V. Anzalone, P. B. Randolph, J. R. Davis, A. A. Sousa, L. W. Koblan, J. M. Levy, P. J. Chen, C. Wilson, G. A. Newby, A. Raguram, D. R. Liu, Search-and-replace genome editing without double-strand breaks or donor DNA. Nature. 576, 149–157 (2019).

54. G. A. Newby, J. S. Yen, K. J. Woodard, T. Mayuranathan, C. R. Lazzarotto, Y. Li, H. Sheppard-Tillman, S. N. Porter, Y. Yao, K. Mayberry, K. A. Everette, Y. Jang, C. J. Podracky, E. Thaman, C. Lechauve, A. Sharma, J. M. Henderson, M. F. Richter, K. T. Zhao, S. M. Miller, T. Wang, L. W. Koblan, A. P. McCaffrey, J. F. Tisdale, T. A. Kalfa, S. M. Pruett-Miller, S. Q. Tsai, M. J. Weiss, D. R. Liu, Base editing of haematopoietic stem cells rescues sickle cell disease in mice. Nature. 595, 295–302 (2021).

55. Y. Zhang, Z. Liang, Y. Zong, Y. Wang, J. Liu, K. Chen, J. L. Qiu, C. Gao, Efficient and transgene-free genome editing in wheat through transient expression of CRISPR/Cas9 DNA or RNA. Nat. Commun. 7, 1–8 (2016).

56. J. P. Han, M. J. Kim, B. S. Choi, J. H. Lee, G. S. Lee, M. Jeong, Y. Lee, E. A. Kim, H. K. Oh, N. Go, H. Lee, K. J. Lee, U. G. Kim, J. Y. Lee, S. Kim, J. Chang, H. Lee, D. W. Song, S. C. Yeom, In vivo delivery of CRISPR-Cas9 using lipid nanoparticles enables antithrombin gene editing for sustainable hemophilia A and B therapy. Sci. Adv. 8, 1–11 (2022).

57. F. M. Platt, A. d’Azzo, B. L. Davidson, E. F. Neufeld, C. J. Tifft, Lysosomal storage diseases. Nat. Rev. Dis. Prim. 4 (2018), doi:10.1038/s41572-018-0025-4.

58. B. K. Burton, S. Kar, P. Kirkpatrick, Sapropterin. Nat. Rev. Drug Discov. 7, 199–200 (2008).

