## Supplementary material for "Multimodal engineering of extracellular vesicles for efficient intracellular protein delivery": Suppelmentary figures

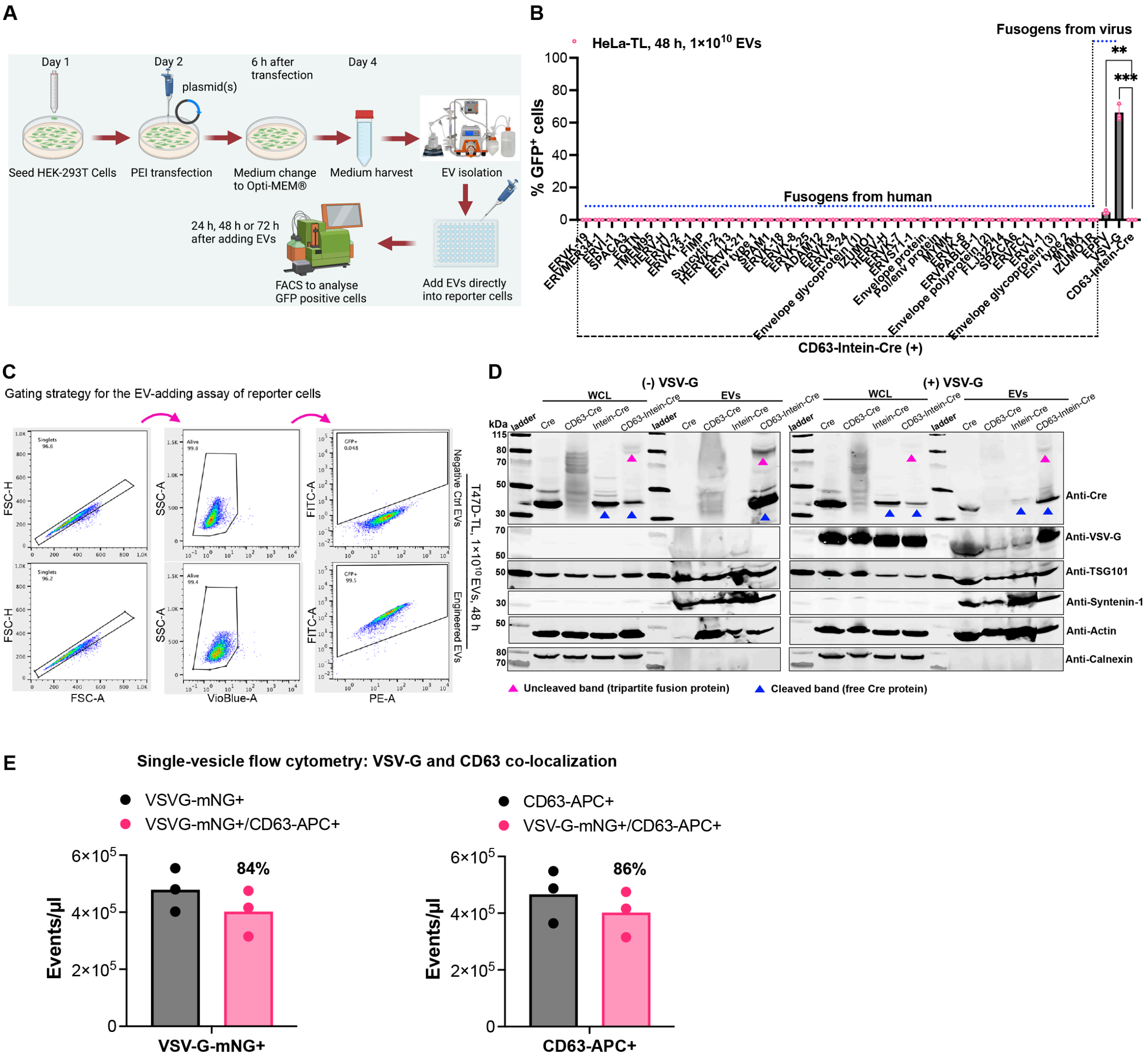


**Fig. S1. Characterization of fusogens on EV cargo loading and delivery** (**A**) A workflow schematic for the evaluation of EV-mediated Cre delivery to Cre recombinase fluorescent Traffic Light (TL) reporter cells and the subsequent analysis of GFP positive cells in recipient cells after adding isolated EVs. (**B**) Fusogen screen in HeLa-TL cells after a two-day incubation with EVs. (**C**) Example of the applied flow cytometry gating strategy for the analysis of GFP positive cells to evaluate Cre recombination after adding isolated EVs to fluorescent TL reporter cells. (**D**) Protein expression of various engineered Cre fusion constructs in whole cell lysates (WCL) and isolated EVs derived from HEK293T cells evaluated by western blot analysis. Lysates from $5\times10^5$ EV-producing cells and $1\times10^{10}$ engineered vesicles were used for the assay. TSG101, syntenin-1 and β-actin were used as EV markers and Calnexin was used as a cellular organelle marker (endoplasmic reticulum) and should be absent for EV samples. (**E**) Co-localization of VSV-G and CD63 in isolated vesicles measured by single-vesicle flow cytometry. One-way ANOVA multiple comparisons test was used for analysis of (B). Data are shown as mean+SD, ** $p < 0.01$; *** $p < 0.001$.

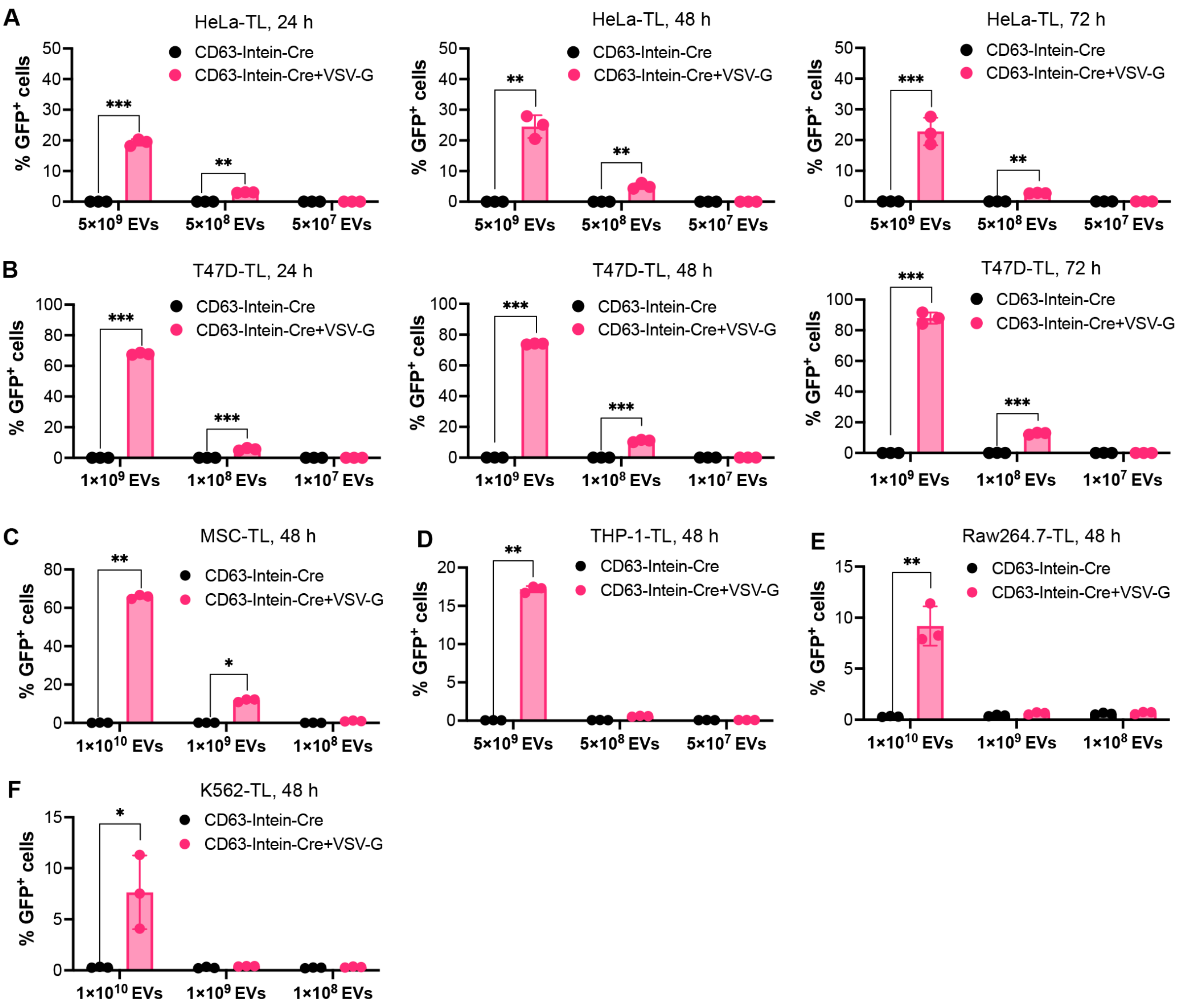


**Fig. S2. Flow cytometry analysis of recombination in reporter cells by EV-mediated Cre delivery using the VEDIC system.** (**A** and **B**) Percentage of GFP positive cells in HeLa-TL and T47D-TL reporter cells respectively after adding different doses of isolated engineered EVs for 1 to 3 days. (**C**-**F**) Percentage of recombined GFP positive cells mediated by intracellular Cre delivery through different doses of engineered EVs for 48 hours (h) in hard-to-transfect cells (C: MSC-TL; D: THP-1-TL; E: Raw264.7-TL; F: K562-TL). Two-way ANOVA multiple comparisons test was used for analysis of (A) to (F). Data are shown as means+SD, * $p < 0.05$; ** $p < 0.01$; *** $p < 0.001$.

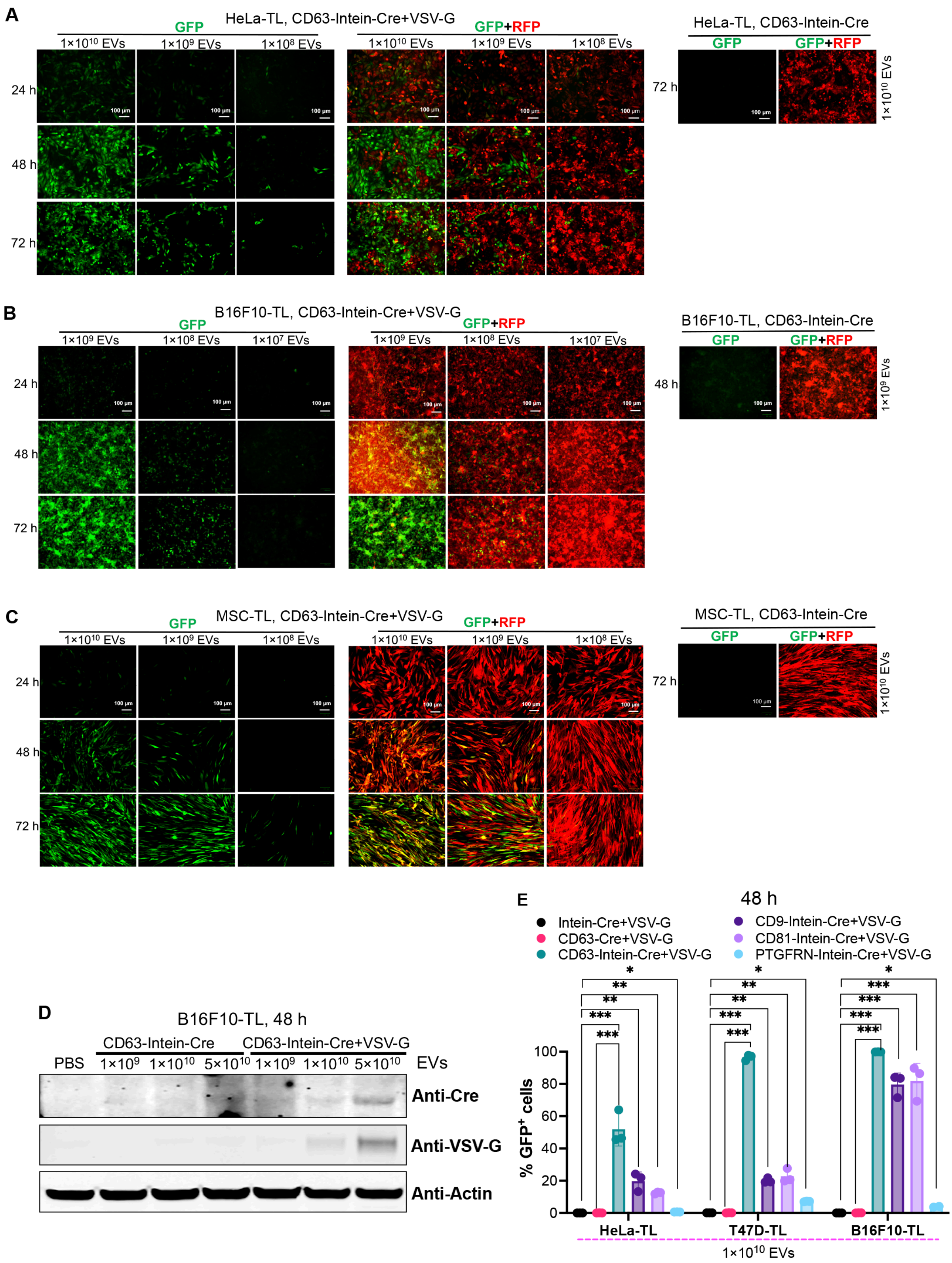


**Fig. S3. Fluorescent microscopy Analysis of EV-mediated Cre delivery and test of different fusogenic proteins and EV-sorting domains for VEDIC system.** (**A** to **C**) Fluorescent microscopy analysis of Cre-mediated HeLa-TL (A), T47D-TL (B), and B16F10-TL (C) reporter cell activation after adding indicated doses of EVs for indicated time of incubation. Scale bar, 100 μm. (**D**) Cre and VSV-G proteins measured by WB in B16F10-TL cells after adding the indicated doses of engineered EVs in 24-well plates. WB: western blot. (**E**) Recombination in reporter cells mediated by EVs derived from engineered cells using different EV-sorting domains. Two-way ANOVA multiple comparisons test was used for analysis of (E). Data are shown as mean±SD, * $p < 0.05$; ** $p < 0.01$; *** $p < 0.001$.

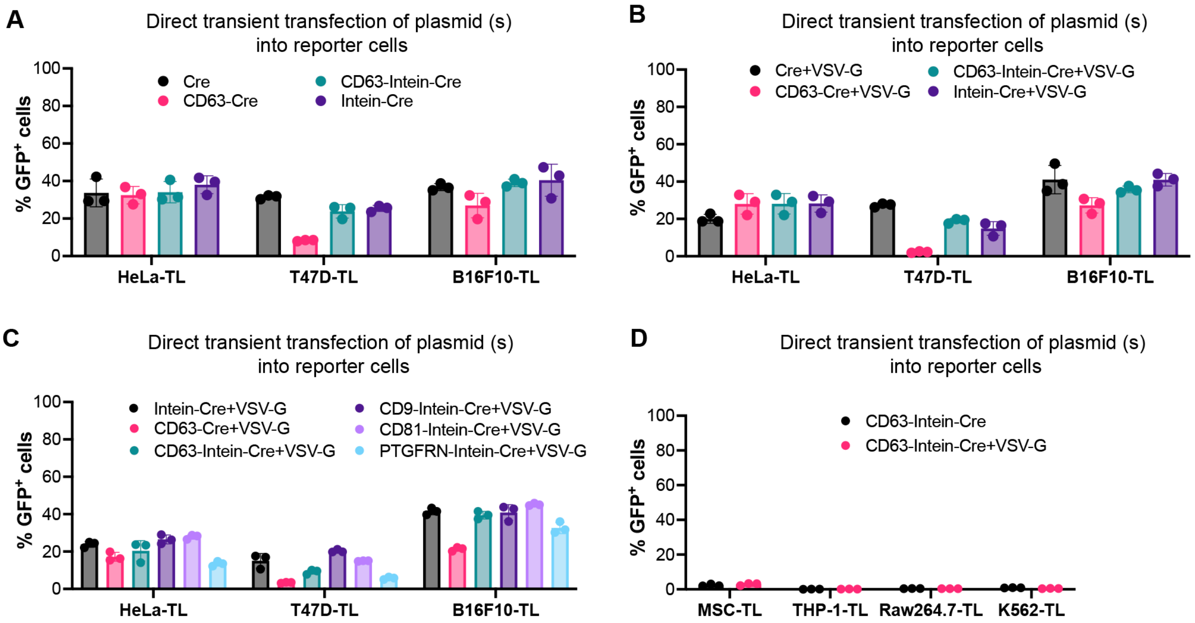


**Fig. S4. Cre-mediated recombination in reporter cells follwing transient transfection of indicated plasmids evaluated by flow cytometry.** (**A-D**) Percentage of GFP positive cells after transient transfection of the indicated plasmids into reporter cells after 48 h.

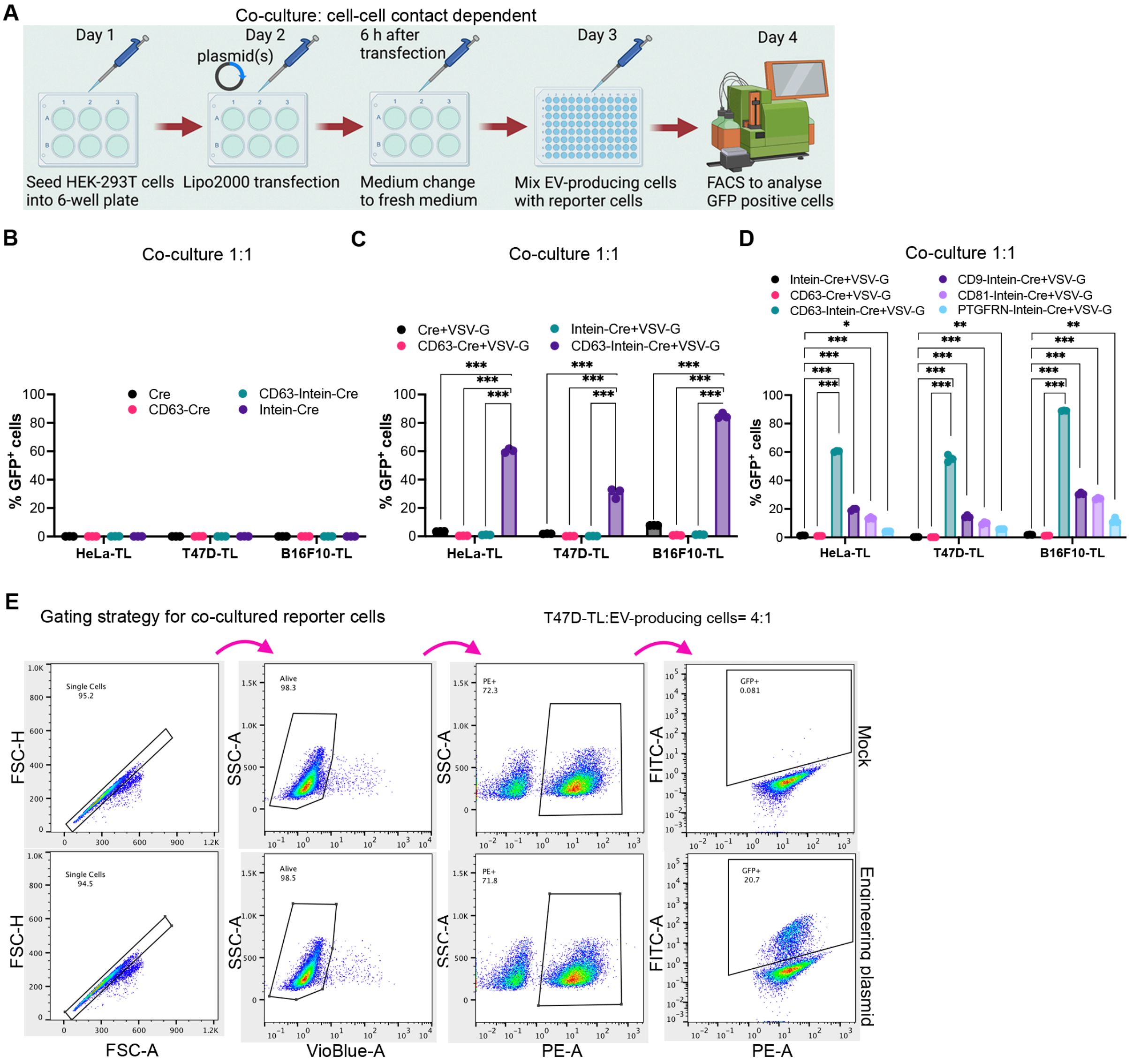


**Fig. S5. Flow cytometry analysis of Cre transfer in direct co-culture conditions by the VEDIC system.** (**A**) Brief workflow for the direct co-culture assay of EV-producing cells and reporter cells to show EV-mediated Cre delivery. (**B**) Co-culture assay for evaluation of Cre transfer without expression of VSV-G. (**C**) Analysis of Cre-mediated recombination in cells after a co-culture assay with expression of VSV-G. (**D**) Comparison of Cre transfer efficiency using different EV loading domains in a direct co-culture assay. (**E**) Gating strategy utilized for the analysis of Cre recombination, as indicated by the percentage of GFP positive cells, in co-culture assay. Two-way ANOVA multiple comparisons test was used for analysis of (C) and (D). Data were shown as means+SD, * $p < 0.05$; ** $p < 0.01$; *** $p < 0.001$.

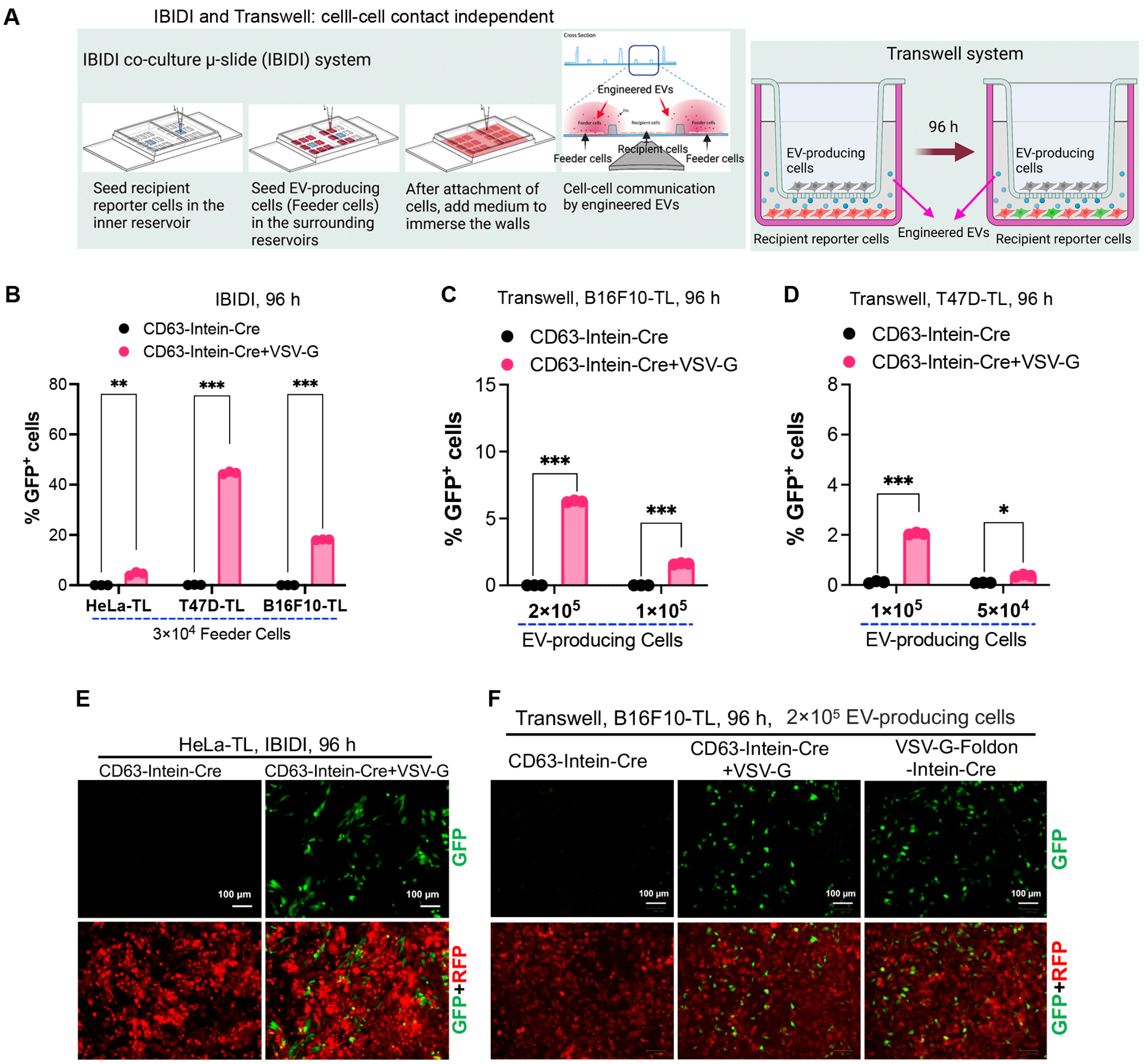


**Fig. S6. Recombination in reporter cells by VEDIC system in co-culture experiments without cell contact.** (**A**) Schematic graph to show the principle and workflow of IBIDI co-culture μ-slide (IBIDI) and Transwell co-culture assays demonstrating cell-cell contact independent Cre delivery from donor cells to recipient cells. The pore size for Transwell assay was 0.4 μm. (**B**) Flow cytometry analysis of Cre delivery mediated by engineered EVs in a contact-independent co-culture assay using IBIDI assay. $3\times10^4$ EV-producing cells and $4\times10^4$ reporter cells were seeded into the surrounding reservoirs and central reservoir respectively and the analysis was performed after 4 days. (**C** and **D**) Flow cytometry analysis of Engineered EV-mediated Cre delivery from donor cells to B16F10-TL and T47D-TL reporter cells in a co-culture Transwell assay in 24-well plates. Indicated numbers of EV-producing cells were seeded into the upper chamber and the numbers of B16F10-TL and T47D-TL reporter cells in the lower chamber were $5\times10^4$ and $8\times10^4$ respectively in 24-well plate. Flow cytometry analysis was performed after 96 hours. (**E**) Representative fluorescence microscopy images showing Cre recombinase-induced GFP expression in HeLa-TL cells by IBIDI assay. Scale bar, 100 μm. (**F**) Fluorescence microscopy images demonstrating GFP positive cells in B16F10-TL after a Transwell assay in 24-well plates. Scale bar, 100 μm. Two-way multiple comparisons ANOVA test was used for analysis of (B) to (D). Data are shown as means+SD, * $p < 0.05$; ** $p < 0.01$; *** $p < 0.001$.

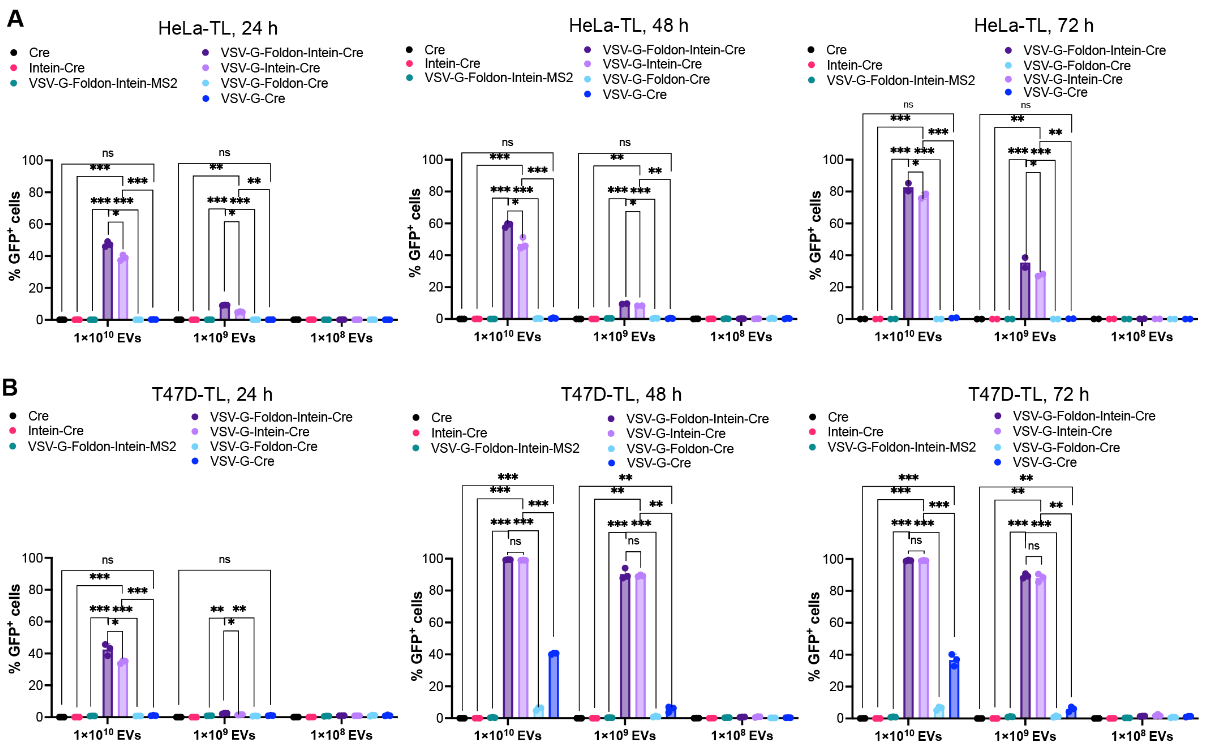


**Fig. S7. Flow cytometry analysis of Cre recombination in reporter cells by VFIC system.** (**A**) Percentage of GFP positive cells in HeLa-TL cells after adding engineered EVs. Indicated doses of EVs were added and the indicated time and analyzed at the various indicated timepoints. (**B**) Percentage of recombined cells mediated by Cre from adding engineered EVs in T47D-TL cells after adding indicated doses of EVs analyzed at the various indicated timepoints. Two-way ANOVA multiple comparisons test was used for analysis of (A) and (B). Data are shown as means+SD, * $p < 0.05$; ** $p < 0.01$; *** $p < 0.001$, ns: non-significant.

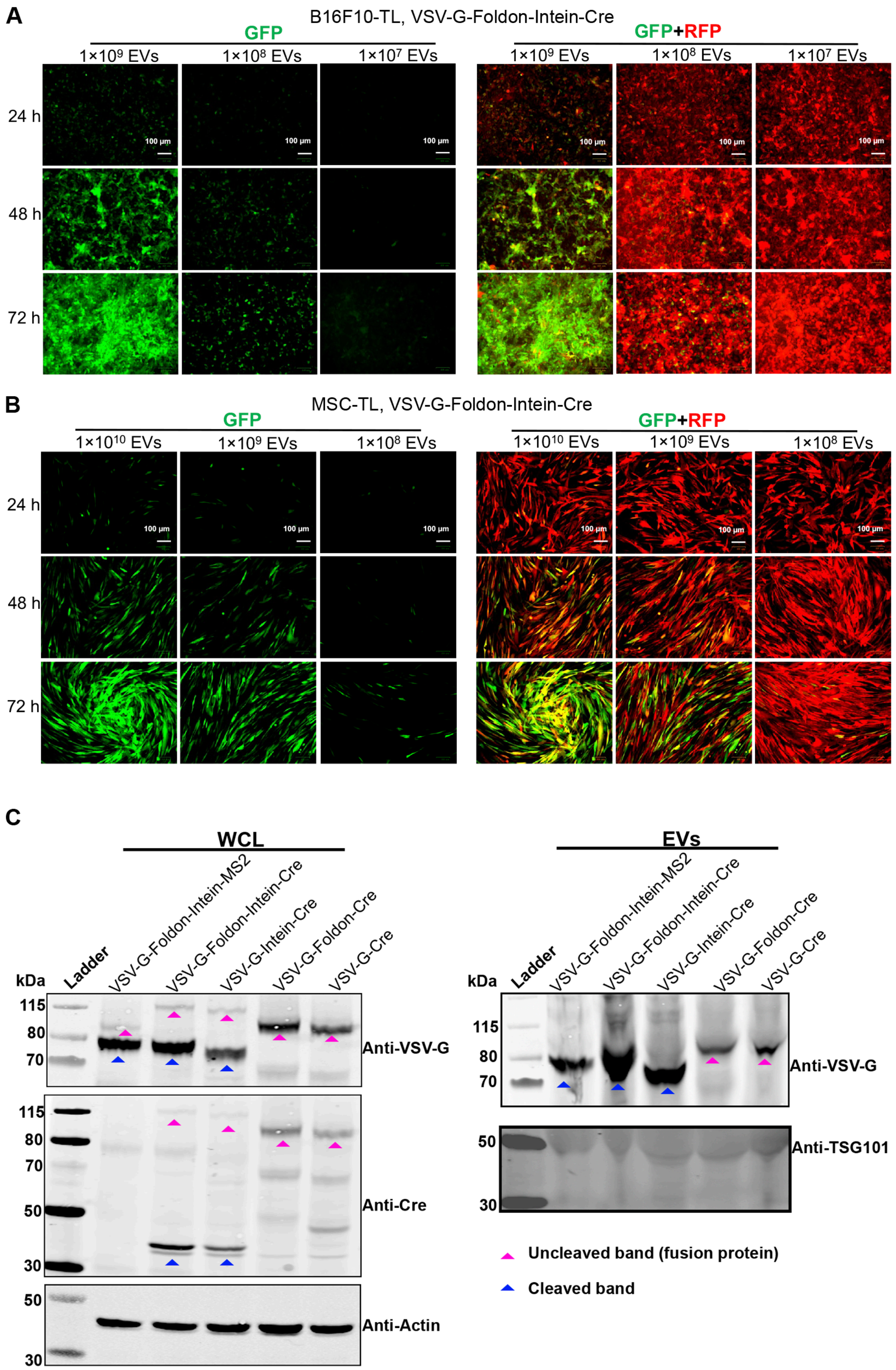
A
B16F10-TL, VSV-G-Foldon-Intein-Cre
GFP
GFP+RFP
1×10^9 EVs
1×10^8 EVs
1×10^7 EVs
24 h
48 h
72 h
100 μm
B
MSC-TL, VSV-G-Foldon-Intein-Cre
1×10^10 EVs
C
WCL
EVs
kDa
Ladder
VSV-G-Foldon-Intein-MS2
VSV-G-Foldon-Intein-Cre
VSV-G-Intein-Cre
VSV-G-Foldon-Cre
VSV-G-Cre
115
80
70
50
30
Anti-VSV-G
Anti-Cre
Anti-Actin
Anti-TSG101
Uncleaved band (fusion protein)
Cleaved band

**Fig. S8. Recombination in reporter cells by VFIC system (adding EVs) evaluated by fluorescent microscope and the protein expression of VFIC related constructs.** (**A** and **B**) Representative fluorescence microscopy images showed the GFP expression in HeLa-TL and MSC-TL cells respectively after EV-mediated Cre delivery. Indicated doses of isolated EVs were incubated for the indicated time. Isolated EVs from VFIC system demonstrated dose and time dependent Cre delivery into recipient cells. Scale bar, 100 μm. (**C**) The expression of constructs related to VFIC system as measured by western blot analysis. Proteins from $5\times10^5$ EV-producing cells and $1\times10^{10}$ engineered vesicles were analyzed. WB: western blot. TSG101 was included as EV marker.

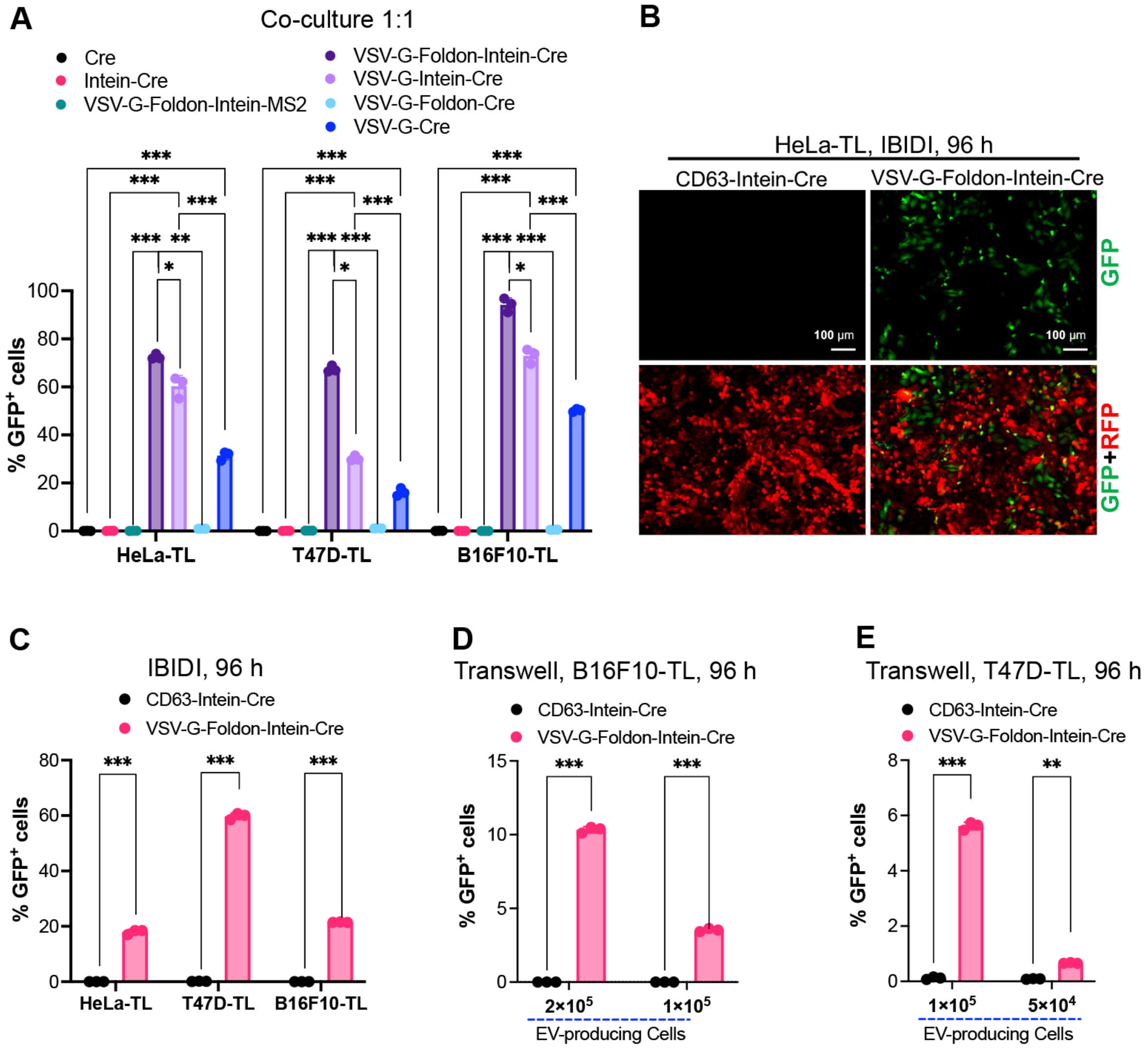


**Fig. S9. Flow cytometry analysis of recombination in reporter cells by EV-mediated Cre delivery using the VFIC system** (**A**) Flow cytometry analysis of a direct co-culture assay using various EV engineering strategies to assays Cre transfer after 24 hours. (**B**) Fluorescent microscopy demonstrating the Cre-mediated GFP expression after a IBIDI assay in HeLa-TL cells. Scale bar, 100 μm. (**C**) Percentage of GFP positive cells evaluated by IBIDI assay for 96 h. $4\times10^4$ EV producing cells and $4\times10^4$ reporter cells were seeded into the surrounding reservoirs and central reservoir respectively and flow cytometry was performed after 4 days. (**D** and **E**) Percentage of GFP positive cells in B16F10-TL and T47D-TL cells respectively evaluated by IBIDI assay for 96 h. Indicated numbers of EV-producing cells were added into the up chamber of the Transwell system and co-cultured with $5\times10^4$ B16F10-TL or $8\times10^4$ T47D-TL reporter cells in the lower chamber in 24- well plates. Two-way ANOVA multiple comparisons test was used for analysis of (A) and (C) to (E). Data are shown as means+SD, * p < 0.05; ** p < 0.01; *** p < 0.001.

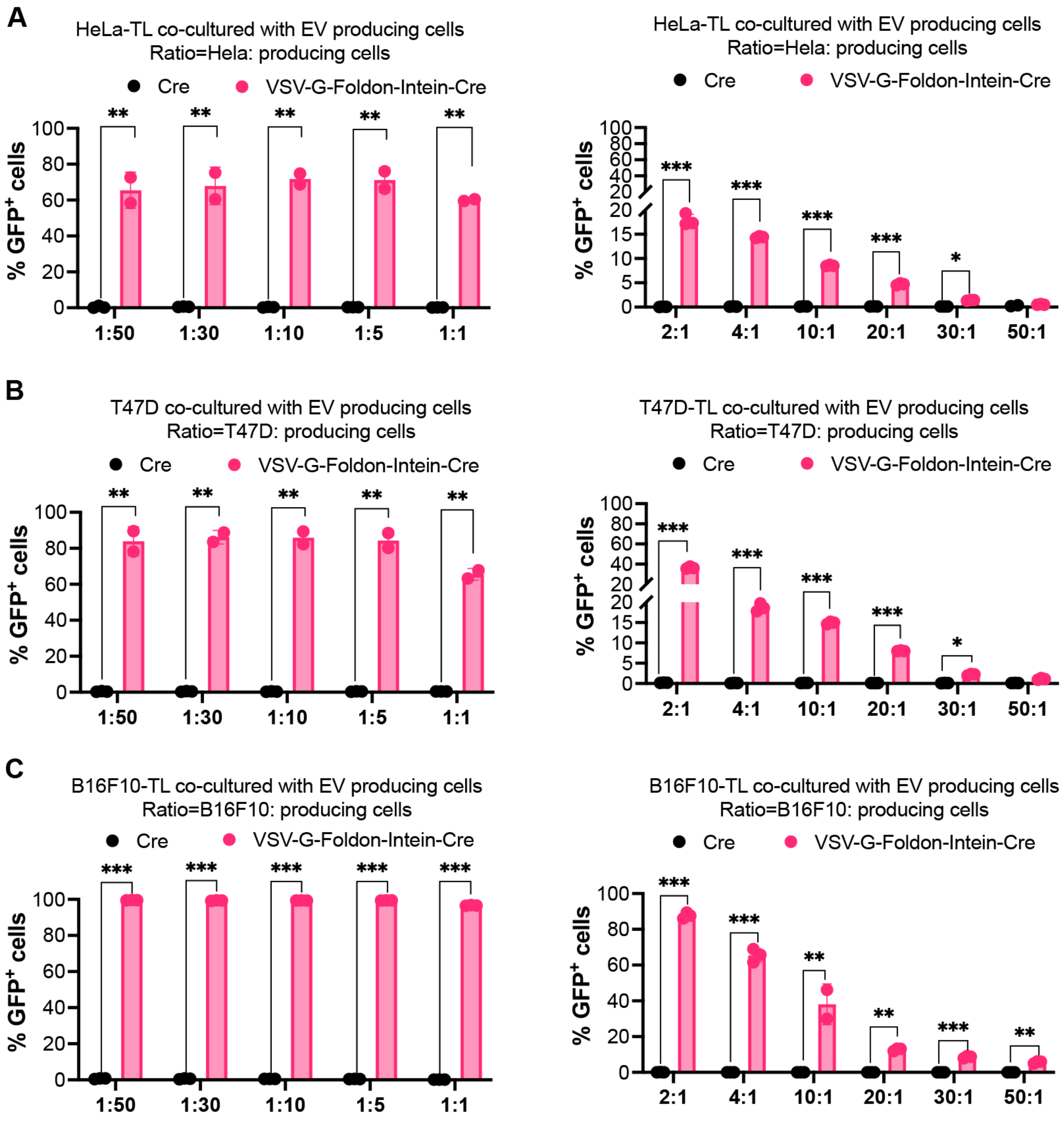


**Fig. S10. Efficient Cre transfer at varying cell ratios using the VFIC system** (Direct co-culture at varying ratios of EV-producing cells to reporter cells using **(A)** HeLa-TL, **(B)** T47D-TL, and **(C)** B61F10-TL cells. Two-way ANOVA multiple comparisons test was used for analysis of (A) to (C). Data are shown as means+SD, * p < 0.05; ** p < 0.01; *** p < 0.001.

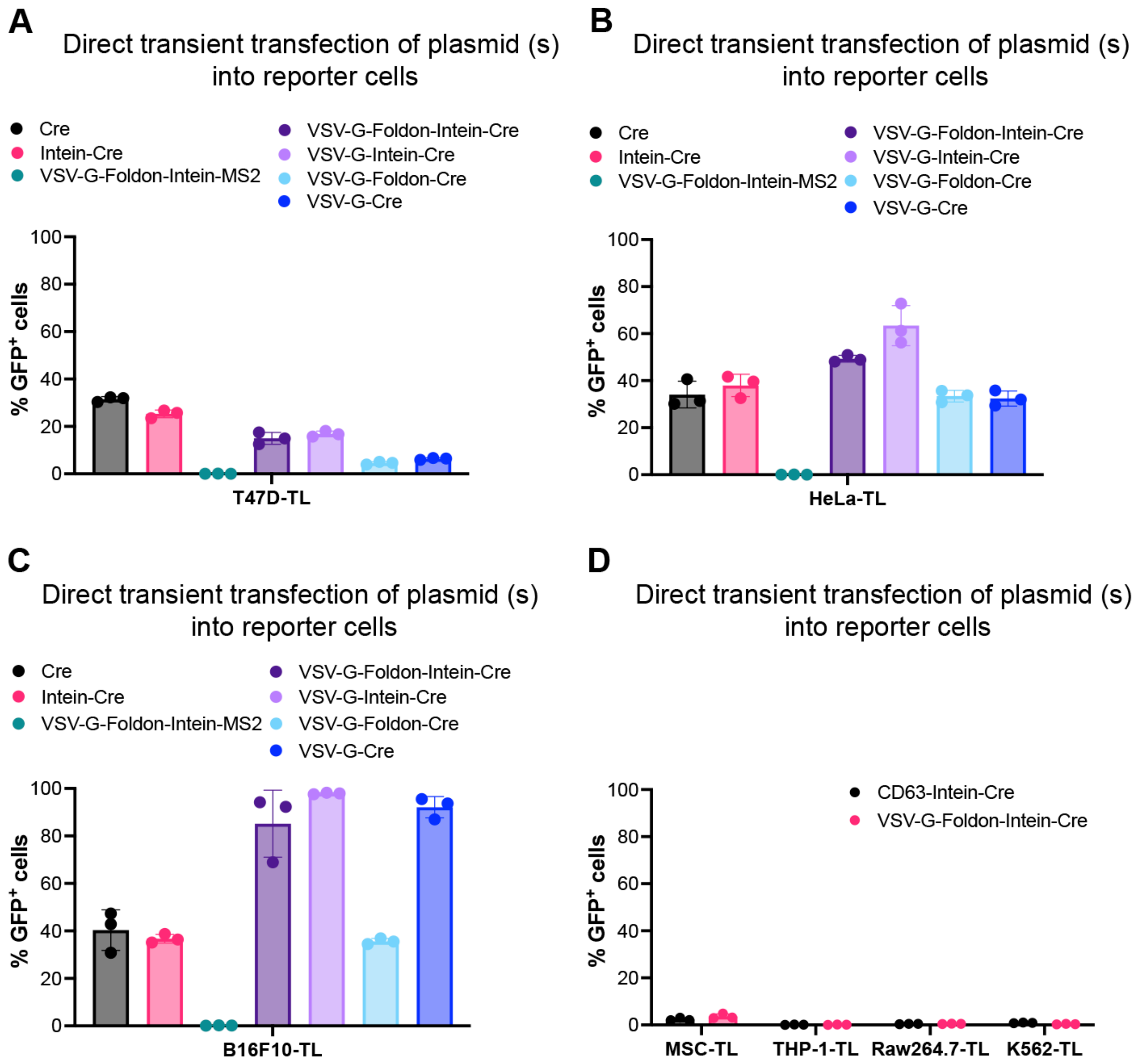


**Fig. S11. Cre-mediated recombination in reporter cells following transient transfection of indicated plasmids evaluated by flow cytometry.** (**A-D**) GFP positive cells measured by flow cytometry 48 h after transient transfection of the indicated plasmids into reporter cells.

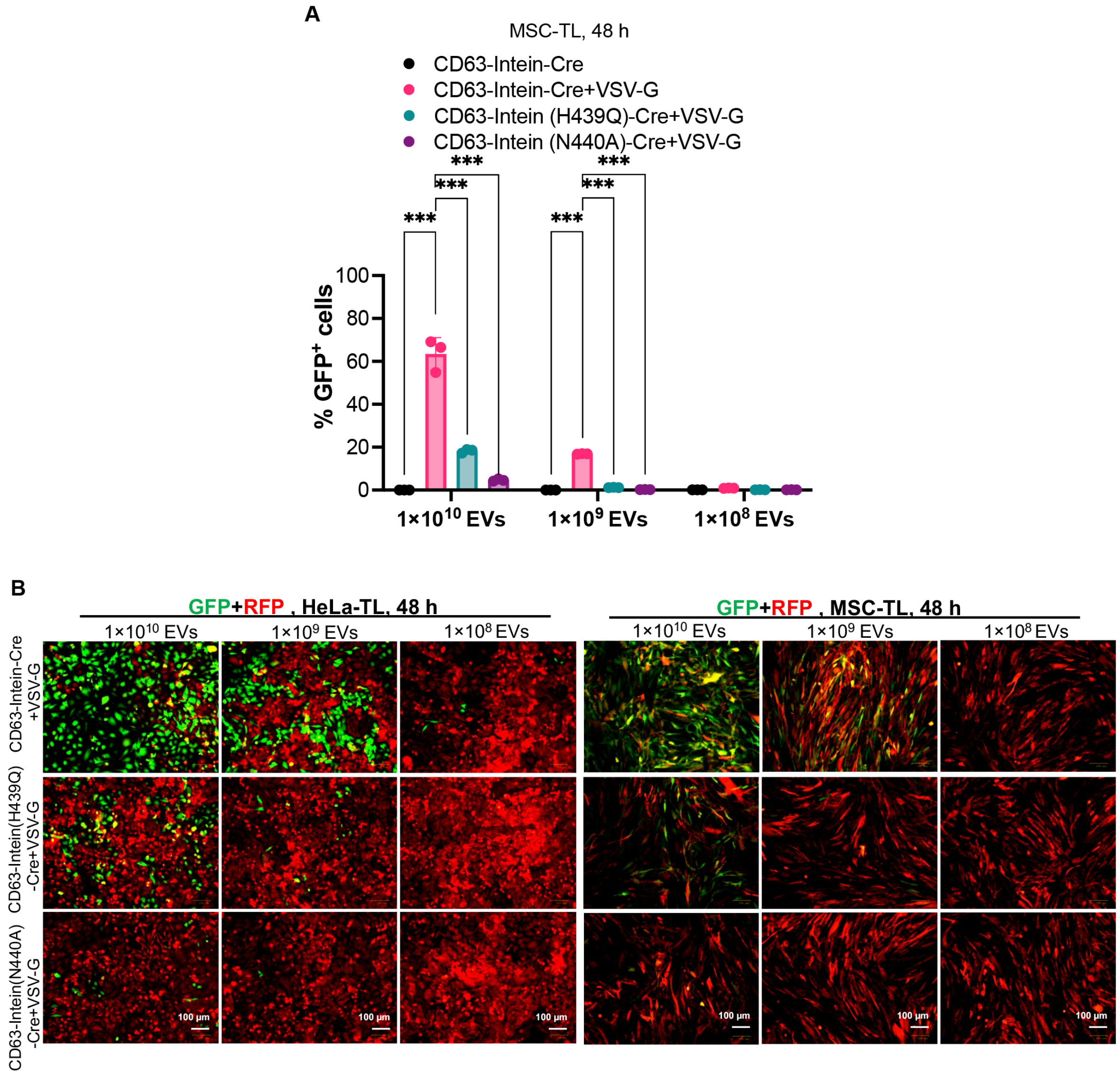


**Fig. S12. Intein variants decreased C-terminal cleavage for the VEDIC system evaluated by adding EVs directly. (A)** GFP positive cells measured by flow cytometry in MSC-TL cells. **(B)** Fluorescent images showing the GFP positive cells in Hela-TL and MSC-TL cells. Scale bar, 100 μm. Two-way ANOVA multiple comparisons test was used for analysis of (A). Data are shown as means+SD, *** $p < 0.001$.

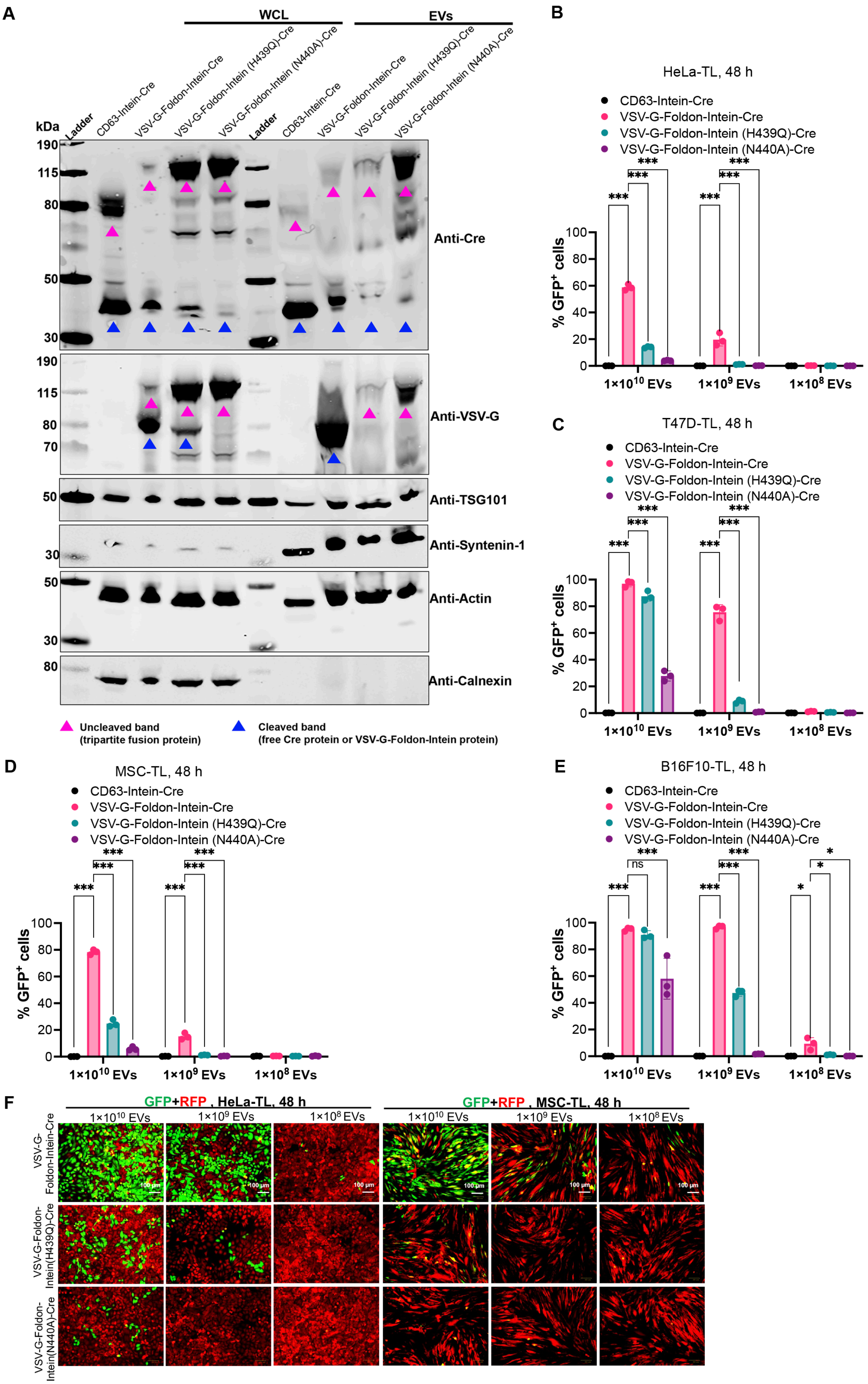

A
WCL
EVs
Ladder
CD63-Intein-Cre
VSV-G-Foldon-Intein-Cre
VSV-G-Foldon-Intein (H439Q)-Cre
VSV-G-Foldon-Intein (N440A)-Cre
kDa
Anti-Cre
Anti-VSV-G
Anti-TSG101
Anti-Syntenin-1
Anti-Actin
Anti-Calnexin
Uncleaved band (tripartite fusion protein)
Cleaved band (free Cre protein or VSV-G-Foldon-Intein protein)
B
HeLa-TL, 48 h
C
T47D-TL, 48 h
D
MSC-TL, 48 h
E
B16F10-TL, 48 h
% GFP+ cells
1×10^10 EVs
1×10^9 EVs
1×10^8 EVs
ns
F
GFP+RFP, HeLa-TL, 48 h
GFP+RFP, MSC-TL, 48 h
VSV-G-Foldon-Intein-Cre
VSV-G-Foldon-Intein(H439Q)-Cre
VSV-G-Foldon-Intein(N440A)-Cre
100 μm

**Fig. S13. Intein variants decreased C-terminal cleavage for the VFIC system evaluated by adding EVs directly.** (**A**) Protein expression of various engineered mutant intein constructs in whole cell lysates (WCL) and isolated EVs derived from HEK-293T cells evaluated by western blot analysis. Lysates from $5\times10^5$ EV-producing cells and $1\times10^{10}$ engineered vesicles were used for the assay. TSG101, syntenin-1 and β-actin were used as EV markers and Calnexin was used as a cellular organelle marker (endoplasmic reticulum) and should be absent for EV samples. (**B** to **E**) Recombination in reporter cells mediated by EVs derived from engineered cells using mutant inteins (H439Q and N440A) in VFIC system detected by flow cytometry. (**F**) Representative fluorescent images for the GFP positive cells after 48 hours' adding various mutant intein variants engineered EVs at different doses for VFIC system. Scale bar, 100 μm. Two-way ANOVA multiple comparisons test was used for analysis of (B) to (E). Data are shown as means+SD, * $p < 0.05$; *** $p < 0.001$.

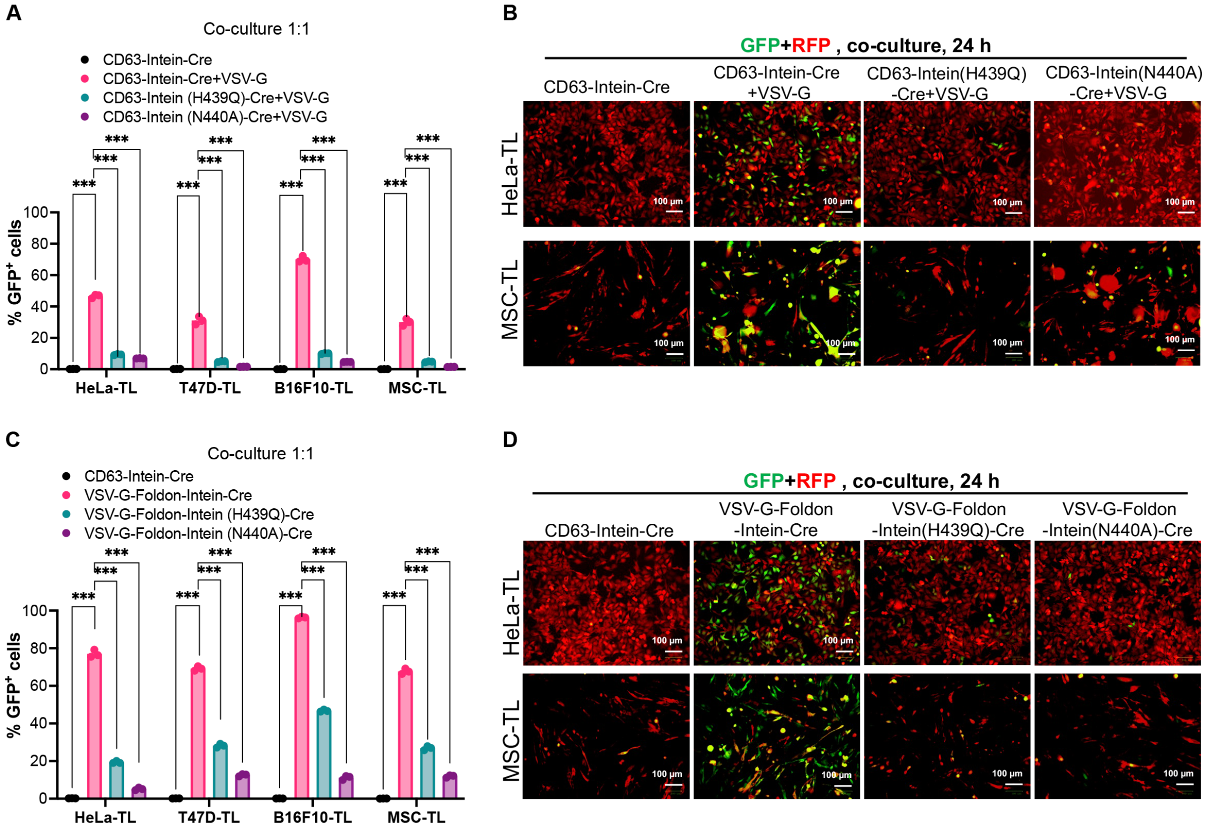


**Fig. S14. Intein variants decreased C-terminal cleavage for the VEDIC and VFIC systems evaluated by direct co-culture assay.** (**A**) Comparison of Cre transfer efficiency using different intein variants in a direct co-culture assay for VEDIC system. (**B**) GFP positive cells evaluated by fluorescent microscopic images using different intein variants in a direct co-culture assay for VEDIC system. Scale bar, 100 μm. (**C**) Flow cytometry analysis of a direct co-culture assay using intein variants to decrease Cre transfer for VFIC system. Co-culture was analyzed after 24 h incubation of the EV-producing cells and reporter cells. (**D**) Fluorescent microscopic images demonstrated GFP positive cells using intein variants in direct co-culture assay for VFIC system. Scale bar, 100 μm. Two-way ANOVA multiple comparisons test was used for analysis of (A) and (C). Data are shown as means+SD, *** $p < 0.001$.

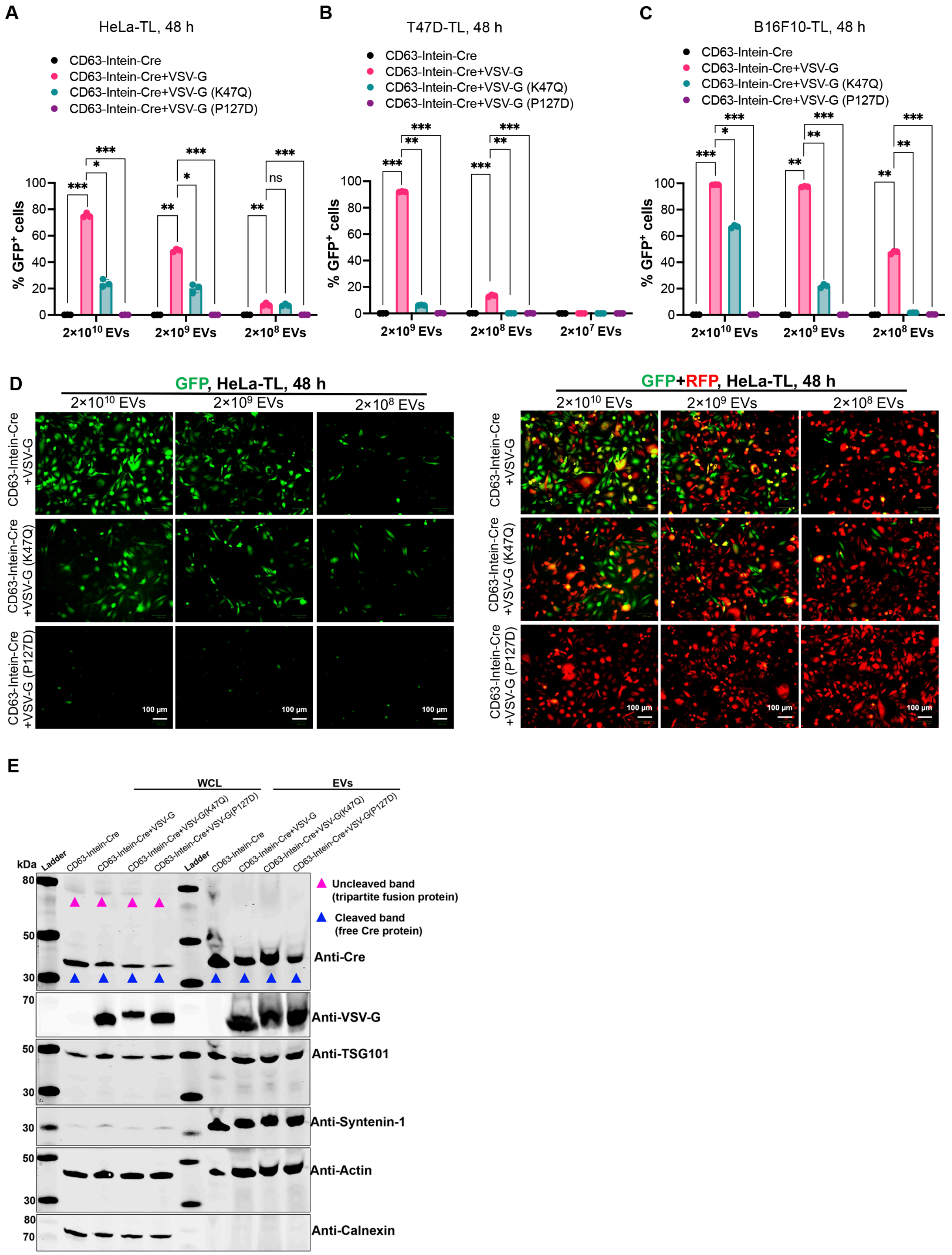


**Fig. S15. Evaluation of the function of VSV-G to enhance endosomal escape and receptor-mediated endocytosis for the entry of VSV-G engineered EVs using P127D and K47D mutants with the VEDIC system by adding EVs directly.** (**A** to **C**) Percentage of GFP positive HeLa-TL, T47D-TL and B16F10-TL cells after adding CD63-Intein-Cre EVs or wild type, P127D or K47Q VSV-G Foldon-Intein-Cre EVs, as evaluated by flow cytometry. (**D**) Representative images showing GFP positive cells in HeLa-TL cells after adding the indicated engineered EVs for 48 h. (**E**) Protein expression of mutated VSV-G-related constructs in both whole cell lysate (WCL) and isolated EVs evaluated by western blot analysis. Proteins from $5\times10^5$ EV-producing cells and $1\times10^{10}$ engineered vesicles were used for the assay. TSG101, syntenin-1 and β-actin were used as EV markers while Calnexin was used as a cellular organelle marker (endoplasmic reticulum). Two-way ANOVA multiple comparisons test was used for analysis of (A) to (C). Data are shown as means+SD, * $p < 0.05$; ** $p < 0.01$; *** $p < 0.001$.

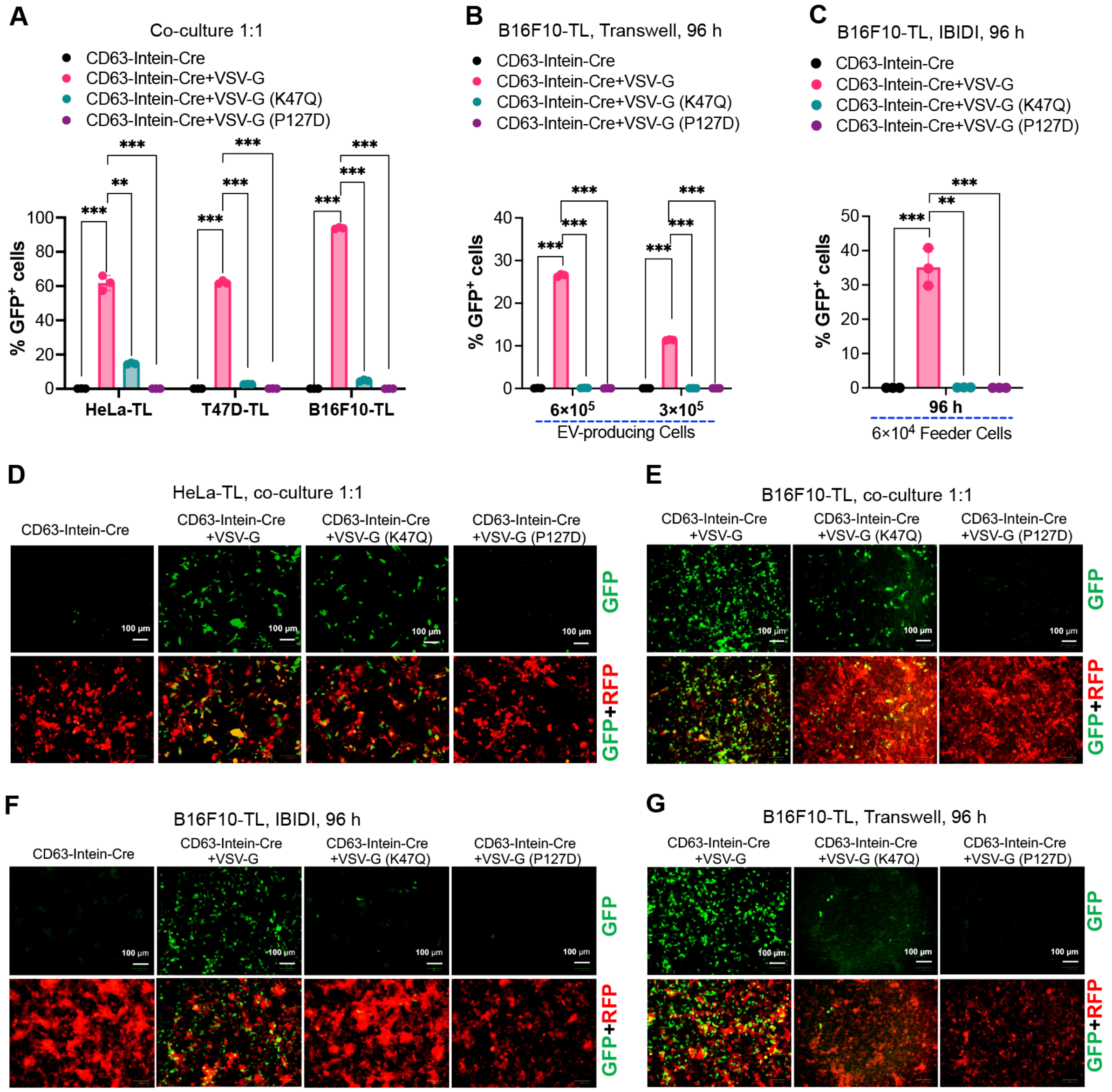


**Fig. S16. Evaluation of the function of VSV-G to enhance endosomal escape and receptor-mediated endocytosis for the entry of VSV-G engineered EVs using P127D and K47D mutants with the VEDIC system by co-culture assays.** (**A**) Direct co-culture assay to show the recombination efficiency of different engineered EVs. VSV-G (K47Q) and VSV-G (P127D) mutants were expressed in the EV-producing cells and resulting in abolished Cre delivery. (**B**) Percentage of recombined GFP positive cells after a Transwell assay with B16F10-TL cells. VSV-G (K47Q) and VSV-G (P127D) mutants were expressed in the EV producing cells. Indicated numbers of EV-producing cells and $5\times10^4$ B16F10-TL cells were used for the assay in 24-well plates. (**C**) Percentage of GFP positive cells evaluated by IBIDI assay in B16F10-TL cells. Indicated number of EV-producing cells and $4\times10^4$ reporter cells were used for this assay. (**D** and **E**) Representative fluorescent images showing the GFP positive cells from co-culture assay in HeLa-TL and B16F10-TL cells, respectively. Scale bar, 100 μm. (**F** and **G**) GFP positive cells as demonstrated by fluorescent microscopy from IBIDI and Transwell assay (24-well plates) respectively in B16F10-TL cells. Scale bar, 100 μm. Two-way ANOVA multiple comparisons test was used for analysis of (A) and (C). Data are shown as means+SD, * $p < 0.05$; ** $p < 0.01$; *** $p < 0.001$.

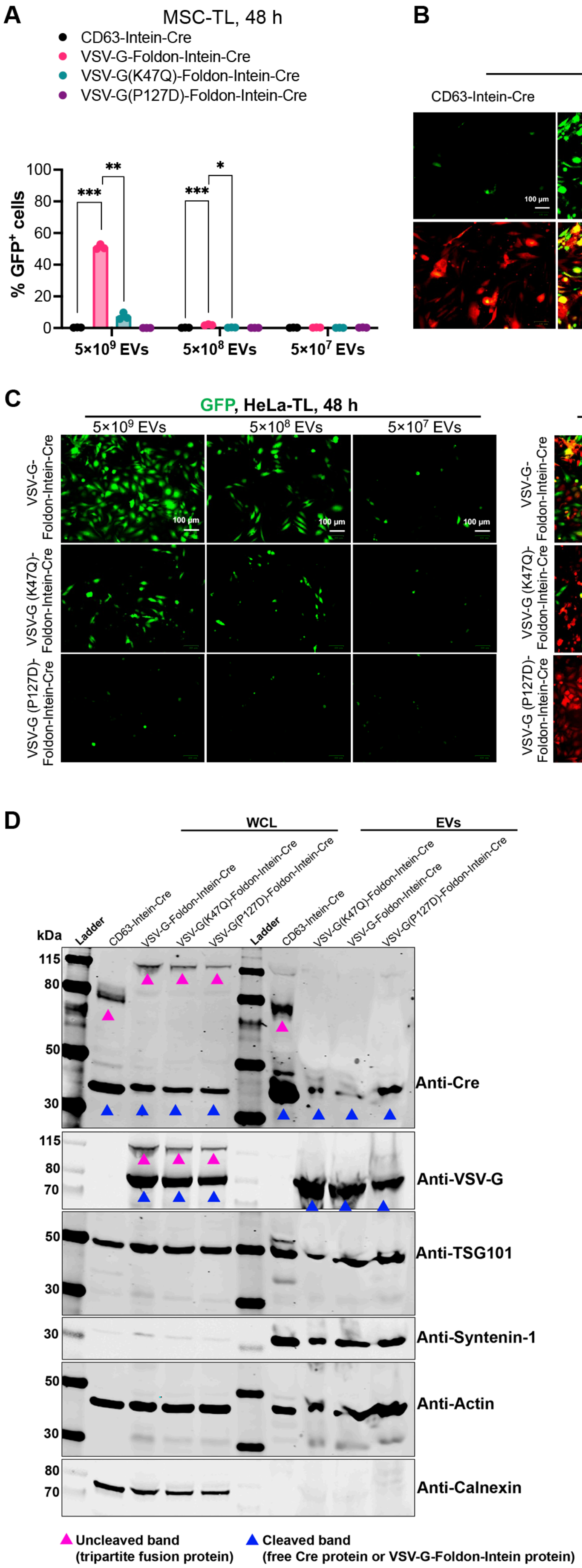
A
MSC-TL, 48 h
CD63-Intein-Cre
VSV-G-Foldon-Intein-Cre
VSV-G(K47Q)-Foldon-Intein-Cre
VSV-G(P127D)-Foldon-Intein-Cre
% GFP+ cells
5×10^9 EVs
5×10^8 EVs
5×10^7 EVs
B
MSC-TL, 48 h, 5×10^9 EVs
CD63-Intein-Cre
VSV-G-Foldon-Intein-Cre
VSV-G (K47Q)-Foldon-Intein-Cre
VSV-G (P127D)-Foldon-Intein-Cre
GFP
GFP+RFP
C
GFP, HeLa-TL, 48 h
GFP+RFP , HeLa-TL, 48 h
VSV-G-Foldon-Intein-Cre
VSV-G (K47Q)-Foldon-Intein-Cre
VSV-G (P127D)-Foldon-Intein-Cre
100 μm
D
WCL
EVs
Ladder
kDa
Anti-Cre
Anti-VSV-G
Anti-TSG101
Anti-Syntenin-1
Anti-Actin
Anti-Calnexin
Uncleaved band (tripartite fusion protein)
Cleaved band (free Cre protein or VSV-G-Foldon-Intein protein)

**Fig. S17. Evaluation of VSV-G function to enhance endosomal escape and receptor-mediated endocytosis for VSV-G engineered EVs on EV-mediated Cre delivery using P127D and K47D mutants for VFIC system by direct EV addition.** (**A**) Flow cytometry analysis after adding EVs from VSV-G engineered EVs for 48 h in MSC-TL cells. VSV-G (K47Q) and VSV-G (P127D) were directly fused with Foldon-Intein-Cre and were used to produce the engineered EVs. (**B**) Fluorescent microscopy images showing the GFP positive cells in MSC-TL cells after adding the indicated dose of EVs for 48 h. Scale bar, 100 µm. (**C**) Fluorescence microscopy images demonstrate GFP positive cells after adding the indicated doses of EVs for 48 h to HeLa-TL cells. Scale bar, 100 µm. (**D**) Protein expression of directly fused mutated VSV-G-related constructs both in whole cell lysates (WCL) of and isolated EVs evaluated by western blot analysis. Proteins from $5\times10^5$ EV producing cells and $1\times10^{10}$ engineered vesicles were used for the assay. TSG101, syntenin-1 and β-actin were used as EV markers and Calnexin was used as cellular organelle marker (endoplasmic reticulum) Two-way ANOVA multiple comparisons test was used for analysis of (A). Data are shown as means+SD, * $p < 0.05$; ** $p < 0.01$; *** $p < 0.001$.

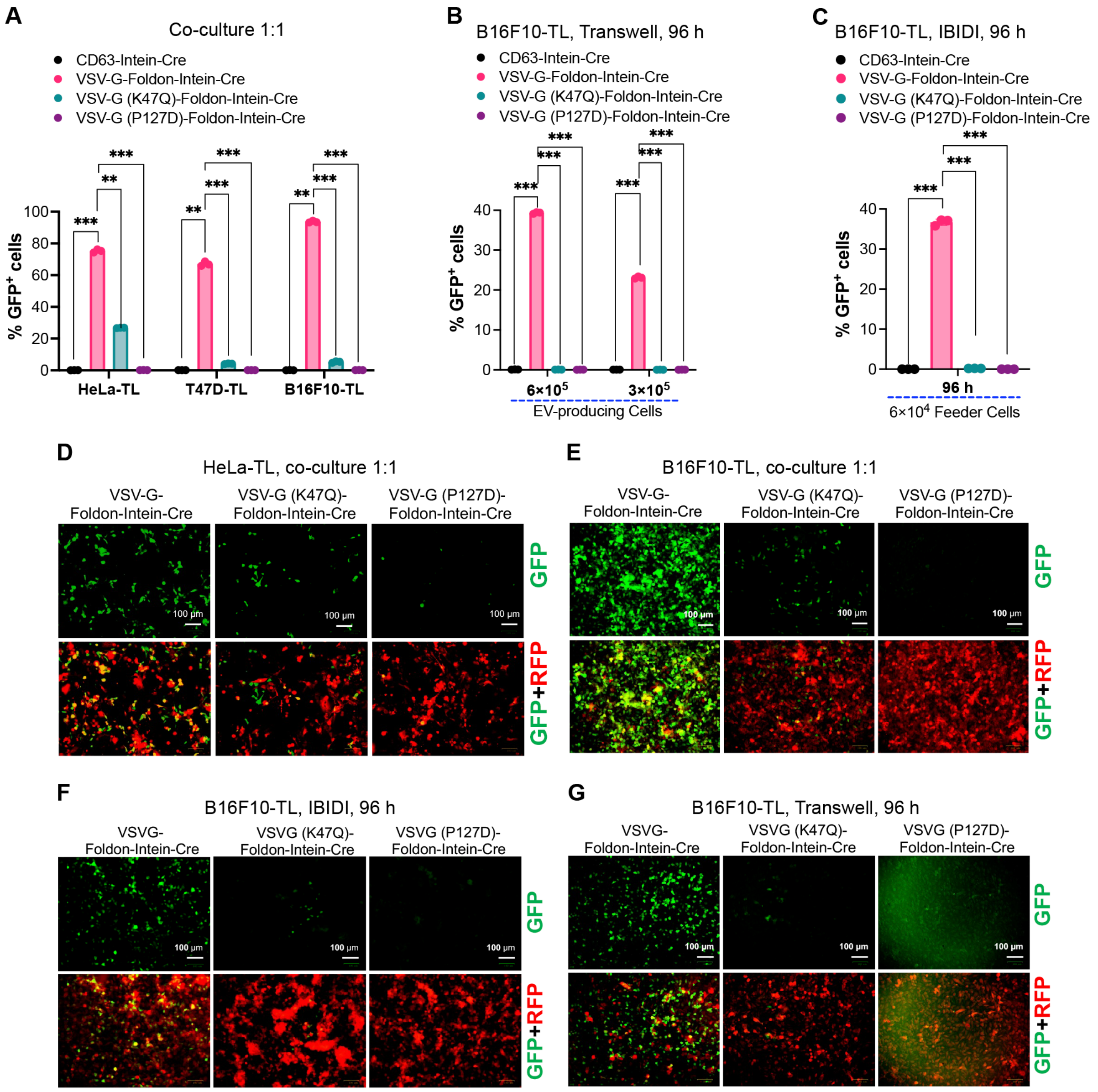

**Fig. S18. Evaluation of VSV-G function to enhance endosomal escape and receptor-mediated endocytosis for VSV-G engineered EVs on EV-mediated Cre delivery using P127D and K47D mutants for VFIC system in co-culture assays.** (**A**) Direct co-culture assay demonstrating the percentage of GFP positive cells in HeLa-TL, T47D-TL and B16F10-TL cells after 24 h. P127D and K47D mutants were included to assess the role of VSV-G for endosomal escape and receptor-mediated uptake of engineered EVs. (**B**) Percentage of GFP positive cells in B16F10-TL cells evaluated after a Transwell co-culture assay after 96 h. The indicated numbers of EV producing cells were seeded in the upper chamber and the reporter cells were added to the lower chamber in a 24-well plates. (**C**) Percentage of GFP positive cells in B16F10-TL cells evaluated by IBIDI assay. $6\times10^4$ EV-producing HEK293T cells and $4\times10^4$ B16F10-TL reporter cells were seeded into the surrounding reservoirs and central reservoir respectively for 4 days. (**D** and **E**) Representative images demonstrated GFP positive cells from a direct co-culture assay using HeLa-TL and B16F10-TL cells. Scale bar, 100 µm. (**F**) GFP positive cells in B16F10-TL cells as shown by fluorescent microscopy from IBIDI assay. Scale bar, 100 µm. (**G**) GFP positive cells in B16F10-TL cells as shown by fluorescence microscopy after a Transwell assay. Scale bar, 100 µm. Two-way ANOVA multiple comparisons test was used for analysis of (A) to (C). Data are shown as means+SD, ** $p < 0.01$; *** $p < 0.001$.

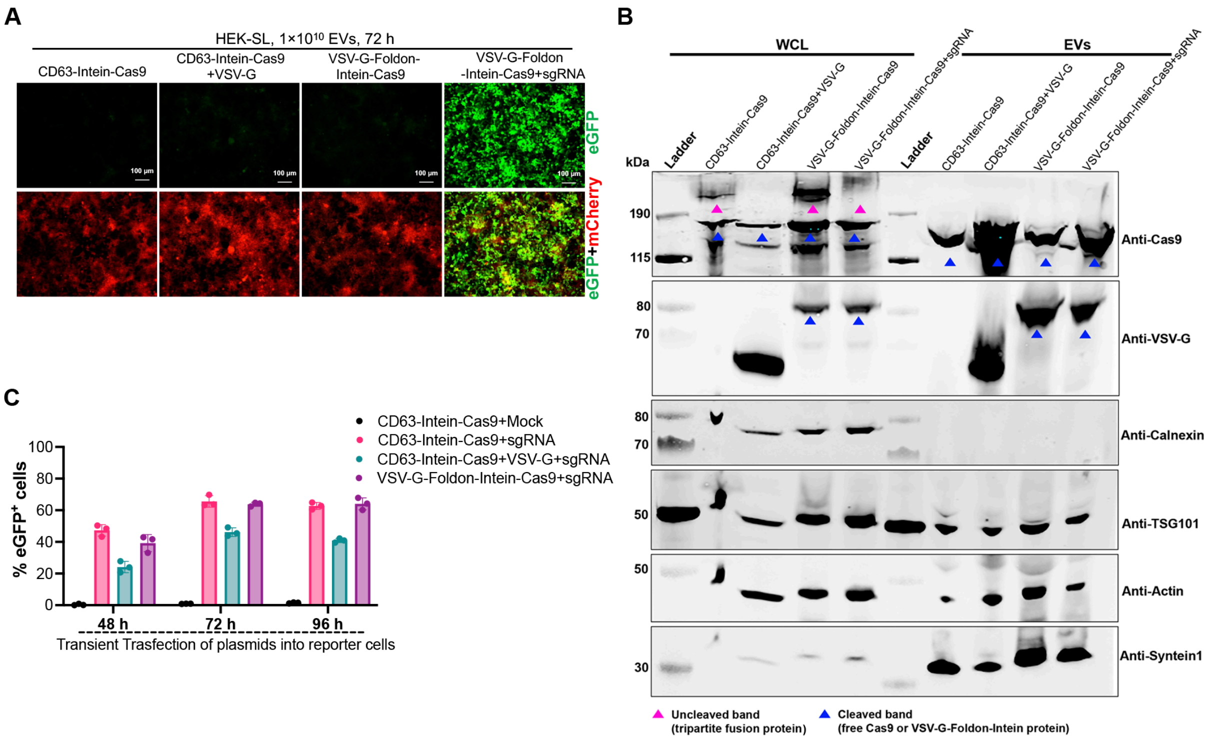


**Fig. S19. Characterization of Cas9-mediated genome editing in reporter cells by delivery of the engineered EVs** (**A**) Fluorescence microscopy images demonstrating functional Cas9/sgRNA RNPs delivery, as indicated by GFP positive signals, in HEK293T stoplight (HEK-SL) reporter cells after addition of indicated doses of EVs for 72 h. Scale bar, 100 µm. (**B**) Protein expression of Cas9 related constructs in both whole cell lysate (WCL) and EVs evaluated by western blot analysis. Proteins from $5\times10^5$ EV-producing cells and $1\times10^{10}$ engineered vesicles were used for the assay. TSG101, syntenin-1 and β-actin were used as EV markers while Calnexin was used as cellular organelle marker (endoplasmic reticulum). (**C**) Genome editing efficiency in HEK-SL reporter cells after transient transfection of the indicated constructs for indicated time.

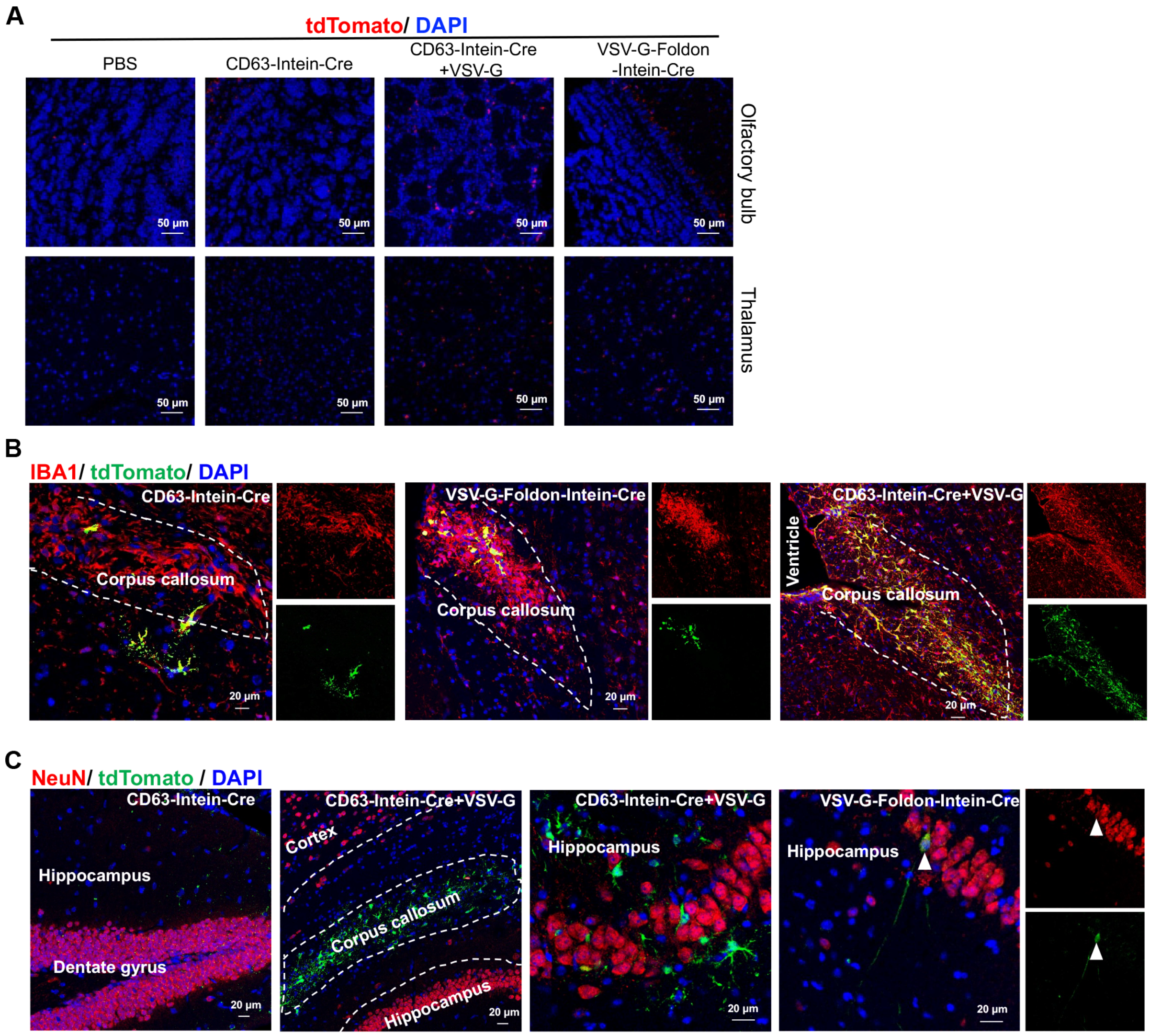


**Fig. S20. Cre mediated in vivo delivery in LoxP-Cre R26-LSL-tdTomato reporter mice by using VEDIC and VFIC systems after ICV injection.** (**A**) IHC staining of the olfactory bulb and thalamus one week after ICV injection of the engineered EVs. Scale bar, 50μm. (**B**) Co-staining of microglia marker IBA1 with tdTomato in corpus callosum (highlighted by dashed lines) after ICV injection of engineered EVs. Scale bar, 20 μm. (**C**) Co-staining of neuron cell marker NeuN with tdTomato in different regions of brain after ICV injection of different engineered EVs. Scale bar, 20 μm. White arrow indicates sporadic cells with co-localization of NeuN and RFP. n=3 mice per group, representative images.

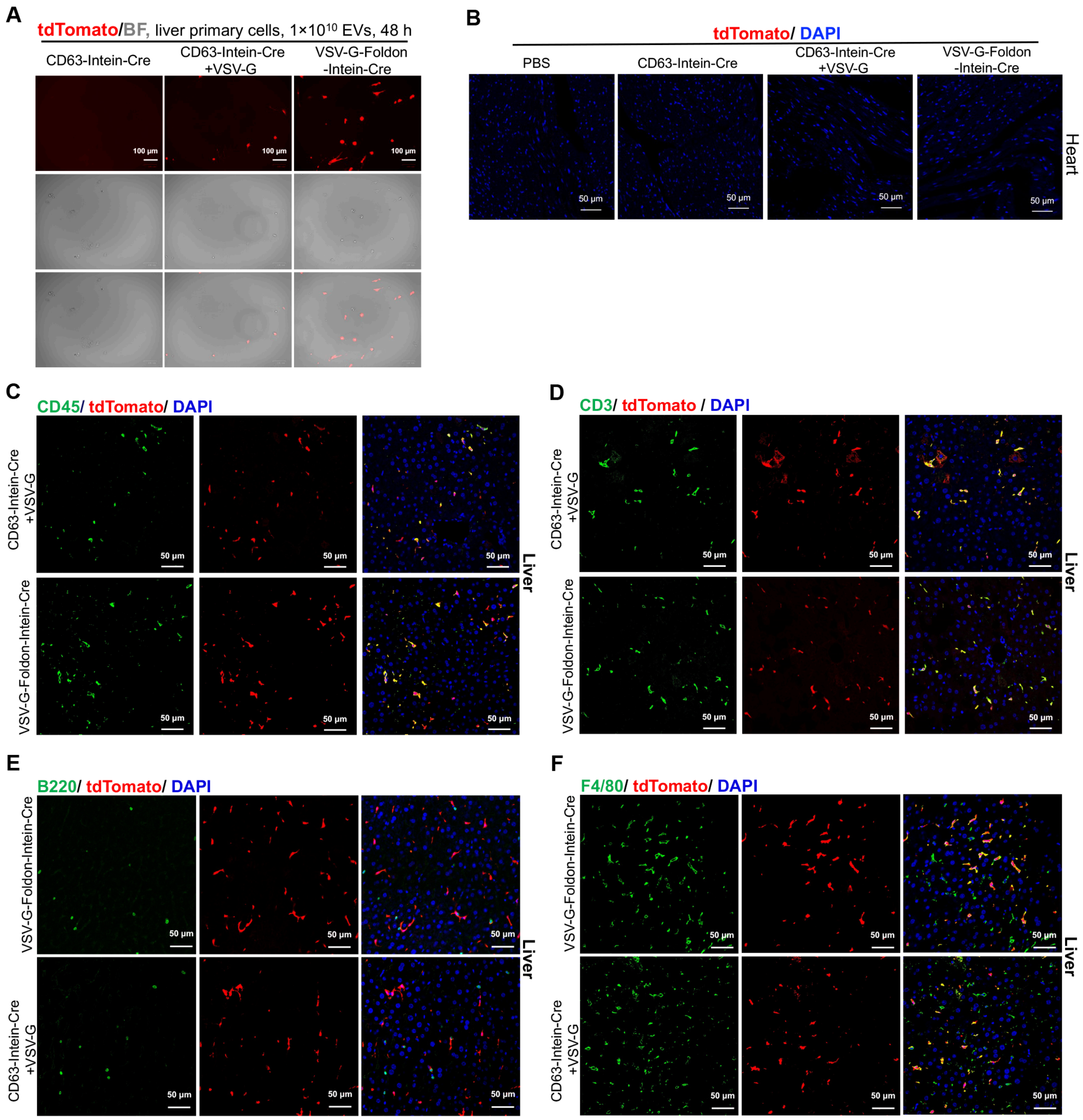


**Fig. S21. Cre mediated delivery in LoxP-Cre R26-LSL-tdTomato reporter mice using VEDIC and VFIC systems after IP injection.** (**A**) Representative fluorescent microscopic images demonstrated tdTomato positive liver primary cells from R26-LSL-tdTomato reporter mice after adding VEDIC and VFIC EVs for 2 days. Scale bar, 100 µm. (**B**) IHC staining of the heart, one week after IP injection of the engineered EVs. Scale bar, 50 µm. (**C**) Co-staining of general leukocyte marker CD45 with tdTomato expression in liver after IP injection of engineered EVs. Scale bar, 50 µm. (**D**) C-localization of T cell marker CD3 with tdTomato in liver one week after IP injection of engineered EVs as detected by IHC co-staining for. Scale bar, 50 µm. (**E**) Co-localization of B cell marker B220 with tdTomato in liver one week after IP injection of different engineered EVs. Scale bar, 50 µm. (**F**) Co-expression of macrophage marker F4/80 with tdTomato in liver evaluated by IHC co-staining one week after injecting engineered EVs. Scale bar, 50 µm. n=3 mice per group, representative images.

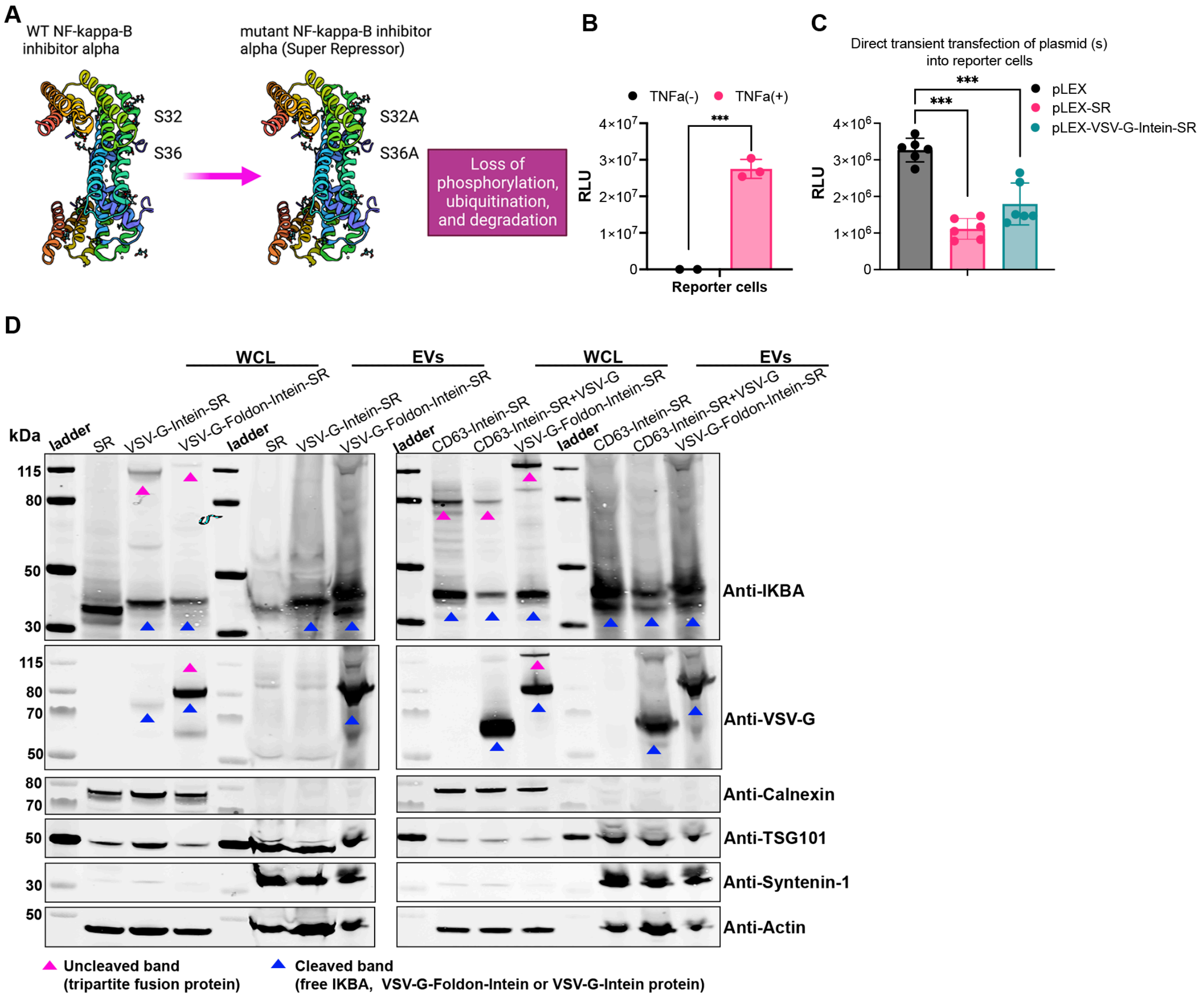


**Fig. S22. Design of super repressor (SR) inhibitor of NF-κB constructs and their expression and function in vitro.** (**A**) The mutations and properties of the super repressor inhibitor of NF-κB. (**B**) Activation of luciferase expression 6 h after TNF-α stimulation (10 ng/ml) in HEK-Blue-NF-κB reporter cells. (**C**) Significant decrease of the luciferase signals from lysate of HEK-Blue-NF-κB reporter cells after transient transfection of SR-related constructs. 48 h after transfection cells were stimulated with TNF-α for 6 h prior to luciferase detection. (**D**) Protein expression of SR-related constructs in both whole cell lysate (WCL) and isolated EVs evaluated by western blot analysis. Proteins from $5\times10^5$ EV-producing cells and $1\times10^{10}$ engineered vesicles were used for the assay. TSG101, syntenin-1 and β-actin were used as EV markers while Calnexin was used as cellular organelle marker (endoplasmic reticulum). Two-tailed T-test was used for the analysis of (B); One-way ANOVA multiple comparisons test was used for analysis of (C). Data are shown as means+SD, *** p < 0.001.
